# PantheonOS: An Evolvable Multi-Agent Framework for Automatic Genomics Discovery

**DOI:** 10.64898/2026.02.26.707870

**Authors:** Weize Xu, Erwin Poussi, Quan Zhong, Zehua Zeng, Christopher Zou, Xuehai Wang, Yifan Lu, Miao Cui, Daiji Okamura, Cinlong Huang, Jiayuan Ding, Zhe Zhao, Yuheng Yang, Xinhai Pan, Varshini Vijay, Naoki Konno, Nianping Liu, Lei Li, X. Rosa Ma, Stephanie D. Conley, Colin Kern, William R. Goodyer, Bogdan Bintu, Quan Zhu, Neil C. Chi, Jiang He, Lorenz Rognoni, Xiuwei Zhang, Jun Wu, David Ellison, Marlene Rabinovitch, Jesse M. Engreitz, Xiaojie Qiu

**Affiliations:** Department of Genetics, Stanford University, Stanford, CA, USA; Basic Sciences and Engineering Initiative, Betty Irene Moore Children’s Heart Center, Lucile Packard Children’s Hospital, Stanford, CA, USA; Department of Computer Science, Stanford University, Stanford, CA, USA; Stanford Cardiovascular Institute, Stanford University, Stanford, CA, USA; Department of Physiology and Pharmacology, Karolinska Institutet, Stockholm, Sweden; Hippocratic AI, Palo Alto, CA, USA; Department of Molecular Biology, University of Texas Southwestern Medical Center, Dallas, TX 75390, USA; Department of Advanced Bioscience, Faculty of Agriculture, Kindai University, Nara, Japan; School of Computational Science and Engineering, Georgia Institute of Technology, Atlanta GA 30332, USA; Department of Computer Science, University of California Irvine, Irvine, CA, USA; Novo Nordisk Foundation Center for Genomic Mechanisms of Disease, Broad Institute, Cambridge, MA, USA; Department of Cellular and Molecular Medicine, University of California San Diego, La Jolla, CA, USA; Department of Bioengineering, University of California San Diego, School of Medicine, La Jolla, CA, USA; Center for Epigenomics, Department of Cellular and Molecular Medicine, University of California San Diego, La Jolla, CA, USA; Department of Medicine, Division of Cardiology, University of California San Diego, La Jolla, CA, USA; Institute for Genomic Medicine, University of California San Diego, La Jolla, CA, USA; Institute of Engineering in Medicine, University of California San Diego, La Jolla, CA, USA; Vizgen, Cambridge, MA, USA; Hamon Center for Regenerative Science and Medicine, University of Texas Southwestern Medical Center, Dallas, TX 75390, USA; Cecil H. and Ida Green Center for Reproductive Biology Sciences, University of Texas Southwestern Medical Center, Dallas, TX 75390, USA; Infrastructure Solution Group, Lenovo; Department of Pediatrics, Stanford University School of Medicine, CCSR-1215A, 269 Campus Drive, Stanford, CA, 94305-5162, USA; Vera Moulton Wall Center for Pulmonary Vascular Diseases, Stanford University School of Medicine, Stanford, CA, 94305, USA; Maternal and Child Health Research Institute, Stanford University, Stanford, CA, USA; Shandong University, Inspur Jointly Launch AI Institute; Canchen Technology, Shenzhen 518000, China; Independent Researcher, Jiangxi, China

## Abstract

The convergence of large language model-powered autonomous agent systems and single-cell biology promises a paradigm shift in biomedical discovery. However, existing biological agent systems, building upon single-agent architectures, are narrowly specialized or overly general, limiting applications to routine analyses. We introduce PantheonOS (https://PantheonOS.stanford.edu), an evolvable, privacy-preserving multi-agent framework designed to reconcile generality with domain specificity. Critically, PantheonOS enables agentic code evolution, allowing evolving state-of-the-art batch correction and our reinforcement-learning augmented gene panel selection algorithms to achieve super-human performance. PantheonOS drives biological discoveries across systems: uncovering asymmetric paracrine *Cer1–Nodal* inhibition in proximal–distal axis formation of novel early mouse embryo 3D data; integrating human fetal heart multi-omics with whole-heart data to reveal molecular programs underpin heart diseases; and adaptively selecting virtual cell models to predict cardiac regulatory and perturbation effects. Together, PantheonOS points towards a future where scientific discoveries are increasingly driven by self-evolving AI systems across biology and beyond.

**Website:** https://pantheonos.stanford.edu

**Ecosystem:** https://github.com/aristoteleo

**Summary:** Large language model–powered agent systems are driving a paradigm shift in scientific discovery by automating, scaling, and accelerating data analysis. This transformation is particularly profound in biology, where the rapid expansion of single-cell and spatial genomics has effectively reshaped the field into a data-intensive science. However, existing biological agent systems are typically constrained to single-agent designs, are narrowly specialized, or are overly general without sufficient domain expertise, limiting their applicability to routine or shallow analyses. Here, we introduce **PantheonOS** (https://pantheonos.stanford.edu), an evolvable, privacy-preserving, and general-purpose multi-agent framework designed to reconcile generality with deep domain specificity. PantheonOS provides an abstract, extensible architecture that enables customized agent composition and supports end-to-end single-cell and multi-omics analysis, spanning reinforcement-learning–augmented gene panel design, raw FASTQ processing, multimodal data integration, and three-dimensional spatial genomics reconstruction. Central to this framework, **Pantheon-Evolve** enables agentic code evolution, allowing the system to autonomously improve state-of-the-art batch-correction algorithms and new reinforcement-learning based gene panel design algorithms, achieving performance beyond manually designed baselines. We demonstrate the power of PantheonOS across multiple biological domains. In early mouse embryogenesis, PantheonOS automatically reconstructs three-dimensional spatial gene expression landscapes and resolves asymmetric *Cer1* expression and paracrine *Cer1–Nodal* inhibition, revealing a robust proximal–distal axis at embryonic day six (E6.0). In human development, PantheonOS integrates fetal heart single-cell multi-omics with whole-heart 3D MERFISH+ data at post-conception week 12, uncovering spatially resolved molecular programs underlying heart disease ontogeny. Finally, an intelligent model-routing mechanism enables PantheonOS to adaptively select optimal virtual cell models across heterogeneous tasks, revealing minimal regulatory networks of cardiogenesis and predicting spatially resolved perturbation effects in the developing heart. Together, PantheonOS establishes a foundation for fully automated, evolvable, and domain-aware agentic analysis in genomics, and points toward a future in which scientific discovery is increasingly driven by self-improving AI systems across biology and beyond.

## Introduction

Recent advances in large language models (LLM) and LLM-powered agentic systems have substantially expanded capabilities in reasoning, planning, code generation, and tool execution, facilitating automatic, iterative exploration of complex data at a scale and pace that are difficult to sustain through purely manual analysis^1^. These advances are particularly relevant to biology, where progress in single-cell and spatial genomics has generated unprecedented large-scale, high-resolution, and highly heterogeneous datasets that increasingly challenge conventional analysis workflows^2^. Despite these progresses, however, the practical impact of LLMs and agentic systems on real-world data science, especially in biological applications, remains limited. Most existing biological agent frameworks such as STAgent^3^, SpatialAgent^4^, CellVoyager^5^ and Biomni^6^ rely on single-agent designs, lacking principled multi-agent architectures that enable agent team collaboration and collective problem solving. As a result, these systems either adopt a narrow task focus or remain overly general, without the deep domain expertise required for nontrivial biological analyses. With the exception of Biomni, there is no mature, end-to-end platform that biologists can directly adopt in practice. Moreover, current biological agent systems are predominantly web graphic user interface or GUI-based, constraining flexibility and integration into existing computational workflows. They typically lack mechanisms for community-driven extension and are not designed as distributed systems capable of decoupling agents from execution environments for privacy protection of sensitive genomic datasets. Lastly, they do not support automatic or recursive code evolution and optimization. Consequently, the analyses they enable are largely limited to standard tasks, such as cell type annotation or trajectory inference, rather than driving complex investigations that create truly innovative algorithms and yield genuinely novel biological insights.

We introduce PantheonOS as a first-generation, evolvable, distributed, and general-purpose AI agent platform that establishes a new paradigm for data-driven scientific discovery. PantheonOS is built on an abstract, domain-agnostic foundation while simultaneously deploying a purpose-built multi-agent team for genomics research, decisively resolving the dilemma between generality and domain-specific intelligence. The platform is engineered for real-world scientific practice, offering a well-designed GUI-based integrative platform but also command-line interface for power users, native support for team-based collaboration through Slack, and seamless integration with Jupyter notebooks to embed agentic intelligence directly into existing analytical workflows. To meet the stringent requirements of biomedical research, PantheonOS adopts a distributed architecture design to meet privacy protection needs, and introduces the Pantheon Store as a mechanism for community-driven extension. The system enables truly end-to-end analysis pipelines, spanning reinforcement learning–based gene panel design, raw FASTQ processing, and downstream single-cell and spatial transcriptomics analyses, and is validated on novel large-scale 3D spatial transcriptomics and single-cell multi-omics datasets. Critically, we develop Pantheon-Evolve, an automated framework for recursive algorithmic evolution that surpasses human-designed baselines, achieving super-human performance on evolved versions of Harmony^7^, Scanorama^8^, and BBKNN^9^. This evolution framework generalizes to newly developed reinforcement learning algorithms, underscoring its capacity to evolve essentially all existing and emerging computational methods and to accelerate the pace of algorithmic innovation. Leveraging PantheonOS as a virtual research assistant, we generate a novel OpenST-based 3D spatial transcriptomics dataset of early mouse embryogenesis at E6.0, revealing spatially segregated *Cer1* and *Nodal* domains within the epiblast and demonstrating active paracrine inhibition of *Nodal* signaling by *Cer1*. Furthermore, PantheonOS enables large-scale integration of single-cell multi-omics data from the human fetal heart with whole-organ 3D MERFISH+ maps at post-conception week 12, uncovering spatial gene expression programs and intercellular communication mechanisms that illuminate cardiac disease ontogeny. Finally, we establish an intelligent single-cell foundation model router that unifies major single cell foundation models or virtual cell models^10^ and demonstrates the recovery of core cardiogenic regulatory networks directly within their native three-dimensional spatial context.

Together, these results establish PantheonOS as an evolvable operating system for biological data science. By combining distributed multi-agent design, community extensibility, privacy-aware infrastructure, and algorithmic self-evolution, PantheonOS redefines how complex biological analyses can be conducted. PantheonOS is publicly deployed at **PantheonOS.stanford.edu**, where we invite the community to contribute, extend, and collectively reimagine the future of automated biological discovery.

## Results

### PantheonOS: an evolvable distributed general-purpose multi-agent framework, instantiated on automated single-cell and spatial genomics analysis

The rapid growth of single-cell and spatial omics data has created a data deluge in genomics, effectively transforming the field into a large-scale data science discipline. More broadly, across data science domains, the increasing scale, complexity, and privacy sensitivity of modern datasets demand automated, reproducible, and sophisticated systems capable of accelerating expert-level analysis and discovery. Recent general-purpose AI agent frameworks (e.g. Langchain^11^, AutoGen^12^, CrewAI^13^) and platforms (OpenAI frontier^14^) have begun to address this need by orchestrating tools through natural-language interfaces. However, these systems remain fundamentally static: they primarily invoke predefined tools and lack the ability to evolve the underlying algorithms they depend on, let alone recursively improve themselves to surpass existing state-of-the-art methods. In addition, data privacy is rarely treated as a first-class design principle. Within biology, existing multi-agent systems, either emphasize generality at the expense of deep domain expertise, limiting their effectiveness for advanced genomics tasks^6^, or adopt highly specialized designs that are difficult to extend beyond narrow applications^3–5,15^. To address these limitations, we introduce PantheonOS, an abstract, evolvable, and distributed multi-agent framework.

PantheonOS employs a four-layer pyramid architecture (**Fig. 1A**) that builds from the LLM layer, to the Agent, Interface, and finally to the Application layer, embraces flexible user interfaces (**Fig. 1B)** and establishes an evolvable distributed multi-agent system (**Fig. 1C**), reconciling the dilemma with both generality as a framework and specificity for domain applications, enabling flexible and privacy protected data analyses that have the potential to achieve super-human performance. The LLM Layer 1 includes a unified LLM interface supporting 100+ LLMs with automatic retry and fallback. It also enables distributed communication via NATS (https://nats.io/), enabling flexible cross-device deployment (**Fig. S1B**). For the Agent Layer 2, it provides the runtime execution model that enables agents to coordinate through a unified agentic loop, structured inter-agent transfer, and a formal Modal Task Protocol (MTP) that governs task states and artifacts (**Methods, Fig. S1A**). Agentic Context Engineering (ACE) further supports long-horizon, privacy-aware reasoning by dynamically managing and compressing agent context during execution, allowing PantheonOS to function as a self-organizing system rather than a fixed analysis pipeline (**Methods, Fig. S1A**). Through experimentation, we found that the Pantheon agent outperforms Biomni^6^ across multiple evaluation metrics (**Fig. S1C, D**). Furthermore, ablation studies demonstrate that the MTP design within its operational mechanism significantly enhances performance (**Fig. S1E**). This layer also ships with a series of pre-built toolsets (code execution, notebook, web search, foundation model router, etc.), and specialized agents (Data Analyzer, Biology, Coding, Evolution, Browser, Reporter, etc.). It also implements multiple team communication patterns (PantheonTeam, SwarmTeam, MoATeam^16^), and an extensible skill system encoding domain expertise as markdown templates with structured workflows (**Fig. 1C**). Critically, the Pantheon-Evolve module enables agents to iteratively improve external and internal algorithms, packages or skills through agent-guided evolution to achieve super-human performance. Importantly, the external MCP ecosystem interfaces can connect to many existing MCPs including BioMCP (https://biomcp.org/), BioContextAI^17^, Context7 (https://context7.com/) and others. Interface Layer 3 has a very flexible design to account for different user needs, including CLI, Jupyter notebook, web GUI and Slack chatbot (**Fig. 1B**). The CLI interface is suitable for power users familiar with the Linux ecosystem, allowing them to start it directly in the terminal and access existing scripts and data in the system. It is also particularly useful for data preprocessing, e.g. FASTQ file mapping and quantification. The GUI provide an intuitive user interface that include an interactive file manager, communication window, and timeline tracker for data organization and progress observation, an embedded Jupyter notebook enabling dynamic analysis interaction and real-time result inspection, an embedded LaTeX editor/PDF viewer for writing assistance, and specialized visualization support for Pantheon-Evolve that displays live algorithm evolution progress with fitness trajectories and code mutation history (**Fig. S2**). It also features other windows such as the Teams, Knowledge and others for configuring agent teams, storing prior knowledge for specific fields, such as internally collected paper or generated code. The Slack interface can integrate the Pantheon system into the daily communication scenarios of research teams, facilitating timely access to research-related information during team collaboration for easier discussion. For the Application Layer 4, configuration-driven assembly allows rapid construction of domain-specific agentic systems by combining lower-layer components. Therefore, PantheonOS’ layered separation of design makes it serve as a general data science framework while being readily customizable for genomics and other domains. Lastly, Pantheon App Store enables community-driven component development and sharing. The definitions of skill, package, agent, and team on the App Store can be installed into any of the above interfaces with one click, thus facilitating the extension of the existing systems to new application cases (**Fig. 1B**).

**Figure 1.**
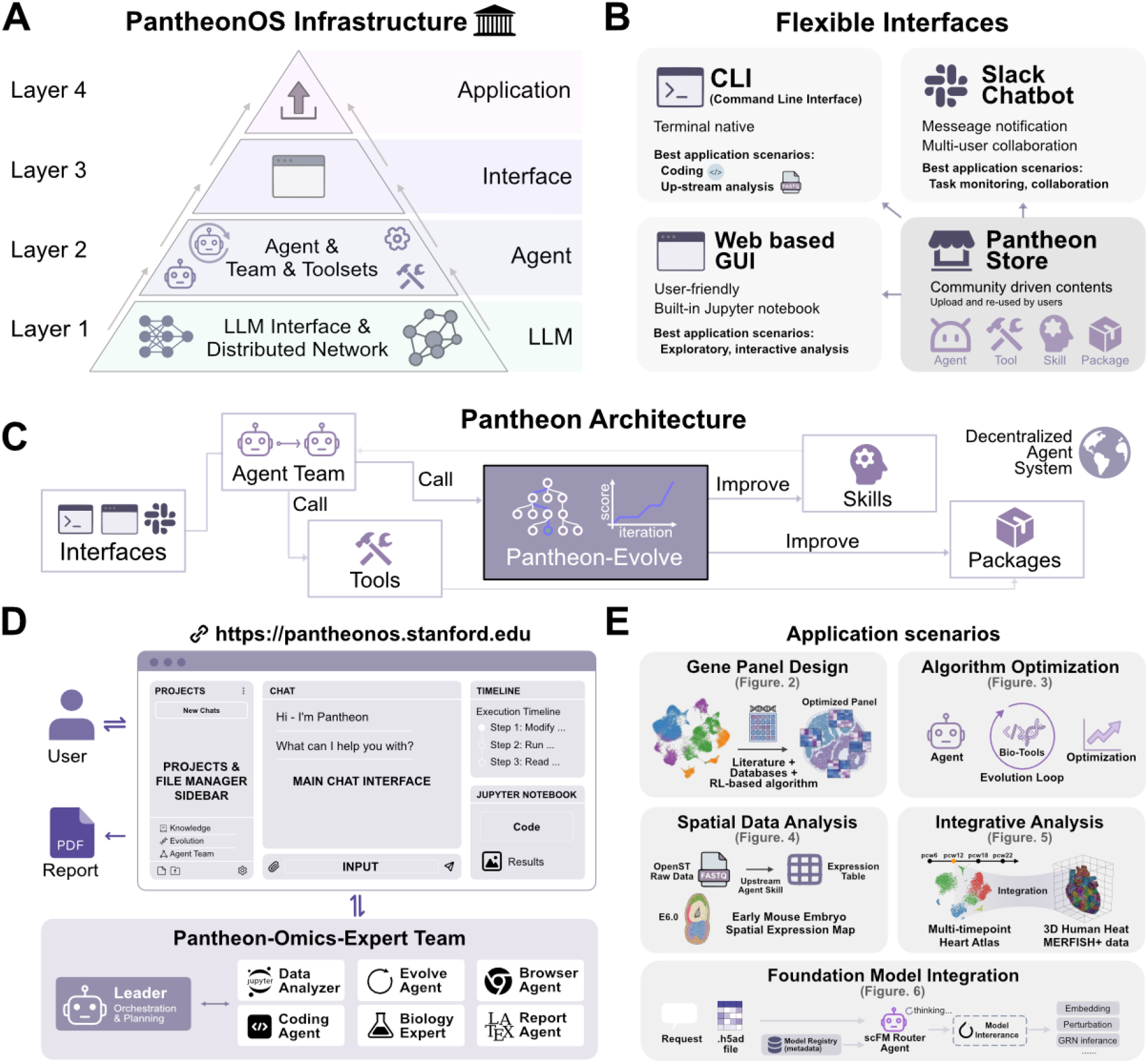
PantheonOS: an evolvable, distributed and integrative multi-agent system for automated data science studies, instantiated on single cell and spatial genomics analysis. **(A)** Four-layer pyramid architecture of Pantheon, from bottom to top, comprising LLM Interface and Distributed Network (Layer 1), Agent, Team and Toolsets (Layer 2, including evolvable agents), Interface (Layer 3), and Application (Layer 4) components. **(B)** Flexible user interfaces including command-line interface (CLI) for coding and upstream analysis, Slack chatbot for task monitoring and team collaboration, web-based GUI with built-in Jupyter notebook for interactive analysis, and Pantheon Store for community-driven agent system component (including domain specific skills, tools, agents or agent systems, vectorized knowledge database, GUI) sharing. **(C)** Core architecture workflow showing how user interfaces connect to agent teams, which leverage tools, the Pantheon-Evolve module and distributed communication to iteratively improve packages and agent skills, enabling a decentralized agent system. **(D)** The Pantheon-Omics-Expert agent team, featuring an agent Leader for orchestration and planning, Data Analyzer for workspace setup and dependency management, Evolve Agent for algorithm optimization, Biolog Expert for biological interpretation, Coding Agent for tool development, Browser Agent for literature retrieval, and Report Agent for automated report generation. **(E)** Automatic genomics data analyses enabled by PantheonOS: upstream fully automated reinforcement learning-based gene panel design for targeted spatial transcriptomics (**Fig. 2**), evolution of key single-cell genomics algorithms such as batch effect correction methods and even our RL algorithm for target panel design (**Fig. 3**), complete conversational analysis of new 3D dataset of early mouse embryogenesis (**Fig. 4**) and 3D spatial omics data of human fetal heart with multi-modal integration (**Fig. 5**); and finally moving towards the exciting direction of virtual cell through a unified access to diverse single-cell foundation models through intelligent routing (**Fig. 6**), promoting the integration virtual cell models and AI agents.

**Figure 2.**
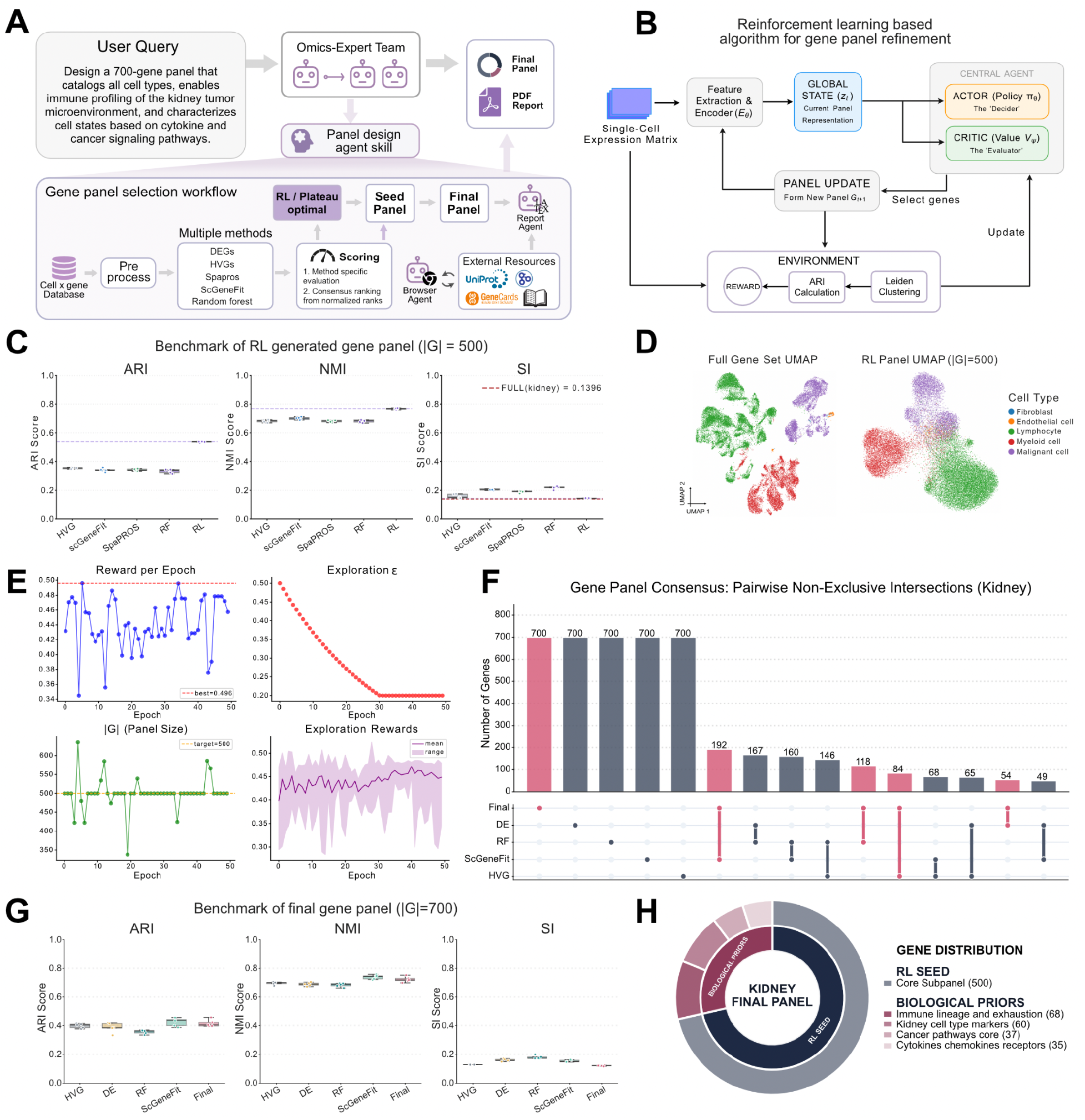
Pantheon-omics enables reinforcement learning-based gene panel design for immuno-oncology applications. **(A)** Workflow overview: user query specifying panel requirements is processed by the Omics-Expert Team, which employs a panel design agent skill that integrates an reinforcement learning algorithm (see Panel **B**) to generate and iteratively refine gene panels, and finally output a well-formatted PDF report. **(B)** Reinforcement learning algorithm architecture for gene panel refinement, showing the interaction between Global State encoder, Actor (Policy Network), Critic (Value Network), and Environment components that update the panel based on the Adjusted Rand Index or ARI of Leiden clustering, compared to ground-truth clusters, as a reward. **(C)** Benchmark comparison on the Kidney dataset showing RL-generated seed panel performance across ARI, NMI, and SI metrics against traditional methods. Note that the SI for the full data is low (0.1396) and the result from the RL based method is comparable. **(D)** Visualization of UMAPs with full gene set versus RL panel (G=500), demonstrating preserved cell type separation with the optimized panel. **(E)** Training Dynamics and Exploration Performance. The training process is characterized by four metrics across 50 epochs: Reward per Epoch (Top-Left): A snapshot of the evaluation reward at the start of each epoch, illustrating the inherent variance in initial policy performance (maximum reward achieved: 0.496). Exploration Strategy (ε-greedy) (Top-Right): The exploration probability (ε) decays exponentially from 0.50 to a minimum threshold of 0.20 at epoch 30. This strategy prioritizes stochastic environment traversal in early stages before transitioning to a more deterministic, policy-driven approach. Panel Size (|*G*|) (Bottom-Left): Tracking the panel size over time, with a target size of 500. The agent demonstrates increasing stability in maintaining the target size as training progresses. Exploration Rewards (Bottom-Right): Aggregate performance during exploration steps. The solid line represents the mean reward, while the shaded area denotes the reward range (min/max), indicating a gradual improvement in average exploration quality and a narrowing of performance volatility over time. **(F)** Upset plot of the pairwise non-exclusive intersection of gene panels from different methods, quantifying consensus among selection approaches. **(G)** Performance metrics (ARI, Normalized Mutual Information or NMI, and Silhouette Index or SI) of the agent-generated final gene panel compared to traditional methods. **(H)** Functional architecture of the final 700-gene kidney immuno-oncology panel, categorized by biological priors.

**Figure 3.**
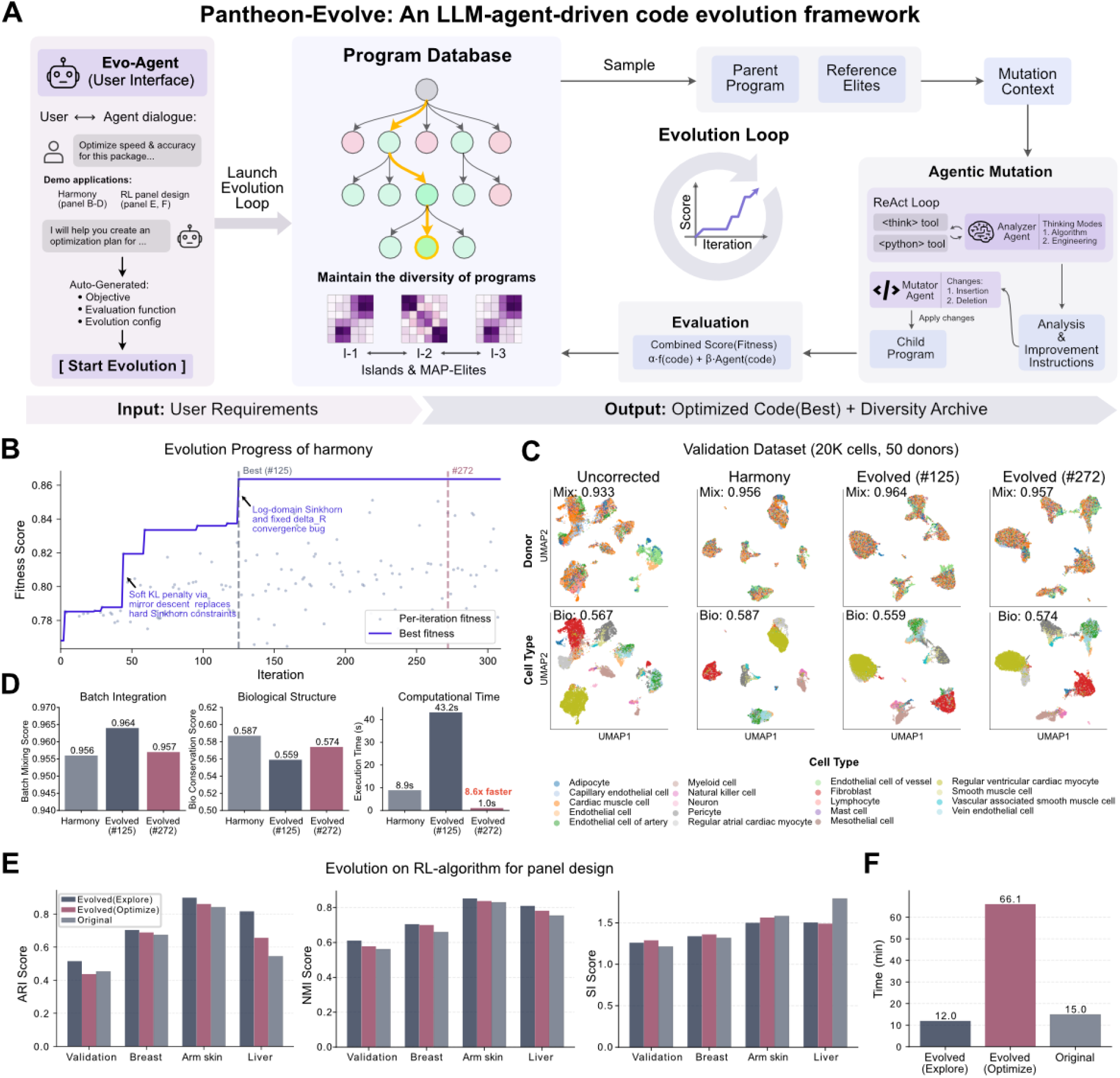
Pantheon-Evolve enables automated optimization of state-of-the-art bioinformatics algorithms through LLM-guided evolution. **(A)** Framework overview: user requirements are processed by the Evo-Agent interface, which auto-generates objectives, evaluation functions, and evolution configurations. The system maintains a Program Database using MAP-Elites to maintain code diversity, with an Evolution Loop that samples parent programs and reference elites to generate mutation context. Two-Phase Mutation employs an Analyzer Agent for strategic planning (Algorithm Optimization and Engineering Optimization) followed by a Mutator Agent for code modification, generating child programs for the archive. Hybrid Evaluation combines function-based metrics (accuracy, speed) with LLM-based code review, producing combined fitness scores. **(B)** Evolution trajectory of Harmony batch correction algorithm over iterations, showing fitness improvement trajectories for different evolved variants with progressive performance gains. **(C)** Performance on validation dataset (20,000 cells, 50 donors61) comparing uncorrected data, original Harmony, and two evolved variants (#125, #272), with metrics for batch mixing (Mix) and biological conservation (Bio) scores. Evolved variants achieve mixing scores of 0.964-0.967 while maintaining biological structure (Bio: 0.559-0.574). **(D)** Batch Integration Score comparison for Harmony evolution, demonstrating improved batch mixing across evolved variants. Biological Structure preservation scores for Harmony variants, showing maintained cell type separation after evolution. Computational time comparison for Harmony variants, with evolved versions achieving variable execution speeds while maintaining quality. **(E)** Performance comparison of the evolved RL-algorithm for panel design in terms of ARI, NMI, and SI metrics. Evolved (Explore) and Evolved (Optimize) are results obtained using different optimization objectives during evolution. **(F)** Comparison of training time between the evolved RL algorithm and the original algorithm

**Figure 4.**
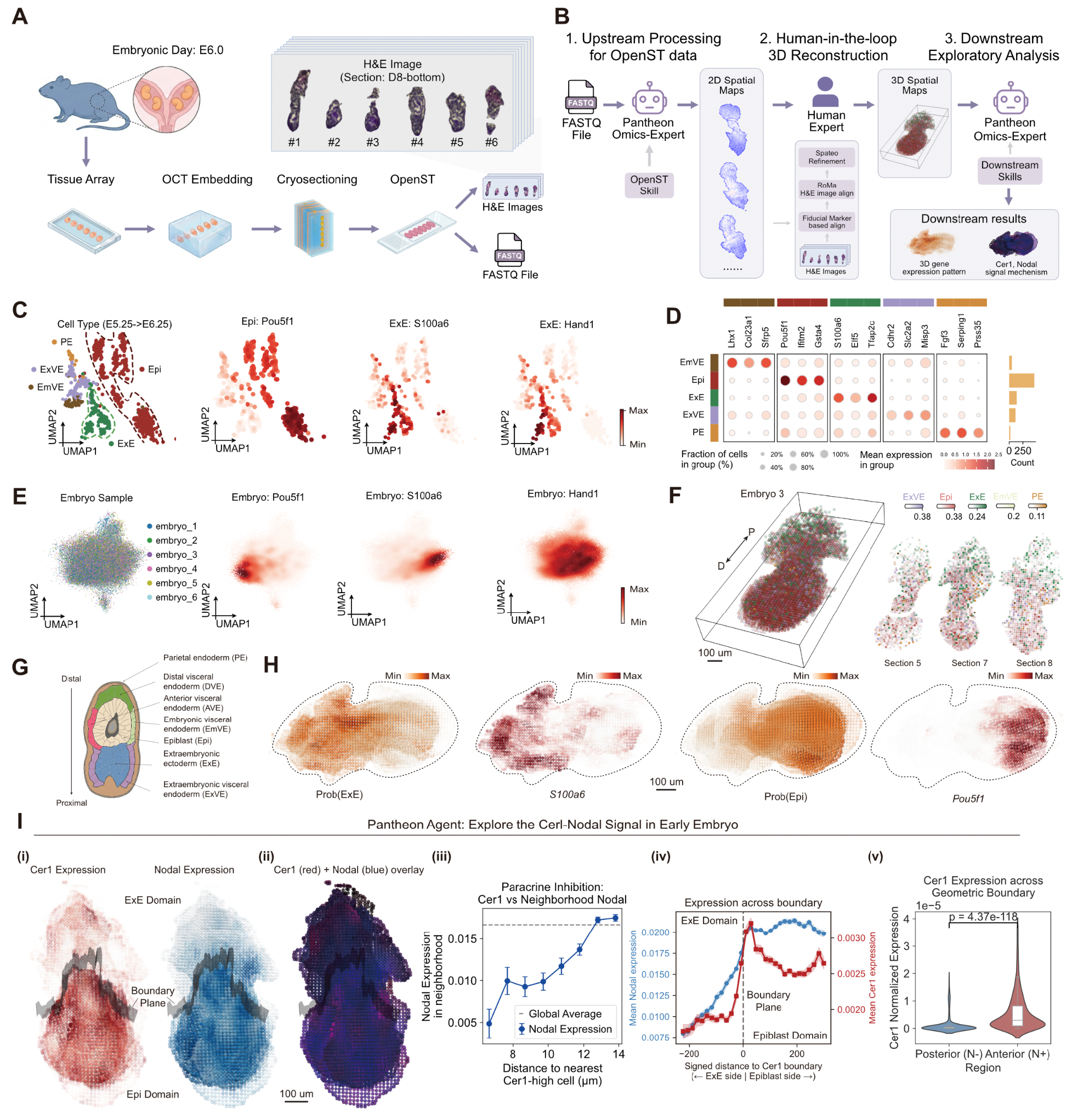
3D Spatial Transcriptomic Mapping of Cell Types, Gene Expression, and Signaling in the Early Mouse Embryo from E6.0. **(A)** OpenST spatial transcriptomics experimental workflow for profiling serial sections of six E6.0 mouse embryos co-embedded in a linear array. **(B)** Overview of the PantheonOS agentic computational analysis pipeline. (1) The Pantheon Omics-Expert agent automatically processes FASTQ data to generate 2D spatial maps. (2) Human-in-the-loop 3D reconstruction using sequential H&E images and Spateo refinement (See **Methods**). (3) The agent conducts downstream exploratory tasks such as spatial quantification of cell type, gene expression, and paracrine signalling. **(C)** UMAP visualization of single-cell RNA-seq reference data from the TOME (Transcriptome Of Mouse Embryos) atlas spanning E5.5–E6.2534. Cells are colored by cell type annotation (left panel): embryonic visceral endoderm (EmVE), epiblast (Epi), extraembryonic ectoderm (ExE), extraembryonic visceral endoderm (ExVE), and parietal endoderm (PE). Expression of the epiblast marker *Pou5f1* (center-left), and ExE markers *S100a6* (center-right) and *Hand1* (right) is shown on the UMAP embedding. **(D)** Dot plot showing the expression of selected marker genes across cell types in the TOME reference. Dot size indicates the fraction of cells expressing each gene; color intensity represents mean expression level. Bar plot (right) shows the total cell count per cell type. **(E)** UMAP visualization of six E6.0 embryos profiled in this study after batch correction using Pantheon-Evolve-Harmony, the evolved Harmony algorithm. Cells are colored by embryo of origin (left). Expression of *Pou5f1* (epiblast), *S100a6* and *Hand1* (ExE) confirms the expected lineage separation across the integrated dataset. **(F)** Spatial deconvolution of OpenST data (bin200 resolution) using Tangram35 for three representative sections from Embryo 3. Each bin is colored by the dominant (highest-probability) predicted cell type, and the accompanying composition bar on the top summarizes the predicted fraction of each cell type. The spatial arrangement recapitulates the expected tissue architecture of the E6.0 egg cylinder, with ExE in the proximal domain and epiblast in the distal region. **(G)** Schematic diagram of the E6.0 mouse embryo (egg cylinder stage) illustrating the spatial organization of cell types along the proximal–distal axes (figures adapted from *Srinivas77*). The epiblast occupies the distal portion of the egg cylinder, flanked by the embryonic visceral endoderm (EmVE), while the extraembryonic ectoderm (ExE) and extraembryonic visceral endoderm (ExVE) reside in the proximal domain. **(H)** Spatial visualization of Tangram-deconvolved cell type proportions and imputed marker gene expression for Embryo 3. Left two panels show the predicted epiblast (Epi) and ExE proportions per bin; right two panels display imputed expression of *Pou5f1* (epiblast marker) and *S100a6* (ExE marker) **(I)** Spatial analysis of the *Cer1*–*Nodal* signaling axis in the E6.0 embryo. (i) Spatially smoothed expression of *Cer1* and *Nodal* with the computationally derived *Cer1* expression boundary (transparent black plane). (ii) Dual-channel overlay of *Cer1* (red) and *Nodal* (blue) expression with the boundary (transparent black plane). (iii) Relationship between *Nodal* expression and the distance of these *Nodal* expressing cells from cells with high *Cer1* expression. (iv) Mean Nodal expression as a function of signed distance to the *Cer1* boundary (negative = ExE side/outside *Cer1* domain; positive = Epiblast side). Error bars represent SEM. (v) Violin and box plots comparing *Cer1* expression across two lateral domains (normal-positive (N+) and normal-negative (N-)) defined by a “blind” geometric axis (N) constructed perpendicular to the Proximal-Distal (P-D) boundary (ExE-Epiblast interface). P-value is calculated based on Mann-Whitney U test.

**Figure 5.**
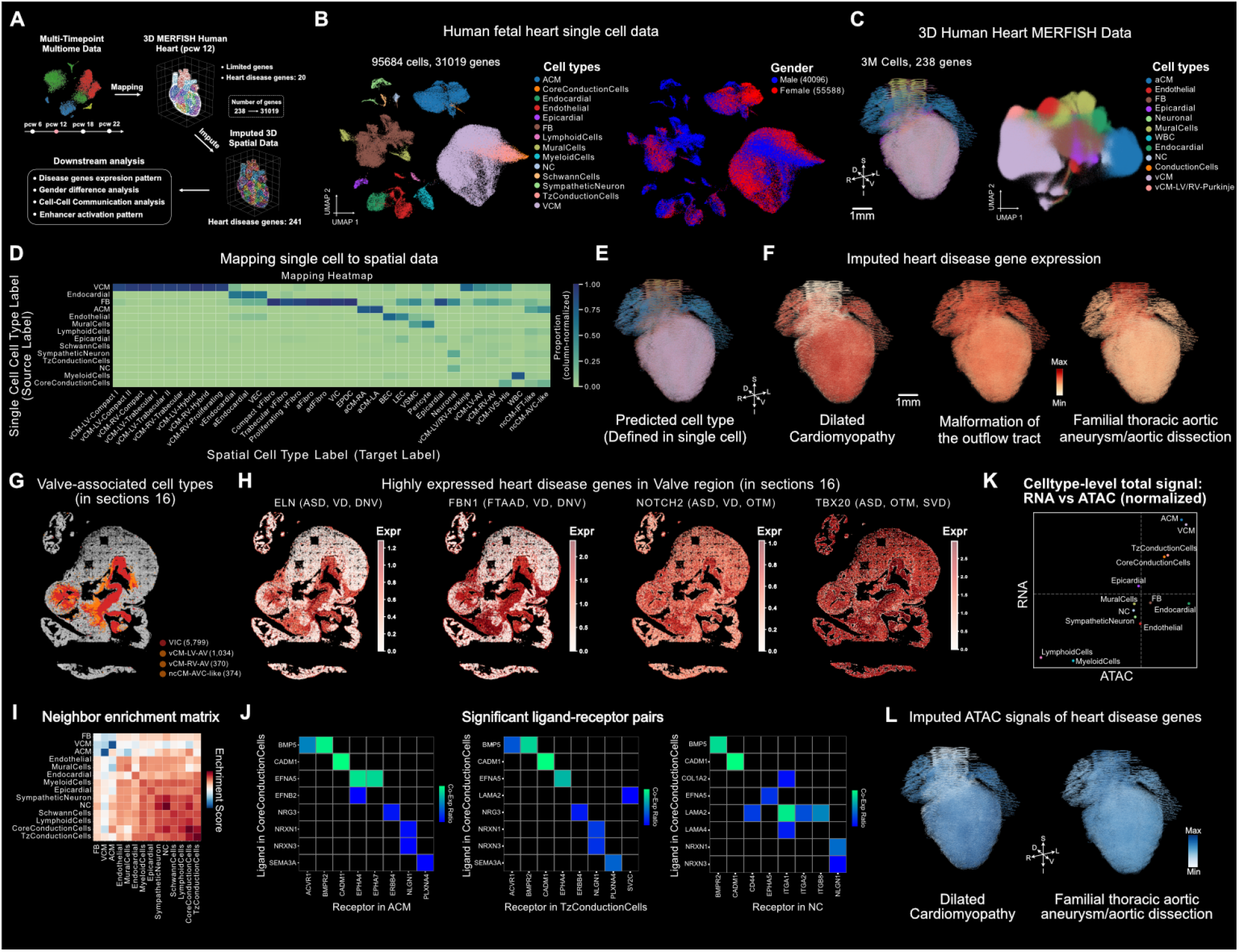
Pantheon-omics integrates human fetal heart single-cell multiomics and 3D MERFISH+ data of human fetal heart (post-conception week 12) for exploring congenital heart disease mechanisms. **(A)** Analysis workflow: multi-timepoint single cell multiomics data and 3D MERFISH human heart data are integrated through unbalanced optimal transport mapping for downstream 3D whole heart transcriptomic-level analyses including sex difference analysis, cell-cell communication, enhancer activation patterns, and imputed heart disease gene expression. **(B)** Human fetal heart single-cell dataset comprising 95,584 cells and all genes across multiple developmental timepoints, with UMAP visualization colored by cell types including fibroblasts, cardiomyocytes, endothelial cells, and endocardial cells.. **(C)** 3D Human Heart MERFISH+ dataset39 containing 3 million cells and 238 genes, visualized in 3D space with cell type annotations across multiple tissue layers. **(D)** Heatmap showing mapping results from single-cell to spatial data, with source cell types (rows) mapped to target spatial cell types (columns), enabling label transfer and expression imputation. The color of each cell on the heatmap corresponds to the proportion of the single cells from the multi-omics data mapped to the corresponding cell types derived from the spatial atlas. **(E)** Spatial visualization of predicted cell types in the 3D heart model. **(F)** Imputed heart disease gene expression patterns for Dilated Cardiomyopathy, Malformation of the outflow tract, and Familial thoracic aortic aneurysm/aortic dissection. **(G)** Cross-sectional view (Section 16) of the valve region, highlighting valve-associated cell types: VIC (valve interstitial cells, red, 5,799 cells), vCM-LV-AV (orange, 1,034 cells), vCM-RV-AV (yellow, 370 cells), and ncCM-AVC-like (pink, 374 cells). Non-valve cells are shown in gray. Section 16 has the highest VIC enrichment (16.1%) among all 53 tissue sections. **(H)** Imputed expression of four valve defect-associated genes in the valve region cross-section (Section 16): ELN (associated with ASD, VD, DNV), FBN1 (FTAAD, VD, DNV), NOTCH2 (ASD, VD, OTM) and TBX20 (ASD, OTM, SVD). Each gene displays a distinct spatial expression pattern within the valve region, with ELN concentrated in vascular smooth muscle areas, FBN1 in connective tissue/fibroblast domains, and NOTCH2, TBX20 broadly distributed across mesenchymal and pericyte populations. **(I)** Spatial neighbor enrichment matrix showing cell-cell proximity patterns. **(J)** Significant ligand-receptor pairs identified in spatial context. **(K)** Cell type-level total signal comparison between RNA and ATAC modalities (normalized). **(L)** Imputed ATAC signals for heart disease genes across disease categories.

**Figure 6.**
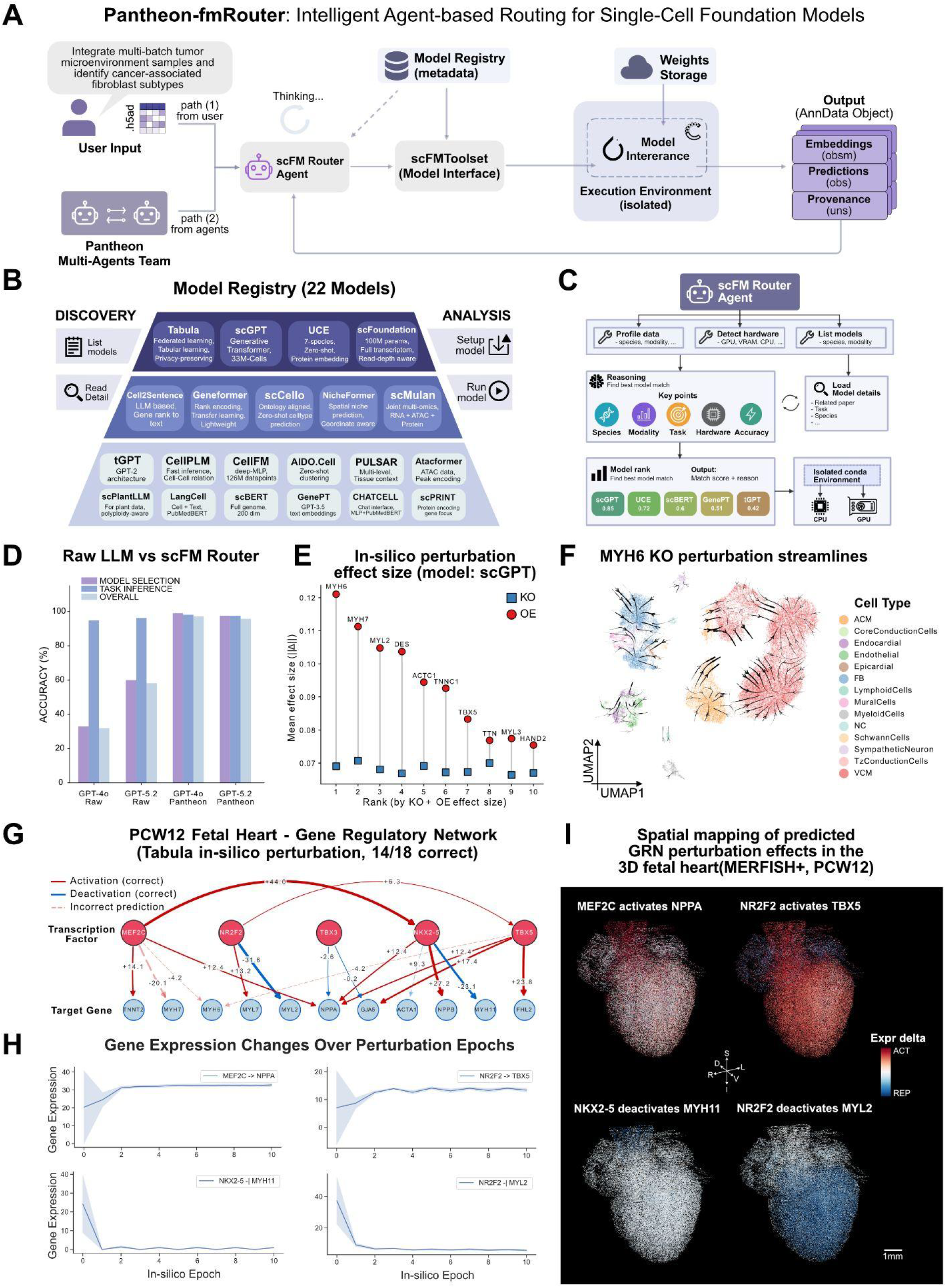
Pantheon-fmRouter provides intelligent routing to single-cell foundation models (scFM). **(A)** fmRouter architecture with two input paths: Path 1 receives user task descriptions along with datasets (e.g., h5ad files); Path 2 receives analysis requests from agents within the Pantheon Multi-Agents Team (e.g., Omics Expert Team). The scFM-Router agent interprets the task and previews the dataset, consults the Model Registry for model capabilities and requirements, selects the most appropriate model, and invokes scFMToolset to execute model inference. The router agent then interprets the outputs and returns results or feedback to the caller. **(B)** Model Registry containing 22 registered single-cell foundation models with two categories of tool operations: DISCOVERY (list available models, read model details) and ANALYSIS (setup model environment, run model inference). **(C)** Detailed workflow of the scFMRouter agent: (1) invoking tools to understand the dataset characteristics and local computing resources, then listing all available models; (2) comprehensive reasoning considering species, modality, task requirements, hardware constraints, and accuracy to select the most suitable model; (3) outputting a ranked list of models with scores and selection rationale; (4) executing inference using the selected model in an isolated environment, e.g. a conda environment, on the appropriate device (CPU or GPU). **(D)** Performance comparison between Raw LLM and pantheon scFM Router, with the vertical axis representing task execution accuracy. There are two different types of tasks: Model selection is used to measure the router’s ability to choose the appropriate model, while Task inference is used to measure the accuracy of scFM Router in identifying user intent, specifically in determining task types based on descriptions. **(E)** Effect sizes of the top ten genes identified through perturbation analysis using the scGPT model on 241 heart disease-related genes in human fetal heart single-cell data. **(F)** Perturbation streamlines of in-silico MYH6 knockout on fetal heart UMAP. Streamlines depict the direction and magnitude of scGPT embedding shifts upon MYH6 ablation, projected onto UMAP via kNN velocity graph. Line width scales with perturbation strength. Background points colored by cell type. ACM cells show strong directional flow, while VCM and FB populations exhibit centrifugal displacement toward cluster boundaries (n = 30,000 cells, PCW 11–13). **(G)** Predicted gene regulatory network from in silico perturbation of five cardiac TFs in PCW12 fetal heart scRNA-seq (10,000 cells) using Tabula. Red/blue edges indicate activation/repression; solid lines denote correct predictions and dashed lines incorrect predictions (accuracy: 14/18). Edge width and labels reflect the mean expression change across tested cells. **(H)** Representative in silico perturbation trajectories for four TF–target pairs. Mean target-gene expression (± SD) across epochs is shown for MEF2C→NPPA (activation), NKX2-5→MYH11 (repression), NR2F2→TBX5 (activation), and NR2F2→MYL2 (repression), demonstrating convergence of predicted expression changes within ∼3 epochs. **(I)** Spatial mapping of predicted perturbation effects onto the 3D fetal heart. Expression changes (epoch 9 − epoch 0) for NPPA, MYH11, TBX5, and MYL2 were imputed from scRNA-seq to MERFISH+ 3D coordinates via optimal transport. Red indicates upregulation(activation); blue indicates downregulation(repression).

We configured the Omics-Expert-Team to instantiate Pantheon for single-cell and spatial genomics analyses, comprising one Leader agent and six specialized sub-agents, namely Data Analyzer, Biology, Coding, Evolution, Browser, Reporter (**Fig. 1D**) that collaborate through autonomous task delegation. Through sub-agents, context generated from each agent will be kept without continuous accumulation, thus making it token-efficient. The leader agent receives user queries, analyzes requirements, designs execution plans, confirms with the user, dispatches tasks, monitors progress, and aggregates results. On the other hand, sub-agents play specific roles: Data Analyzer operates Jupyter notebooks for analysis pipelines; Biology Agent provides hypothesis generation and interpretation of the underlying biological system; Coding Agent builds reusable scripts; Evolution Agent triggers algorithm optimization; Browser Agent browses internet and retrieves literature; Report Agent generates publication-ready outputs. The Analyzer agent has a set of built-in skills related to omics data analysis (See **Tables S1** for and **Methods** for details). In addition, agents can find each other and delegate tasks, enabling autonomous decisions about when to invoke other agents. We demonstrate the power of Pantheon’s Omics-Expert-Team in enabling end-to-end genomics research across multiple systems, as detailed in the following sections (**Fig. 1D**).

To sum up, this hierarchical yet flexible design enables efficient, transparent, and user-controllable data science workflows while maintaining specialized genomics expertise across analysis aspects.

### Fully automated agentic gene panel design workflow with integrated reinforcement learning outperforms conventional methods

In order to demonstrate PantheonOS’ end-to-end single cell and spatial genomics data analyses capabilities, we start with the problem of gene panel design in imaging-based spatial transcriptomics (ST), where limited gene capacity creates a fundamental optimization challenge: how to design compact gene panels that preserve cell-type separability and biological interpretability while operating under strict measurement constraints. Existing gene panel selection approaches, such as Highly Variable Genes (HVG), Differential Expression (DE), and feature-importance–based methods including Random Forest (RF), SpaPROS^18^, and scGeneFit^19^, optimize isolated statistical criteria and operate independently. As a result, they frequently produce divergent panels and lack a unified framework for integrating complementary evidence or optimizing context-dependent biological objectives.

To address these limitations, we developed a fully automated, agentic gene panel design workflow within PantheonOS, implemented as a dedicated panel design agent skill orchestrated by the Omics-Expert Team (**Fig. 2A**). PantheonOS provides an adaptive orchestration layer that evaluates the dataset-specific performance of existing methods, arbitrates between their complementary strengths, and refines candidate panels under explicit biological constraints. In contrast to static ensemble strategies, combining e.g., HVG, DE, RF, SpaPROS or scGeneFit with fixed weighting schemes, the framework performs a context-aware meta-evaluation of competing approaches and selects and improves panels accordingly.

Given a user query specifying target panel size and optional biological constraints, the agent autonomously coordinates multiple gene selection strategies to form a core seed gene panel with Pareto optimum (see below) or an innovative reinforcement learning (RL) approach, which is then completed by explicitly optimize biologically relevant, context-dependent objectives. The workflow eventually outputs an automatically generated, publication-ready PDF report that documents panel composition, benchmarking results, and biological interpretation.

As an initial, agent-coordinated screening step, we evaluate multiple complementary gene-selection strategies mentioned above. Each method produces a ranked list of candidate genes according to its objective function. For each strategy independently, the panel design agent constructs incremental sub-panels of increasing size (100, 200, …, N), recomputes Leiden clustering using default Scanpy^20^ parameters on the training data, and evaluates clustering agreement against ground-truth cell-type annotations using the Adjusted Rand Index (ARI). The agent then analyzes ARI–panel size trade-off curves to identify Pareto-optimal seed panels, defined as gene sets for which clustering performance cannot be further improved without increasing panel size (**Fig. S3A**). This Pareto screening step yields compact, high-performing seed gene sets that balance statistical performance and panel parsimony. Importantly, rather than committing to a single heuristic output, the agent optionally aggregates Pareto-optimal genes across methods to form a structured candidate pool that serves as the initialization space for subsequent reinforcement learning–based combinatorial refinement.

At the core of this agentic workflow is an integrated reinforcement learning (RL) approach^21^ that enables adaptive combinatorial refinement beyond heuristic ranking (**Fig. 2B**). The RL skill formulates panel optimization as a sequential decision-making problem, in which panel modifications of the initial combined search space are proposed by a policy network and evaluated through re-clustering of the data. Clustering agreement with ground-truth annotations, quantified by ARI, serves as the reward signal, allowing the agent to directly optimize biologically meaningful cell-type separability.

We applied this agentic workflow to kidney cancer and immune oncology datasets where the latter has a well-curated gene panel to benchmark against (**Fig. 2** and **Fig. S3**). On the kidney dataset, the agent-generated RL seed panel overall outperforms conventional methods across ARI and normalized mutual information (NMI) (**Fig. 2C**). Despite using only 500 genes, the optimized panel preserves global and fine-grained cell-type structure comparable to embeddings generated from the full gene set (22,781 genes), as visualized by UMAP (**Fig. 2D**). These results demonstrate that the agent identifies compact yet information-rich gene combinations rather than redundantly expressed markers.

Analysis of training dynamics reveals that the RL skill converges to stable, high-performing solutions through an annealed ε-greedy exploration–exploitation strategy (**Fig. 2E**). High initial ε promotes stochastic exploration of the combinatorial search space, preventing premature convergence, while gradual ε decay shifts the policy toward deterministic exploitation. This transition stabilizes panel size and improves reward consistency, demonstrating the agent’s ability to discover robust solutions rather than overfitting to transient high-reward configurations.

While algorithmic screening ensures statistical predictability, biologically meaningful panel construction requires contextual grounding in domain knowledge. PantheonOS therefore introduces a structured biological retrieval phase that systematically integrates curated knowledge sources into the panel completion process (**Methods**), resulting in high biological relevance (**Fig. S2D, S3F**). This phase incorporates evidence from the scientific literature and established databases, including GeneCards^22^, UniProt^23^, and oncogene repositories. Comparative analysis across methods shows that while the agent-derived panels share partial overlap with traditional approaches, they also contain distinct gene subsets, as revealed by pairwise intersection analysis (**Fig. 2F**). Importantly, the final agent-generated panels achieve overall favorable performance across all evaluated metrics relative to heuristic baselines (**Fig. 2G**). Functional categorization of the finalized 700-gene kidney immuno-oncology panel further demonstrates coherent biological organization (Note that these 700 genes have shown to have highest statistical performance while the inclusion of additional 300 genes reduces the performance, see **Methods**), with genes grouped into immune, stromal, epithelial, and tumor-associated programs (**Fig. 2H**). Furthermore, in immune oncology study, PantheonOS is also shown to have the highest overlap with the expert-curated Vizgen panel compared to all other methods (23% gene overlap, **Fig. S3F**). Pantheon achieves superior performance across both predictive metrics while maintaining contextual biological alignment.

Together, these results establish an agentic paradigm for gene panel design, in which autonomous reasoning, integrated reinforcement learning, and biological context awareness collectively enable compact, high-performance panels that outperform conventional selection strategies while remaining biologically interpretable.

### Pantheon-Evolve: agentic code evolution improves batch correction and RL algorithms

Advances in computational algorithms have been instrumental in enabling and interpreting complex data, particularly that of single-cell genomics. Writing these algorithms, however, requires significant domain expertise and many iterations of formalization, implementation, and testing. To accelerate this process, several groups have combined LLMs with optimization ideas from genetic programming to create evolutionary agent harnesses, such as AlphaEvolve^24^, OpenEvolve^25^, and ShinkaEvolve^26^. Our approach takes inspiration from previous work, incorporating a Multi-dimensional Archive of Phenotypic Elites or MAP-Elites program^27^ database with genetic programming as an optimization framework. Previous implementations use two different LLM calls for evolution planning and code modification. We use two Reasoning and Acting or ReAct agents^1^ as modifiers instead of direct LLM calls, a planning agent and an analyzer agent. To more closely mimic human algorithmic development, we designed the analyzer agent with two prompts: high-level ideation and code-level optimization (**Fig. 3A**). The full pseudocode for our algorithm is detailed in **Algorithm 5** (**Methods**). The Pantheon-UI allows for easy tracking of datasets required for evolution and tracking of evolutionary progress (**Fig. 1E**). To demonstrate the power of Pantheon-Evolve, we turned to two case studies: batch correction and the gene panel selection algorithm previously detailed (**Fig. 2A**).

We choose batch correction algorithms for our first case study because there are many extant batch correction algorithms used in the single-cell community and there are commonly used performance metrics for batch correction^28^. When expressed in code, batch correction algorithms are diverse enough to demonstrate single-file and multi-file evolution. Perhaps most importantly, batch correction algorithms have previously been used as an evolution target, and there is a large body of literature on benchmarking them^29^, making our efforts directly comparable. We applied Pantheon-Evolve to Harmony^7^, a well-known batch correction algorithm that uses cycles of soft clustering and centroid-based correction in PCA space. To conduct evolution and validation, we chose a specific subset of the CELLxGENE dataset that has been used in previous work for batch correction evolution due to its consistency with the Open Problems benchmark datasets^30^. Within the dataset, we normalized expression, identified highly variable genes, and calculated principal components. We then selected two disjoint subsets of 20,000 cells to be used for training and validation. These datasets included 19 cell types and 50 donors (**Fig. 3B, C**). We conducted Harmony evolution with an evolution objective that was selected for batch mixing, cell type conservation, and speed (**Methods**). Over 300 rounds of evolution at a cost of about $25, Pantheon-Evolve incorporated a log-domain Sinkhorn transport plan for aligning cells, improving batch integration performance on the training set (**Fig. 3B, C**). We display validation results for two program candidates, #125 and #272, as notable: #125 had improved batch integration performance, while #272 displayed an 8.6x increase in batch correction speed (**Fig. 3D, Tables S2, 3**). In addition to Harmony, we applied Pantheon-Evolve to Scanorama^8^ and BBKNN^9^, other batch correction algorithms, and observed more modest increases in performance (**Fig. S4, Tables S4-7**). In both cases, however, Pantheon-Evolve makes it possible to trial many innovative ideas in a short amount of time. Collectively, these results demonstrate Pantheon-Evolve’s ability to make rapid improvements to ubiquitous single-cell bioinformatics algorithms.

Considering published algorithms have been through many rounds of optimization and review, and their details may be described in LLM training corpuses, we next sought to examine whether Pantheon-Evolve could evolve a complex and novel algorithm, the RL augmented gene panel selection algorithm we developed. We applied Pantheon-Evolve to a GPU-optimized version of the gene panel selection algorithm. At baseline, this algorithm took around 20 seconds to run on an H100 GPU. We used 10,000 cell subsets of the kidney dataset previously described (**Methods**). To evaluate robustness, we leveraged other tissues from the same dataset, consisting of cells from breast, arm, and liver tissue. We then conducted evolution with evolution objectives that measured ARI, NMI, SI, and training speed (**Methods**). As the RL algorithm incorporates multiple components, we explicitly prompted the agent to focus either on the Explore or Optimize subroutines. The Explore subroutine generates new candidate gene panels and retrains the autoencoder that produces the state representation. The Optimize subroutine updates the actor and critic networks used in the reinforcement learning algorithm, fitting the critic using a temporal-difference step and updating the actor using a policy gradient. Over 50 iterations of evolution at a cost of about $4, Pantheon-Evolve is capable of making major changes to the algorithm. For example, in both Optimize and Explore prioritized evolutions, Pantheon-Evolve converges toward a Cross-Entropy Method distribution search with Gumbel-TopK sampling. Compared to the Optimize priority prompting, the Explore prioritized evolution introduces several new ideas, such as a Jaccard similarity of every candidate against the current best, rejecting similar candidates, and multiple methods for generating candidate pools in each round, including local mutations of existing panels and prior-preserving samples (**Tables S8, 9**). The modified algorithms achieve favorable ARI and NMI scores across the validation and hold out datasets (**Fig. 3E**), albeit at the cost of comparable or higher complexity (**Fig. 3F**). These results demonstrate Pantheon-Evolve’s ability to make beneficial changes to a novel algorithm that requires changes to model architecture.

In conclusion, we show that Pantheon-Evolve not only enhances highly optimized, state-of-the-art single-cell batch correction methods, including Harmony, Scaronoma, and BBKNN, but also substantially improves nascent reinforcement-learning–based gene panel design algorithms developed in this work, at a fraction of the human labor required. These results demonstrate that evolvable, LLM-guided agents can operate directly on mechanistic code, yielding systematic performance gains across both mature and emerging algorithmic paradigms. More broadly, Pantheon-Evolve points toward a new model of algorithm development in which autonomous, evolvable agents accelerate innovation, expand the space of viable solutions, and amplify human scientific creativity.

### PantheonOS automates 3D spatial mapping of symmetry-breaking molecular and signaling events during early gastrulation

The E6.0 mouse embryo represents a uniquely informative developmental window in which body-axis specification, germ-layer initiation, and spatial regulation emerges from simultaneous signaling events, making it an ideal system to resolve how global embryonic structure arises from local molecular interactions. To resolve the precise spatial architecture and intercellular signaling of the E6.0 mouse embryo, we utilized OpenST^31^ to generate high-resolution transcriptomic maps across six independent embryos organized in a linear array (**Methods, Fig. 4A**). The analysis was orchestrated by PantheonOS’ Omics-Expert-Team, which autonomously synthesized a comprehensive 3D spatial transcriptomics pipeline by integrating core skills built from OpenST preprocess pipeline^31^, Spateo^32^, Scanpy^20,32^, and OmicVerse^33^ to facilitate end-to-end 3D ST analysis, particularly single cell and spatial data integration, cell type deconvolution, signal boundary identification, and biological interpretation (**Fig. 4B**). Specifically, PantheonOS firstly utilized the TOME^34^ (Trajectories Of Mouse Embryos) reference atlas spanning E5.25–E6.25, which provides a comprehensive cell-type catalog of early post-implantation development, including the extraembryonic ectoderm (ExE), extraembryonic visceral endoderm (ExVE), embryonic visceral endoderm (EmVE), and epiblast (Epi) (**Fig. 4C**). PantheonOS then characterized these cell types with a detailed dotplot quantifying marker specificity across all cell types (**Fig. 4D**). This analysis revealed distinct expression signatures, such as *Lhx1* and *Col23a1* for the EmVE, *Pou5f1* and *Ifitm2* for the epiblast, and *Elf5* and *S100a6* for the ExE.

Following reference establishment, we integrated data from the E6.0 embryo array using the optimized Harmony from Pantheon-Evolve and PantheonOS’ Deconvolution Skill. This evolved algorithm successfully mitigated batch effects while preserving the lineage-specific separation of the epiblast and extraembryonic tissues, as evidenced by the spatially distinct expression of *Pou5f1, S100a6*, and *Hand1* within the integrated UMAP embedding (**Fig. 4E**). To assign the cell type identity for each bin of the OpenST spatial profile, PantheonOS proceeds with Tangram deconvolution^35^, accurately revealing the proximal-distal axis of the embryo characterized by the enrichment of ExE and Epi cells, respectively (**Fig. 4F**). This computational reconstruction was highly concordant with our schematic diagram of the E6.0 mouse embryo (**Fig. 4G**), which illustrates the spatial organization of cell types along the proximal–distal axis. In this model, the epiblast occupies the distal portion of the egg cylinder, flanked by the EmVE, while the ExE and ExVE reside in the proximal domain (**Fig. 4G**). To confirm the robustness of these spatial assignments, we visualized Tangram-deconvolved cell type probability alongside imputed marker gene expression for Embryo 3 (**Fig. 4H**). The predicted probability for the epiblast and ExE domains showed high spatial overlap with the imputed expression of their respective markers, *Pou5f1* and *S100a6* (**Fig. 4H**). This strong concordance between cell type assignment and marker expression patterns demonstrates the fidelity of the agent-orchestrated spatial reconstruction and analyses.

To characterize the complex signaling landscape of the E6.0 embryo, PantheonOS established a dual-axis geometric framework to quantify morphogen gradients and axial emergence. By computationally deriving a spatial boundary for *Cer1* (an antagonist secreted by the Anterior Visceral Endoderm (AVE), which functions as the anterior signaling center, first emerges at the distal visceral endoderm and subsequently migrates laterally to the future anterior side)^36^, the agent quantified the antagonistic relationship between *Cer1* and the morphogen *Nodal* (**Fig. 4I-i, ii**)^37^. First, a spatial Boundary Plane was computationally derived to bisect the embryo into ExE and Epiblast domains (**Fig. 4I-i, ii**). Analysis across this interface then revealed a robust paracrine inhibition signature, where *Nodal* expression exhibits a sharp, distance-dependent sigmoidal increase as cells transition from the ExE side into the Epiblast domain, with *Cer1* levels peaking at the boundary, revealing a Proximal-Distal (P-D) axis patterning (**Fig. 4I-iii, iv**). To quantify the break in radial symmetry within the distal epiblast region, a second geometric axis was constructed perpendicular to the P-D boundary. This axis is defined by the Normal vector (N), which originates from the boundary plane and points toward the embryo’s interior. By utilizing this “blind” bisection, PantheonOS established a coordinate system where N^+^ (Normal positive) and N^-^ (Normal-negative) represent two lateral domains defined solely by their orientation relative to the vector, independent of initial marker expression. This approach unveiled a lateral symmetry breaking, quantitatively confirming that signaling molecules like *Cer1* are not uniformly distributed but are sequestered to a specific geographic hemisphere (N^+^) of the embryo. Additional comparison further confirmed the localized nature of the AVE, showing that *Cer1* is significantly sequestered within the N^+^ domain (**Fig. 4I-vi**). Conversely, *Nodal* exhibits a complementary distribution, with increased expression in the N^-^ domain where the inhibitory influence of *Cer1* is absent (Mann-Whitney U test, **Fig. 4I-v**). These results demonstrate that the PantheonOS can autonomously resolve complex paracrine signaling interfaces and identify symmetry-breaking events by quantifying the non-homogeneous distribution of molecules within the intact embryonic disc profiled with OpenST in 3D space.

Collectively, our results demonstrate that PantheonOS autonomously reconstructs the spatial and signaling architecture of the E6.0 embryo array by integrating eight sections with the TOME single-cell reference. Utilizing Pantheon-Evolve optimized Harmony and PantheonOS’ Tangram Deconvolution Skill, the agent accurately delineated the proximal–distal axis of the egg cylinder, yielding a computational reconstruction highly concordant with the known biological schematic and validated by the spatial overlap of canonical markers like *Pou5f1* and *S100a6*. Furthermore, the system revealed that *Cer1* actively drives robust paracrine inhibition to suppress *Nodal* expression, while simultaneously uncovering a marked asymmetric distribution of *Cer1* perpendicular to the boundary. This dual spatial characteristic defines the precise territorial limits of morphogen gradients, effectively mapping the symmetry-breaking events during early gastrulation.

### Multi-modal integration by PantheonOS reveals spatially-resolved gene signatures and sex-specific expression patterns of heart disease genes in 3D fetal heart

To demonstrate Pantheon’s capability for sophisticated multi-modal spatial analysis, we completed a complex multi-omics integration analysis (**Fig. 5A**) through interactive dialogue with the Omics-Expert-Team to analyze the expression patterns of genes associated with heart diseases, especially congenital heart disease (CHD), in the human fetal heart. We integrated a single-cell multiomics dataset^38^ (95,684 cells from post conception weeks (PCW) 11 to 13; **Fig. 5B**) with a 3D MERFISH+ spatial dataset^39^ at post-conception week 12 (3 million cells, **Fig. 5C**). Because the 3D MERFISH+ study captures only 238 genes, of which merely 20 overlap with the 241 known heart disease genes based on the curated from CHDgene database and others^38,40^, direct spatial analysis of disease genes from the spatial data alone is infeasible, motivating cross-modal integration to impute transcriptome-wide expression into the 3D spatial context (**Fig. 5A**). PantheonOS invoked its built-in single-cell-to-spatial mapping skill to perform unbalanced optimal transport via Moscot^41^, enabling bidirectional transfer (between MERFISH+ data and single cell multi-omics data) of cell type labels and gene expression between the two modalities. The resulting mapping heatmap (**Fig. 5D**) shows biologically coherent correspondence: for example, ventricular cardiomyocytes (VCM) from the single-cell data map predominantly to VCM-related spatial populations, and similar consistency is observed across other major cell types. PantheonOS further predicts cell type labels projected onto the 3D heart model (**Fig. 5E**) recapitulate the expected anatomical distribution of cell types, e.g., atrial, ventricular and outflow tract related cells are mapped to the corresponding anatomical structures, confirming the fidelity of the optimal transport integration.

Next, PantheonOS used the integrated data to examine the spatial expression patterns of sets of genes associated with different types of heart disease (**Fig. 5F; Fig. S5A, B**). We examined sets of genes associated with several well-studied adult diseases (hypertrophic cardiomyopathy (HCM), dilated cardiomyopathy (DCM), and familial thoracic aortic aneurysm/aortic dissection (FTAAD)), where the causal genes and cell types are well characterized, as well as genes associated with different heart diseases. However, the causal cell types and their spatial locations are less well understood. Cell type-resolved analysis across all 34 MERFISH-defined cell types (**Fig. S5A**) reveals that sets of genes associated with different heart diseases exhibit distinct cell type expression profiles: genes associated with single ventricle disease (SVD), HCM, and DCM show highest expression in cardiomyocyte subtypes (vCM-LV, vCM-RV. LV/RV: left or right ventricle), while genes associated with malformation of the outflow tract (OTM) are preferentially expressed in vascular and outflow tract-related populations. In 3D space, DCM-associated genes show preferential expression in the ventricular wall regions where cardiomyocytes predominate, consistent with the known myocardial basis of this disease (**Fig. 5F, left**). Genes linked to OTM display concentrated signals in the superior cardiac structures corresponding to the outflow tract region (**Fig. 5F, middle**). FTAAD-associated genes are enriched in the great vessel and aortic regions of the heart (**Fig. 5F, right**). Extending to additional heart disease categories (**Fig. S5B**), atrial septal defect (ASD) and ventricular septal defect (VSD) modules show expression concentrated in septal regions, valve defect (VD) genes localize to valve-proximal areas, atrioventricular septal defect (AVSD) genes highlight the atrioventricular junction, and single ventricle disease (SVD) genes show broad ventricular enrichment; PCGC de novo variant (DNV-PCGC) genes display a more diffuse but cardiomyocyte-biased pattern. Focusing on the valve region, the anatomical site most relevant to valve defects, PantheonOS identified Section 16 of MERFISH+ data as the cross-section with the highest valve interstitial cell (VIC) enrichment (16.1% valve-associated cells; **Fig. 5G**) and visualized the expression of four genes jointly linked to valve defects: *ELN* (associated with ASD, VD, DNV), *FBN1* (FTAAD, VD, DNV), *NOTCH2* (ASD, VD, OTM) and TBX20 (ASD, OTM, SVD) (**Fig. 5H**). These genes display complementary spatial distributions within the valve cross-section, *ELN* marks vascular regions, *FBN1* delineates connective tissue domains, and *NOTCH2, TBX20* highlights pericyte and interstitial zones, suggesting distinct but spatially coordinated cellular mechanisms underlying valve pathology.

To further dissect these disease modules–level patterns at single-gene resolution, we ask the Omics-Expert Team to systematically profile the expression of all 241 heart disease genes across the 34 MERFISH-defined cell types (**Fig. S6A**). This analysis reveals that heart disease genes segregate into functionally coherent expression clusters: sarcomeric genes (*TTN, MYH7, ACTC1, TNNT2, MYL2, ACTN2*) associated with DCM and HCM are almost exclusively expressed in ventricular cardiomyocytes, while transcription factors such as *TBX5* and *PITX2* display atrial-restricted expression, and vascular genes (*ELN, FBN1*) are confined to stromal and smooth muscle populations. Quantification of the number of heart disease genes with peak expression per cell type (**Fig. S6B**) confirms that ventricular cardiomyocytes (vCM-RV-Compact: 27 genes; vCM-RV-Proliferating: 26 genes) host the largest share of peak-expressed disease genes, consistent with the central role of cardiomyocyte dysfunction in cardiomyopathies, while non-myocyte populations including WBC (15 genes), BEC and adFibro (14 genes each) also harbor substantial numbers, highlighting the diverse cellular origins of heart disease pathogenesis. Visualization of the top 5 most highly expressed genes per disease trait (**Fig. S6C**) further reveals a dichotomy between cardiomyopathy traits (DCM, HCM), which are dominated by highly expressed, cell-type–specific sarcomeric genes with high coefficient of variation (CV) across cell types, and structural defect traits (ASD, VSD, SVD), which feature more broadly expressed transcription factors and chromatin regulators. Representative 3D expression maps of individual genes (**Fig. S6D**) illustrate distinct spatial signatures: *TTN* shows broad ventricular expression, *PITX2* is restricted to the left atrium consistent with its established role in left–right patterning, *TBX5* displays atrial enrichment, and *ELN* localizes to vascular smooth muscle regions surrounding the great vessels.

To characterize the spatial organization of cell types in the 3D fetal heart, PantheonOS computed pairwise cell type proximity statistics using kNN-based neighbor enrichment analysis (**Fig. 5I**), revealing preferential co-localization patterns such as the spatial association between cardiomyocytes and endocardial cells. Ligand-receptor interaction analysis via Spateo^32^ identifies significant spatially-informed communication pairs across cell types (**Fig. 5J**): for instance, *BMP5, CADM1, EFNA5, NRG3*, and neurexin family members (*NRXN1, NRXN3*) emerge as prominent ligands in CoreConductionCells communicating with receptors in atrial cardiomyocytes (ACM), transmission zone conduction cells (TzConductionCells), and neural crest cells (NC). These spatial communication patterns implicate neurodevelopmental and BMP signaling pathways in the coordinated development of the cardiac conduction system.

Because the single-cell multiomics data includes both RNA and ATAC modalities, PantheonOS further imputed heart disease-associated enhancer chromatin accessibility signals into 3D space using the scE2G^42^ enhancer-gene links and the same optimal transport framework. Comparison of cell type-level total RNA versus ATAC signals (**Methods, Fig. 5K**) reveals that most cell types show correlated transcriptomic and epigenomic activity, while certain populations such as lymphoid and myeloid cells exhibit notable discordance (marked with asterisks), suggesting post-transcriptional regulation or epigenetic priming in these lineages. Cell type-resolved ATAC profiles across all 10 heart disease gene sets (**Fig. S6E**) show that chromatin accessibility patterns partially mirror but do not fully recapitulate the RNA-based expression profiles (**Fig. S5A**): HCM and DCM modules maintain strong cardiomyocyte-enriched ATAC signals, while modules such as AVSD, ASD, and VSD show comparatively more uniform accessibility across cell types, suggesting that enhancer priming for these disease gene programs may be broader than their transcriptional output. In 3D space, ATAC patterns for DCM and FTAAD (**Fig. 5L**) show broadly similar but not identical distributions compared to their corresponding RNA-based expression maps (**Fig. 5F**); extended visualization of six additional modules (**Fig. S6F**) further confirms disease-specific spatial chromatin landscapes, for instance, SVD and HCM show strong ventricular ATAC enrichment, while OTM chromatin accessibility is concentrated in the outflow region. This multi-layered spatial comparison of transcriptomic and epigenomic signals demonstrates PantheonOS’s ability to dissect the regulatory architecture of disease genes at the 3D tissue level, revealing both concordant active regulation and discordant poised enhancer states across heart disease categories.

Collectively, this analysis, driven entirely through conversational interaction with the Omics-Expert-Team, demonstrates PantheonOS’s capacity to orchestrate complex, multi-modal spatial analyses that integrate single-cell multiomics and 3D spatial profiling data, yielding spatially resolved insights into congenital heart disease mechanisms, sex-specific molecular patterns, cell-cell communication, and epigenomic regulation without manual programming.

### Intelligent router automates optimal virtual cell model selection across heterogeneous single-cell and spatial analysis tasks

The rapidly expanding ecosystem of virtual cell models^10^, epitomized by transcriptomics data based single-cell foundation models (scFMs), each with distinct architectures, training dataset, species support, modality coverage, and hardware requirements, presents a growing challenge for researchers seeking to select the most appropriate model for a given analysis task. To address this, PantheonOS implements an intelligent scFM Router agent that automatically selects and executes the optimal foundation model from an expandable registry (see **Methods**) of 22 leading models spanning diverse architectures and capabilities (**Fig. 6A, B, Tables S10**). The router accepts input through two paths: directly from users who provide task descriptions along with datasets, or from other agents within the PantheonOS multi-agent team (e.g., the Omics-Expert-Team) that delegate foundation model tasks programmatically (**Fig. 6A**). The routing workflow follows a structured pipeline (**Fig. 6C**): the agent first profiles the input dataset (species, modality, gene ID format) and detects available hardware resources (GPU, VRAM, CPU), then queries the Model Registry to retrieve model capabilities and requirements, performs multi-criteria reasoning across species compatibility, modality support, task requirements, hardware constraints, and expected accuracy, and finally outputs a ranked list of candidate models with scores and selection rationale before executing inference in an isolated conda environment.

To quantify the advantage of the agent-based routing approach over direct LLM prompting, we benchmarked the virtual cell or vc Router against raw LLM baselines (GPT-4o and GPT-5.2 without Pantheon’s vcRouter infrastructure) on two evaluation axes: model selection accuracy (whether the correct model is chosen for a given task) and task inference accuracy (whether the system correctly identifies the user’s analytical intent from natural language descriptions). Raw LLMs achieve moderate task inference accuracy but substantially lower model selection accuracy, reflecting their limited knowledge of the specific capabilities and constraints of individual scFMs (**Fig. 6D**). In contrast, the Pantheon vcRouter achieves near-perfect accuracy on both metrics across both backbone LLMs (GPT-4o and GPT-5.2), demonstrating that the structured registry lookup, data profiling, and multi-criteria reasoning pipeline robustly compensates for gaps in the LLM’s parametric knowledge (**Fig. 6D**). These results confirm that agentic tool use, rather than larger or newer models alone, is the key driver of reliable foundation model selection.

To illustrate the router’s utility in a biologically meaningful application, we used PantheonOS to perform *in-silico* gene perturbation analysis on the human fetal heart single-cell data using scGPT^43^, which was automatically selected by the router for this perturbation task. Perturbation effect sizes for 241 heart disease-associated genes^38,40^ reveal that cardiac structural genes, *MYH6, MYH7, MYL2, DES*, and *ACTC1*, exhibit the largest embedding shifts upon both knockout (KO) and overexpression (OE), consistent with their central roles in cardiomyocyte identity and function (**Fig. 6E**). Streamline visualization of the *MYH6* knockout perturbation (**Fig. 6F**) shows that ACM cells exhibit strong directional flow upon *MYH6* ablation, while VCM and fibroblast populations display centrifugal displacement toward cluster boundaries, suggesting cell state destabilization in cardiomyocyte lineages. Extended perturbation streamline analysis across five top-ranked genes (*MYH6, MYH7, MYL2, DES, ACTC1*)^44^ under both OE and KO conditions (**Fig. S7A**) reveals gene- and perturbation-direction-specific cell state transitions: overexpression generally drives convergent flow toward cardiomyocyte cores, while knockout induces divergent trajectories away from lineage centers, with each gene affecting distinct cell type subsets.

To further demonstrate cross-model routing, we next tasked PantheonOS with gene regulatory network (GRN) inference, for which the router automatically selected Tabula^45,46^, a tabular learning based foundation model that explicitly validates its high performance in dissecting regulatory networks across many biological systems, including cardiogenesis, as the most suitable model for in-silico perturbation-based GRN prediction. In-silico perturbation of five well-characterized cardiac transcription factors (*MEF2C, NR2F2, TBX3, NKX2-5, TBX5*)^44^ in PCW12 fetal heart data (10,000 cells) produces a predicted GRN with 14 out of 18 edges correctly matching literature-validated regulatory relationships^47^ (**Fig. 6G, Tables S11**). The network captures both activation (e.g., *MEF2C* activating *TNNT2* with +44.0 mean expression change, *NR2F2* activating *TBX5*) and repression (e.g., *NKX2-5* -| *MYH11* with −20.1 change, *TBX3* -| *GJA5*) with biologically consistent directionality and magnitude (**Fig. 6G**). Representative perturbation trajectories (**Fig. 6H**) show that predicted expression changes converge within approximately 3 epochs: *MEF2C*→*NPPA* and *NR2F2*→*TBX5* exhibit progressive upregulation consistent with activation, while *NKX2-5* -| *MYH11* and *NR2F2* -| *MYL2* shows downregulation consistent with repression. Additional TF-target pairs (**Fig. S7B**) further validate the approach: *MEF2C*→*NKX2-5* and *MEF2C*→*TNNT2* pairs show robust activation trajectories, while *TBX3 -*| *GJA5* and *TBX3 -*| *NPPA* pairs demonstrate repression dynamics, all converging to stable expression levels within the perturbation window.

To connect the predicted regulatory effects back to the 3D tissue context, PantheonOS imputed the Tabula-predicted expression changes (predicted expression from epoch 9 minus original expression from epoch 0) onto the 3D MERFISH+ fetal heart atlas via the previously established optimal transport mapping. Spatial visualization reveals anatomically coherent perturbation patterns (**Fig. 6I**): *MEF2C*-activated *NPPA* shows upregulation (red) broadly across ventricular and atrial regions, *NR2F2*-activated *TBX5* shows enriched upregulation in specific cardiac zones, while repression regulation of *NKX2-5* -| *MYH11* and *NR2F2* -| *MYL2* displays spatially restricted downregulation (blue) predominantly in ventricular cardiomyocyte-dense regions. Extended spatial mapping of additional regulatory pairs (**Fig. S7C**) shows that *MEF2C*→*NKX2-5* activation is broadly distributed across the heart, *MEF2C*→*TNNT2* activation is strong throughout blood vessels, including pulmonary artery and aorta, *TBX3 -*| *GJA5* repression is localized to conduction-related zones, and *TBX3 -*| *NPPA* repression shows a more diffuse but ventricular-biased spatial pattern. These spatially resolved perturbation maps demonstrate that combining foundation model-based in-silico perturbation with optimal transport spatial mapping can predict not only the direction and magnitude of gene regulatory effects, but also their anatomical localization in 3D tissue architecture.

Together, these results demonstrate that PantheonOS’s intelligent routing system lowers the barrier to leveraging the rapidly expanding ecosystem of virtual cell or single-cell foundation models by abstracting model selection, environment configuration, and execution behind a unified agentic interface. Through the vcRouter, researchers can seamlessly orchestrate diverse foundation models, ranging from scGPT for perturbation effect quantification to Tabula for gene regulatory network inference, and integrate their predictions with spatial data to derive biologically contextualized insights. More broadly, this work highlights how the tight coupling of agentic systems with virtual cell models provides a foundation for autonomous and predictive virtual cell frameworks, enabling systematic forecasting of cellular responses to genetic perturbations within spatial and developmental contexts.

## Discussion

We stand at the threshold of a technological singularity in scientific discovery, catalyzed by the emergence of self-improving artificial intelligence systems capable of accelerating human knowledge generation. To harness this transformative potential, we introduced PantheonOS, an evolvable multi-agent operating system designed to scale and accelerate data science. PantheonOS addresses the emergent bottleneck of single cell and spatial data analysis through four foundational innovations: (1) a highly extensible, general-purpose abstract framework; (2) a instantiation for end-to-end single-cell and spatial genomics data analyses; (3) the Pantheon-Evolve module for algorithmic innovation; (4) and an intelligent virtual cell router for adaptive model selection.

### Reconciling Generalizability with Domain Specificity

A persistent dilemma in AI agent is the trade-off between flexible, general-purpose tools and highly specialized, domain-specific applications. In phase one of our development, we resolved this by building PantheonOS from the ground up as an abstract framework that can be instantiated for various domains with deep expertises. By integrating an intuitive graphical user interface (GUI), a robust command-line interface (CLI) for power users, collaborative Slack integration, embedded Jupyter notebook intelligence, and a community App Store, the system remains exceptionally adaptable. This infrastructure allows PantheonOS to seamlessly scale beyond biology to general data sciences, while still providing deep domain expertise when instantiated for genomics.

### Agent-Driven Biological Discovery at Scale

In phase two, we deployed our novel multi-agent architectures, MTP and ACE, which demonstrated performance superior to current state-of-the-art agent systems, though we acknowledge the need for continued systematic benchmarking. Leveraging PantheonOS as a virtual research assistant, we generated and analyzed a novel OpenST-based 3D spatial transcriptomics dataset of early mouse embryogenesis (E6.0), and use it to analyze the data to autonomously reveal spatially segregated *Cer1* and *Nodal* domains within the epiblast, successfully demonstrating the active paracrine inhibition of *Nodal* signaling by *Cer1*. Furthermore, PantheonOS orchestrated the large-scale integration of single-cell multi-omics data from the human fetal heart with whole-organ 3D MERFISH+ maps at post-conception week 12. This analysis uncovered complex spatial gene expression programs and intercellular communication mechanisms critical to understanding cardiac disease ontogeny.

### Agentic Code Evolution for Algorithmic Innovation

Crucial to this work is the self-improving design of Pantheon-Evolve that maximizes exploration and exploitation (phase three). Moving beyond mere execution, Pantheon-Evolve is capable of recursively evolving and optimizing both nascent and heavily optimized state-of-the-art algorithms. We demonstrated its capacity to refine single-cell batch correction methods, e.g. Harmony, Scanorama, and BBKNN, while concurrently improving our newly designed reinforcement learning-based panel design algorithms. This underscores the PantheonOS potential to iteratively evolve any computational methods, thereby accelerating the pace of algorithmic innovation.

### Intelligent Virtual Cell Model Routing

In phase four, we further expanded PantheonOS by establishing an intelligent single-cell foundation model router. This router unifies major virtual cell models, adaptively selecting the optimal framework to successfully recover core cardiogenic regulatory networks and predicting genetic perturbation effects directly within their native three-dimensional spatial context.

As large language models (LLMs) and agentic frameworks continue to master long-horizon tasks and self-evolution, we foresee a fundamental restructuring of the scientific process. PantheonOS provides a blueprint for a future where AI agents collaborate seamlessly with human researchers to automate the entire scientific lifecycle, from hypothesis generation, method development, and code implementation to execution and evolution, result interpretation, and verifiable manuscript drafting.

Ultimately, self-evolving agents possess the capacity to amplify research at an unprecedented scale, driving us toward a scientific singularity where the most intractable challenges in science, particularly genomics and medicine, can be rapidly resolved. PantheonOS is publicly deployed at PantheonOS.stanford.edu, where we invite the global scientific community to utilize, extend, and collectively contribute to this journey of automated, self-evolvable scientific discovery.

## Code availability

The Pantheon framework is available as open-source software at https://github.com/aristoteleo/pantheon-agents. We also released the PantheonOS GUI, available at https://PantheonOS.stanford.edu. All relevant code, jupyter notebooks, prompts, conversational sessions generated or evolved by PantheonOS will be released at https://github.com/aristoteleo/pantheonos-reproducibility. Note that PantheonOS is a research project, please use it with caution for sensitive datasets.

## Data availability

All data used and produced by this work will be released at https://github.com/aristoteleo/pantheonos-reproducibility.

## Acknowledgements

We thank all Qiu lab members for brainstorming and technical discussion of this project. We would like to thank Lu Zhang for providing valuable feedback on platform trials and for offering constructive suggestions for GUI improvements. We also appreciate Ouyang Wei for supportive discussions on the distributed framework and the extensibility of the Foundation Model. We are grateful to Jianhui Lin for assisting with the Tabula model integration. We are grateful to Cencan Xing for guidance on the mechanisms underlying embryonic anterior–posterior axis formation. We are thankful to Kexi Huang for discussing Biomni and sharing insights on AI agents.

X. Q. acknowledged funding support from the Pantas And Ting Sutardja Foundation, the Wu Tsai Neurosciences Institute Big Ideas in Neuroscience Program. We acknowledge funding support from the BASE Research Initiative (to J.M.E. and X.Q), the Applebaum Foundation (to J.M.E.), and Additional Ventures (to J.M.E. and W.G.). We thank Lenovo Infrastructure Solution Group for computing support.

## Author Contributions

X. Q. conceived the project. W. X led the overall development of PantheonOS. E.P. led the development of RL-augmented gene panel design. Q. Zho. led the PantheonOS-GUI development and optimization. Z.Ze. led the development of Pantheon-CLI and contributed to mouse embryo data analyses. C.Z. led the Pantheon-Evolve on RL-based gene panel design. X. W. led the development of the PantheonOS router. Y.F. led to the mouse embryo data analyses. M. C., C. H. contributed to spatial transcriptomics experiment of mouse embryo, D.O. contributed to mouse embryo array sample collection and embedding. J. D. provide technical guidance during the early stage of this project. Z. Zh. and Y.Y. contributed to benchmarking of PantheonOS agent platform. J.M.E. contributed to in-depth discussion of the human fetal heart multi-omics and spatial transcriptomics integration. All authors contributed to the development and/or writing of the manuscript.

## Methods

### Key terms

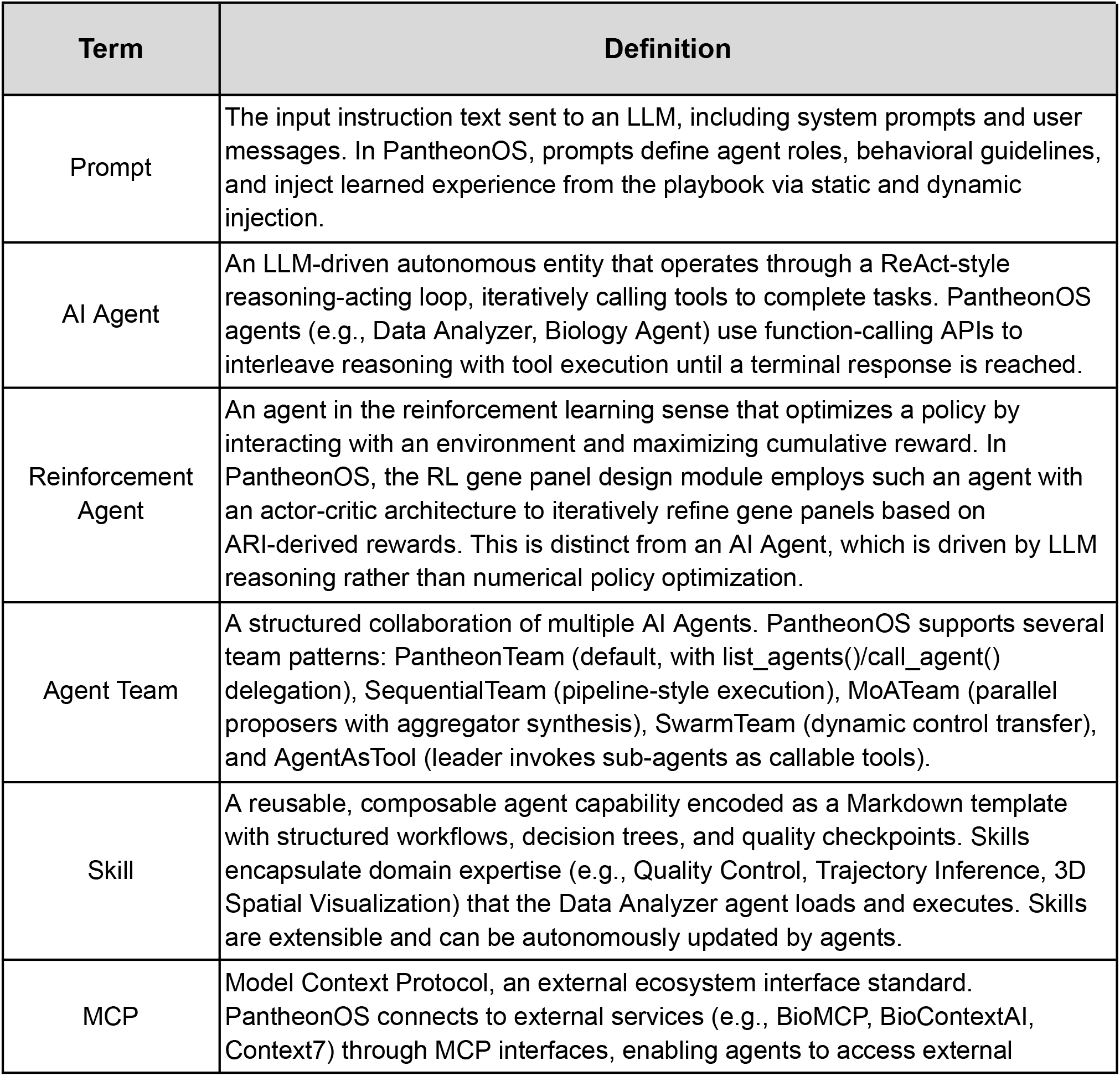
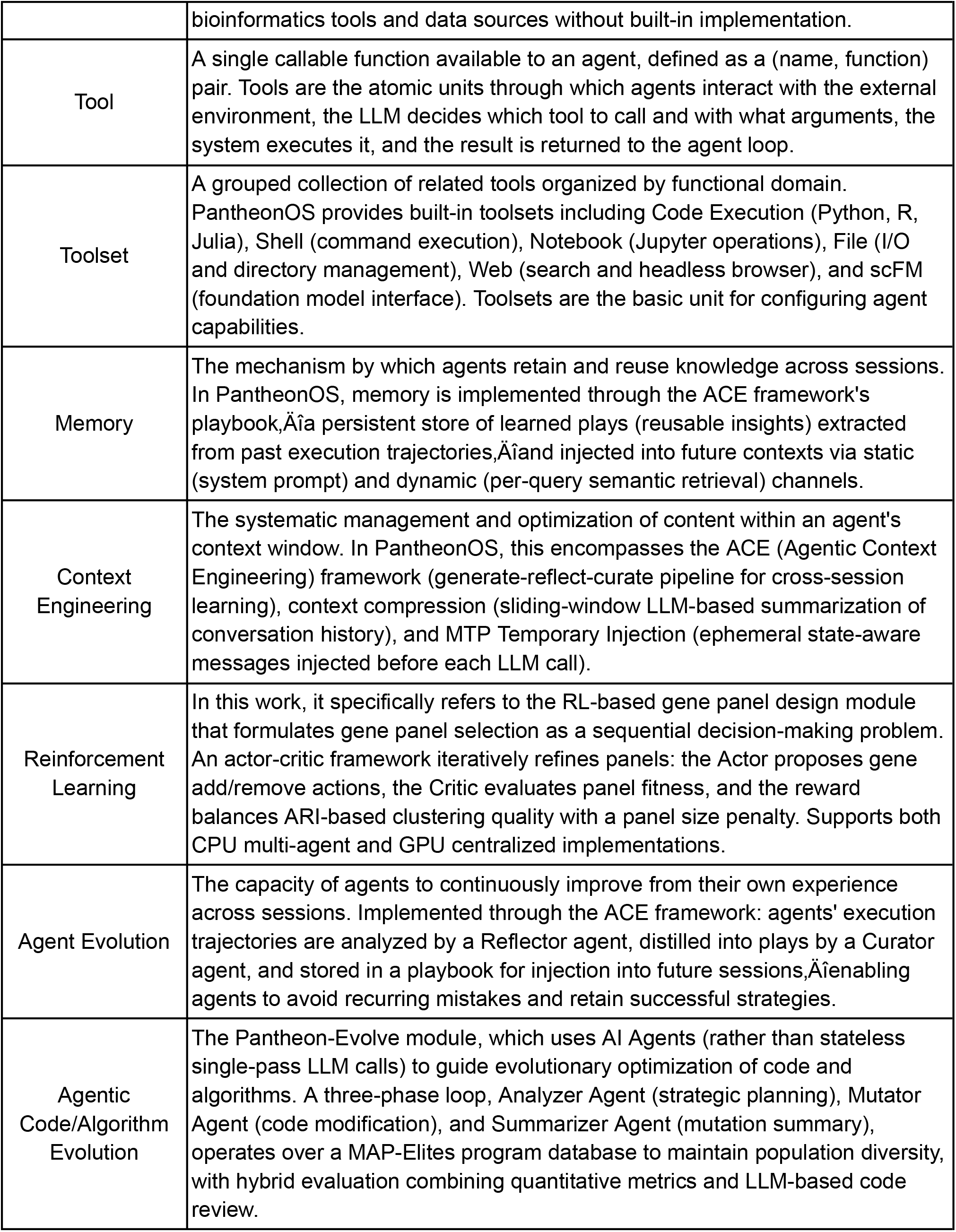

### PantheonOS Architecture and Implementation

#### Design objectives and architecture overview

PantheonOS is a general-purpose, evolvable distributed multi-agent framework to advance data science, with an initial focus in single cell and spatial genomics data analyses. Although PantheonOS is not a real operation system, we choose to name it to include OS because it has a hierarchical abstract design and an extensible ecosystem, is general-purpose and highly modularized. In PantheonOS, we purposely designed four abstract layers to ensure its generality and composability. This design, as detailed in the following, intrinsically enables adaptation to diverse application scenarios:

#### Layer 1 - Infrastructure layer

This is the foundation layer for PantheonOS that provides a universal LLM interface and a privacy preserving distributed communication framework. In the following, we will describe the two integrative infrastructure components, LLM interface and distributed communication, in more detail.

##### LLM Interface

PantheonOS leverages the unified LiteLLM (https://www.litellm.ai/) based API interface to support 100+ LLMs across different providers (OpenAI, Anthropic, Google, etc.) and open source models, e.g., DeepSeek, Qwen, GPT-OSS, and GLM. This interface allows seamless switching between commercial and open-source models. For the open source model, it is connected through the Ollama interface. PantheonOS provides three configurable predefined model invocation chains (High/Normal/Low), tiered by model cost and quality; the system automatically selects the appropriate tier based on the user’s API keys and the agent’s requirements. The chain includes built-in model retry, and fallback: each model is retried up to three times, and on repeated failure the system switches to the next model in the chain until a call succeeds; if the chain is exhausted, it returns an error if no local model can be used. To better manage the user usage, the interface provides mechanisms to track the API cost and token usages.

##### Distributed communication infrastructure

PantheonOS’s distributed communication is based on NATS (https://nats.io/), a lightweight, high-performance messaging system. NATS is an industrial level distributed messaging system that supports low-latency, high-concurrency message distribution. With this infrastructure, PantheonOS implements **Endpoint** and **Chatroom Server** to allow privacy preserving communication between them from different devices and enable flexible deployment across environments and devices. The Endpoint provides an endpoint that connects to a specific computing environment (such as a physical server or a virtualized container), offering independent file access and management tool invocation. It also supports local, remote, and containerized execution. The Chatroom Server, on the other hand, is a centralized conversation history manager. It has the following four features: (1) Agent memory and context persistence; (2) Multi-agent coordination and message routing; (3) Template-based team loading from Markdown configurations

#### Layer 2 - Functional layer

PantheonOS has rich builtin ToolSets, Agents, Team structures and skill systems.

##### Builtin toolsets

PantheonOS supports a variety of builtin toolsets, including (1) code execution: Python, R, Julia; (2) Shell: Command execution, environment management; (3) Notebook Toolset: Jupyter notebook creation, cell execution; (4) File Toolset: file I/O, directory management; (5) Web Toolset: duckduckgo or google (via the scracper API) for internet search, playwright headless browser, for web scraping; (6) scFM: Interface for single-cell foundation models or FM and others.

##### Builtin agents

PantheonOS supports a variety of builtin toolsets, including (1) Data analysis agent: Data analysis, visualization; (2) Biology agent: Domain specific biological interpretation; (3) Browser agent: Web search, literature retrieval, documentation lookup; (4) Coding agent: Code generation; (5) Evolution agent: Algorithm optimization via LLM-guided or more precisely, Agent-guided mutation (see section *Pantheon-Evolve framework: Agentic Code Evolution*); (6) Report agent: Automated report and documentation generation and others.

##### Builtin team structures

PantheonOS includes a variety of agent team structures or agent communication patterns: (1) the default **PantheonTeam**, recommended for most multi-agent workflows, comes with a set of methods, including (a) *list_agents()* for discovering other agents and their capabilities; (b) *call_agent(agent_name, instruction)* for delegating tasks to other agents; (c) *transfer_to_agent(agent_name)* for hands-off control to another agent (optional turn-on); (d) plugin system consists of *LearningPlugin* for learning from interactions and *CompressionPlugin* for context compression. Furthermore, PantheonOS’ team structures support nested delegation with depth control and context execution tracking and metadata management. (2) **SequentialTeam**, recommended for pipeline-style data processing with predefined execution order and configurable connection prompts between agents. (3) **MoATeam (Mixture of Agents)**, built based on MoA^16^ and Self-MoA^48^, allows multiple proposer agents generate solutions in parallel, then aggregator synthesizes the best answer with diverse perspectives and robust decision making. (4) **SwarmTeam**, similar to OpenAI Swarm, in which agents can dynamically transfer control and all context to each other to allow dynamic workflow and adaptive task routing. (5) **AgentAsTool Team**, in which the leader agent treats sub-agents as callable tools or functions. Thus, it has a hierarchical control with a clear leader-worker pattern.

Lastly, PantheonOS comes with a **Skill system** that enables reusable, composable agent capabilities.

In conclusion, PantheonOS’s functional layer provides a rich set of out-of-the-box components for building upper-layer applications.

#### Layer 3 - Interface layer

PantheonOS has a flexible interface layer that consists of multiple interaction modes, including command line interface or CLI, Pantheon-JupyterLab, graph user interface or GUI and Slack integration and App store, Pantheon Store for community sharing.

##### GUI (Web-based)

The overall design of PantheonOS’ GUI includes the two column windows, a built-in Jupyter Notebook for interactive analysis, real-time visualization of agent activities and the interface for knowledge database (papers or literature, etc) and that for the Pantheon-Evolve. We used Vue (https://vuejs.org/) to build the frontend for the GUI.

##### CLI (Command-line interface)

Pantheon’s CLI interface is terminal native and built on fire (https://github.com/google/python-fire), prompt_toolkit (https://github.com/prompt-toolkit/python-prompt-toolkit) and rich (https://github.com/Textualize/rich) for beautiful and powerful UI. It has the following built-in commands, including *!cmd, /view <file>, /edit <file>, /history*, /token, /status and many more. PantheonOS’s CLI has an extensible command handler pattern that allows custom commands.

##### JupyterLab integration

this plugin for JupyterLab enables seamless CLI integration, step-by-step execution and real-time visualization for data analyses tasks, bridging natural language, code, and data in one unified notebook experience.

##### Pantheon Store

Just like Apple’s app store, the Pantheon Store is a community-driven marketplace for sharing components, including Agents, Teams, Skills, Workflows, Knowledge bases, applications. The app store allows one-click installation and updates. The frontend of the Store is built with Vue, and the metadata is stored on GitHub.

In summary, the flexible interface layer lowers barriers to use, promotes community collaboration.

#### Layer 4 - Application layer

The application layer in PantheonOS is highly general. Any specific application can be easily assembled with configured files. This universal design thus allows rapid building of any domain-specific Agentic applications. This paper primarily describes specific use cases for single-cell and spatial transcriptomics analysis, detailed descriptions are provided in the next section.

Overall, PantheonOS’ four layer design ensures the encapsulation and abstraction of infrastructure, function, interface and application, forming a hierarchical pyramid where the lower layer establishes the foundations for the upper layers. While this abstract design is crucial for any data science research, we focus on instantalizing it for single cell and spatial genomics data analyses, an increasingly important area of research in biology.

### Pantheon Omics-Expert-Team

The Omics-Expert-Team is a domain-specific multi-agent system built on PantheonTeam architecture, consisting of one Leader agent and six specialized sub-agents for automated single-cell and spatial transcriptomics analysis. The six sub-agents (Data Analyzer, Biology, Coding, Evolve, Browser, Report) and the leader agent are divided by working context to minimize token overhead. A typical workflow for the leader agent starts with receiving user queries, followed by analyzing user requirements, designing step-by-step execution plans, confirming with users, dispatching tasks to sub-agents, monitoring progress and handling exceptions and finally, aggregating results. Each subagent has specific roles: (1) Data Analyzer: Executes concrete omics analysis tasks, operates Jupyter Notebook for interactive and reproducible analysis, constructs step-by-step pipelines for data loading, preprocessing, downstream analysis and visualization; (2) Biology Agent: Generates biological hypotheses based on data patterns, provides domain-specific interpretation, suggests meaningful follow-up analyses; (3) Coding Agent: Summarizes and catalogs generated code, builds reusable libraries for common operations, enables code reuse instead of regeneration; (4) Evolve Agent: Interfaces with Pantheon-Evolve module, identifies performance bottlenecks, triggers evolutionary optimization of analysis packages; (5) Browser Agent: Retrieves external information from scientific literature and databases, fetches documentation for analysis decisions; (6) Report Agent: Aggregates workflow results, generates comprehensive reports in HTML/Markdown format or LaTeX/PDF with publication-ready figures. To enable smooth communication between agents, agents discover each other via *list_agents()* and delegate tasks through *call_agent()*, making autonomous decisions about when to invoke other agents based on task requirements. To sum up, this hierarchical yet flexible design enables efficient, transparent, and user-controllable genomics analysis workflows while maintaining specialized expertise across all aspects of the analysis.

Our Omics-Expert-Team has a large variety of builtin skills for single cell and spatial genomics data analyses. The Data Analyzer agent loads and utilizes a collection of builtin skills that encode structured, reusable workflows that tells the analyzer agent how to perform single-cell and spatial transcriptomics analysis reliably. PatheonOS’ builtin skills are extensible and the analyzer agents can autonomously update and expand based on emerging analysis requirements. For general data analysis, PantheonOS support: (1) Parallel computing skills: Multiple optimization strategies including multi-core CPU parallelization: Numba(https://numba.pydata.org/), OpenMP(https://www.openmp.org/); GPU acceleration with RAPIDS(https://rapids.ai/) achieving 10-100x speedup, memory optimization patterns, and Dask(https://www.dask.org/) for out-of-core computing; (2) Environment management skills: Built-in knowledge about environment management related to conda, venv, and Docker; under normal circumstances, using conda for environment management is recommended; (3) Database access: using the skills of gget^49^, iSeq^50^, and cellxgene^51^ to access a variety of databases, including SRA, GEO, Ensembl, UniProt^52^, UCSC^53^, Enrichr^53^, and CZI cellxgene^51^. More specifically, to automate single cell and spatial transcriptomic data upstream Analysis, we support over 100 different upstream data analysis pipelines by integrating nf-core^54^, covering fields such as single-cell/bulk RNA-Seq, ATAC-Seq, epigenomics, spatial transcriptomics, 3D genomics, and more. To automate marker gene selection for targeted panels of spatial profiling, such as MERFISH^55^, 10x Xenium^56^, the Omics-Expert-Team has gene-panel design skill that can be used for optimizing gene combinations based on cell type discriminability, relevance to specific biological questions. To facilitate routine single cell analyses, we incorporate the following core Single Cell Analysis Skills: (1) Quality Control: Three-phase workflow including ambient RNA assessment by SoupX^57^ or DecontX^58^, doublet detection by Scrublet^59^ or DoubletFinder^60^, followed by unified cell filtering, normalization to 10,000 counts per cell, and highly variable gene identification with Scanpy^20^; (2) Cell Type Annotation: Support multiple annotation approaches including marker gene-based methods (selecting markers based on differential expression and filtering of LogFC > 0.5), reference tools-based method (CellTypist^61^, scANVI^62^), and automated LLM-based annotation with confidence scoring; (3) Trajectory Inference: Pseudotime based analysis using Diffusion Pseudotime (DPT^63^), PAGA for complex topologies, RNA Velocity (scVelo^64^) for directional dynamics. Additionally, we also incorporate 13 essential skills described in the “Single-cell Best Practices” book, covering differential expression analysis, multi-omics analysis, and more. To enable advanced spatial analysis, we also implement Skills for (1) Single-Cell to Spatial data Mapping: Optimal transport-based mapping using MOSCOT^41^ for gene imputation, cell type label transfer, and handling large datasets (>100k cells) with subsampling and confidence assessment; (2) 3D Spatial Visualization: Interactive visualization using PyVista^65^ for the expression of different genes of 3D tissue or embryo, cell type coloring with custom palettes, rotating GIF animation generation, supporting millions of spatial coordinates. We also support the single cell foundation model workflows. PantheonOS’s automated model selection is based on data compatibility (gene ID format, species) and data analyses request, supporting 22 models including UCE^66^, scGPT^43^, Geneformer^67,68^, and Tabula^45,46^ foundation model. See **Tables S2** for the full list. To allow extensibility and flexibility of the skill system, in PanthenOS, new skills can be added by creating template files following the established schema, and the Coding Agent can propose new skills updates based on analysis patterns. This extensible skill system enables the Data Analyzer to perform sophisticated analyses while maintaining reproducibility, with the team capable of autonomously evolving its capabilities through collaborative skill development and refinement.

### Runtime Mechanisms of Pantheon Agents

PantheonOS’ agents have five different runtime mechanisms: **(1) The basic agentic loop**, a ReAct-style iteration interleaving reasoning and tool execution; **(2) Inter-agent transfer**, including two communication modes: handoff and delegation (agent-as-tool invocation with isolated context); **(3) The Modal Task Protocol (MTP)**, imposing phased workflow via Temporary Injection, mode transitions, and interruption; **(4) Context compression**, an LLM-based sliding-window summarization that replaces old messages with structured checkpoints when token usage exceeds a threshold; **(5) Agentic Context Engineering (ACE)**, a cross-session learning system extracting reusable plays from agent trajectories via a reflector–curator–playbook pipeline. The following explain each mechanism in more detail.

#### The Basic Agentic Loop

The basic agentic loop is the most fundamental operational mode for agents in Pantheon. It is a ReAct-style^1^ reasoning-acting loop realized via the function-calling API. We firstly introduce some key definitions: (1) **Message**: A message, denoted as *m* = (*ρ,c*), includes both the role *ρ* ∈ {*system,user,assistant,tool*} and content *c*. We define ℳ as the set of all messages; (2) **Tool** denoted as *τ* = (*n,f*), is the pair with as the function name and f : 𝒜 → ℛ as a (possibly async, side-effecting) function. We define 𝒯 as the tool registry or a dictionary of tools the agent can call; (3) **Tool Call**: A triple of unique identifier, tool name, argument payload, denoted as *k* = (*id,n,a*); (4) **LLM Call**, denoted as LLM : ℳ * × 𝒫 (𝒯) → ℳ ∪ (ℳ × 𝒦 ^+^) where # means the Cartesian product to create an ordered pairs, not multiplication. Given history *H* and tools *T*, returns either a terminal assistant message *m*_*a*_, or (*m*_*a*_,*K*) with tool calls *K* = [*k*_1_, …, *k*_*k*_], see **Algorithm 1** below.

##### Algorithm 1

Basic Agentic Loop

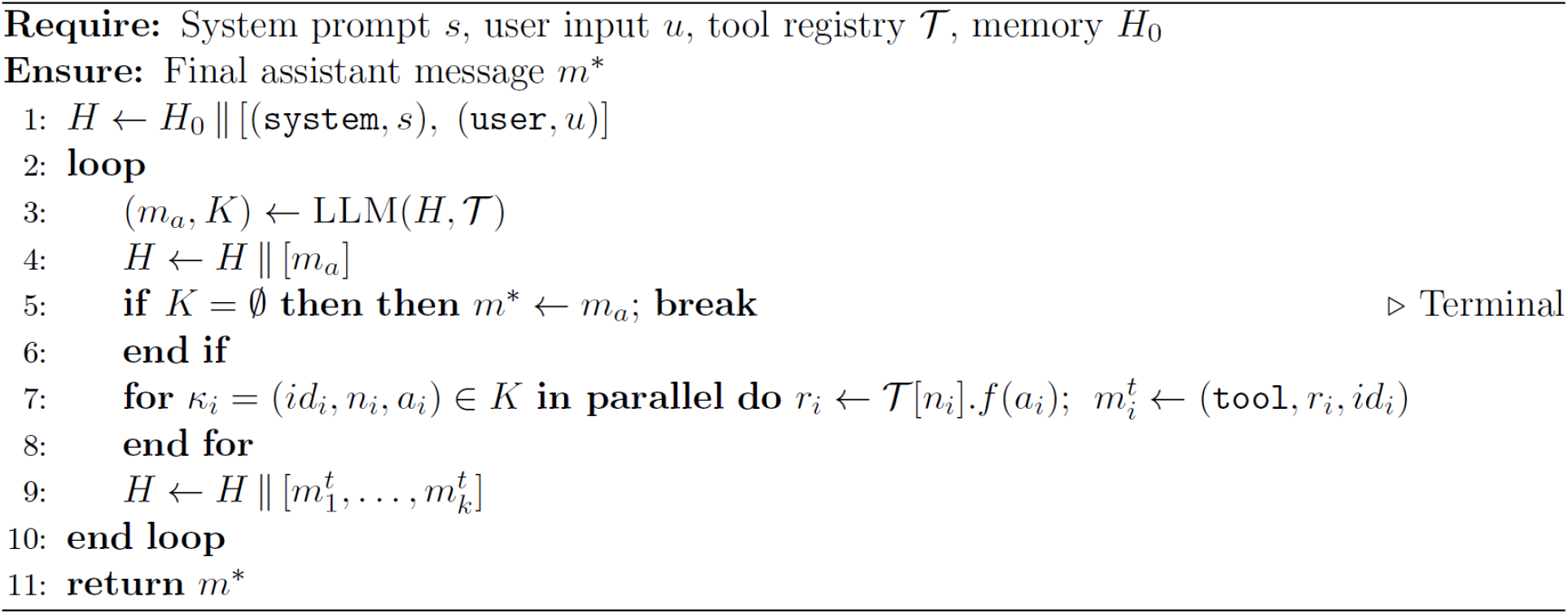

##### We note two special cases in the Algorithm 1 above: (1) Termination

The loop terminates on a no-tool-call response; a max-turns guard *N* is imposed in practices; **Transfer**: If a tool returns a transfer sentinel, the loop yields an *AgentTransfer* record and the orchestrator switches to the target agent (see the next subsection *Inter-Agent Transfer*). It is also worth noting that, in **Algorithm 1**, for **parallelism**, all tool calls *K* within one LLM turn execute concurrently.

#### Inter-Agent Transfer

When multiple agents coexist under a team orchestrator, two communication modes, namely handoff and delegation, are available. Before introducing them in detail, we will start with some definitions (1) **Agent Pool**: A team maintains an agent pool 𝒜 = [*a*_1_, …, *a*_*n*_] with one active agent *a*^*^ ∈ 𝒜 at any time; (2) **Agent Transfer Record**: 𝒳 = (*a*_*s*_,*a*_*t*_,*H, 𝒳,i*_0_) : source agent, target agent, conversation history, context variables 𝒳, and initial message offset *i*_0_ (marking where the transferred context begins).

#### Mode 1: Handoff

Inspired by the Handoffs pattern introduced in the OpenAI Agents SDK (https://openai.github.io/openai-agents-python/handoffs/), in PantheonOS, a handoff transfers control entirely from one agent to another, carrying the full conversation history. Handoff works through the following mechanism: the orchestrator registers a tool *trans*_*to*_*agent*(*a*_*t*_) for each reachable *a*_*t*_ ∈ 𝒜\ {*a*^*^}. When invoked, the tool returns an agent reference *a*_*t*_. The basic loop (**Algorithm 1**) detects this as a transfer sentinel: it constructs 𝒳 = (*a*^*^,*a*_*t*,_ *H* [*i*_0_ :], 𝒳 |*H*|) and yields to the orchestrator. Then the team orchestrator runs a transfer-detecting loop, as shown in the following:

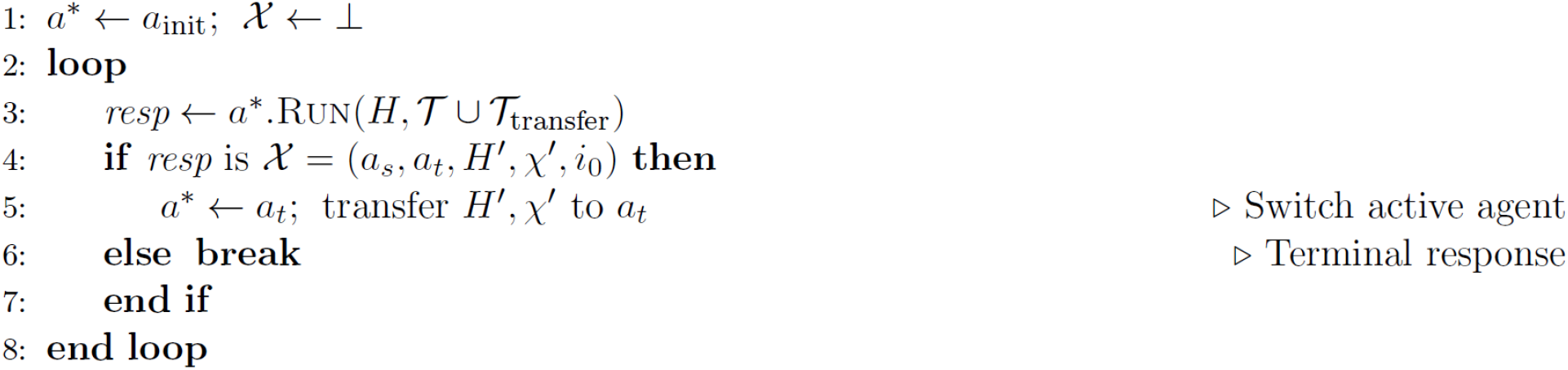

Note that the target agent *a*_*t*_ receives *H*^′^ = *H* [*i*_0_ :], i.e., the full message sequence from the start of the conversation. This preserves conversational continuity across handoffs.

#### Mode 2: Delegation (Agent-as-Tool)

For the delegation mode, it happens when an agent invokes another agent as a tool call: the child executes in an isolated context and returns a result to the parent. The delegation mechanism works as follows. The orchestrator firstly registers *call*_*agent*(*ac,task*, …), which:

1. creates an isolated execution context *ϵ* = *NewContext* (*a*^*^,*a*_*c*_) with a fresh context ID;
2. optionally compresses the parent’s history into a summary message;
3. runs *a*_*c*_.*Run* (*task, ϵ*) ;
4. returns *a*_*c*_ ‘s final response as a tool result to the parent.

Unlike handoff, delegation does not transfer full history, thus ensuring context isolation. The child agent *a*_*c*_ operates with its own memory scope, sees only the task description (plus optional summary), and cannot modify the parent’s state. After completion, control returns to *a*^*^ automatically. Each delegation appends *a*_*c*_ to a chain path *P* = [*a*^*^,…, *a*_*c*_].

Before invoking *call*_*agent*, the orchestrator checks *a*_*c*_ ∉ *P* ; if violated, the call is rejected to prevent infinite delegation cycles.

#### The Modal Task Protocol (MTP)

The basic agentic loop is task-unaware: it iterates until the LLM stops emitting tool calls, with no notion of what phase of work the agent is in, no mechanism to communicate progress to the user about what is currently running, and no way for the agent to voluntarily yield control. Inspired by the artifact-driven, plan-execute-verify workflow of Google Antigravity (https://antigravity.google/blog/introducing-google-antigravity), MTP is an optional intra-session supervisory layer that addresses these limitations by introducing four concepts, namely, **modes, artifacts, Temporary Injection**, and **interruption**. With these concepts, MTP structures different tasks into explicit stages, adapts guidance based on the current stage, and reports progress to the user, interrupting execution when human review is required. MTP does not alter the core loop logic, instead it wraps it by (a) injecting a synthetic message before each LLM call and (b) intercepting specific tool calls to update the protocol state. MTP is activated for each agent by adding *task_toolset* to the agent’s toolsets list in its template configuration. Additionally, it also registers *task_boundary* and *notify_user* tools, and enables the Temporary Injection hook in the agent loop. Agents without this toolset operate with the unmodified basic loop.

In the following, we will introduce the algorithm for MTP, starting with defining key concepts related to state space, state transition, and temporary Injection.

#### State Space

- **Mode**: defined as the system’s current way of thinking or working. A *μ* mode belongs to one of three high-level phase Φ = {*plan,Execute, Verify* }. A function Φ: *ModelLabel* → Φ maps mode labels to phases. More formally,

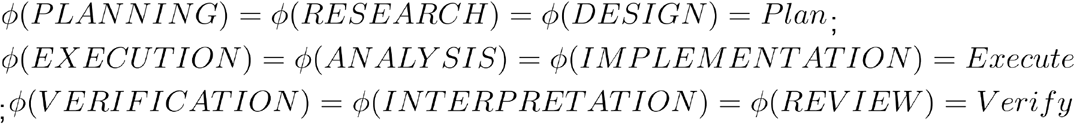
- **Task**: defined as a structured record with four fields, that is a tuple *θ* = (*name, μ, status, summary*) where *μ* is the current mode and others are self-explannatory.
- **Artifact**: Artifacts are **durable outputs** that let the system remember, inspect, and resume work. The function *ρ* will assign the file *α* to a semantic role *ρ* (*α*) ∈ { *task, plan, summary, tracker* } determined by filename.
- **Protocol State**: The **protocol state** Σ is a big container that stores **everything the system needs to know** at a given moment. Σ = (*θ*^?^, 𝒜_*u*_, λ, Δ, *n*_*b*_,,*n*_*u*_,*t*) with active task *θ*^?^ ∈ {*θ*, ⊥}, created artifacts 𝒜_*c*_, last-access map λ: *Paths* → ℕ which maps each file path to how recently it was accessed, per-task modifications, Δ: *Roles* → 𝒫 (*Paths*) in which the Δ records, for each artifact role, the power set or all possible set of file paths modified by the current task, tool counters *n*_*b*_,*n*_*u*_ where *n*_*b*_,*n*_*u*_ indicates blocked and unblocked counter that counts the failed or successful tool invocations, and global step *t*.

#### State Transitions

State transitions move the system from one state to another during execution which are enabled by three transition operators, namely *TaskBoundary, NotifyUser, ToolExeuted*, each triggered by a specific tool call:

- *TaskBoundary* (*name, μ, status, summary*): Sets *θ*^′^= (*name, μ, status, summary*); resets Δ^′^ = ∅ if the task name changed, 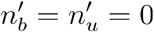. This operator will let the agent create or update a task.
- *NotifyUser* (*paths, blocked, msg*) : Sets 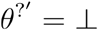, records *pending* = *paths. blocked*: emit Interruption. This operator will let the agent stop the task, send a message to the user, optionally interrupting the loop.
- *ToolExecuted* (*calls, results*): Increments by *t, n*_*p*_, *n*_*u*_ by |*calls*|; for each artifact access *α*, sets λ ^′^ (*α*) = *t*^′^ (update the last access timestamp of the file to the current step) and Δ^′^+ = *α*. This operator will let the agent execute the functional call.

#### Temporary Injection

Before each LLM call, the system adds a temporary or ephemeral message summarizing the current system state that is not stored afterward. The injection function converts the current state into one message, that is, *E* : ∑ → ℳ which generates *m*_*e*_ =(*user,Render* (∑)) . The LLM sees *H*_llm_ = *H* ||[*E*(∑)]; persistent *H* is unchanged.

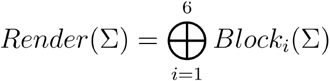

where ⊕ is the concatenation of text blocks and blocks are conditionally included:

| # | Block | Condition |
| --- | --- | --- |
| 1 | ARTIFACTLIST | Always |
| 2a | ACTIVETASKREMINDER | $\theta^? \neq \perp$ |
| 2b | NOTASKREMINDER | $\theta^? = \perp$ |
| 3 | EXCESSIVETOOLSWARNING | $\theta^? = \perp \wedge n_b \geq 5$ |
| 4 | PLANMODIFIEDALERT | $\phi(\mu) = \text{PLAN} \wedge \Delta[\text{plan}] \neq \emptyset$ |
| 5 | PENDINGREVIEWALERT | $\theta^? = \perp \wedge \text{pending} \neq \emptyset$ |
| 6 | STALEARTIFACTALERT | $\exists \alpha : t - \lambda(\alpha) > 10$ |

#### Modified Agent Loop under MTP

With the Modal Task Protocol, the basic agentic loop will be updated into **Algorithm 2** with inclusion of *TaskBoundary, NotifyUser*, and *ToolExeuted* after all tools has been executed in parallel:

##### Algorithm 2

Agentic Loop with MTP

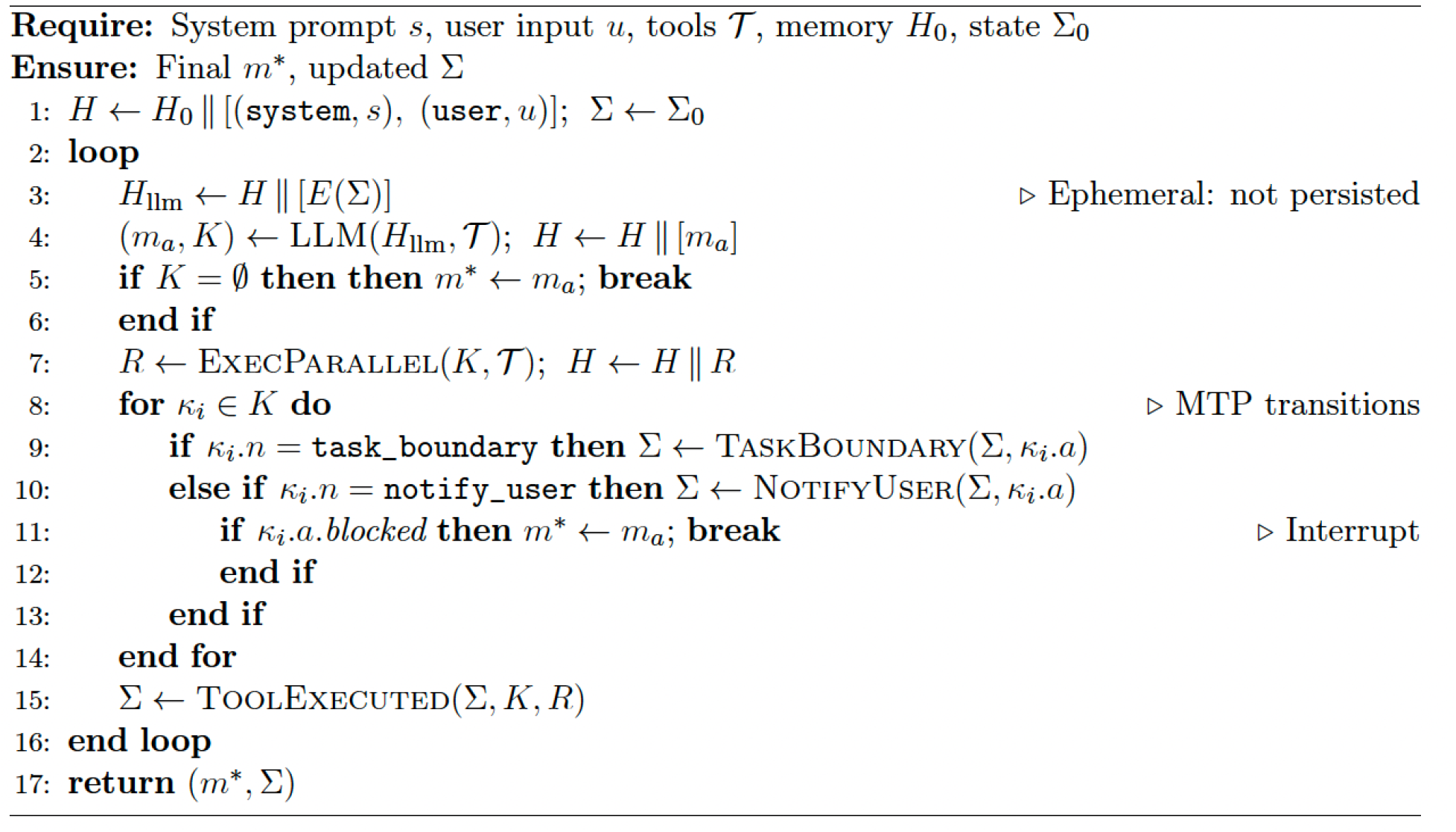

Note that in **Algorithm 2**, *k*_*i*.*n*,_ *k*_*i*.*a*_, *k*_*i*.*a*.*blocked*_ denotes tool name, tool argument, and the boolean tag about whether the tool is blocked.

Several properties follow from this design. The ephemeral message (*E*(∑)) is exposed to the language model at step (*t*) but is not recorded in the persistent history (*H*); instead, it is deterministically regenerated from the updated state (∑) at step (t+1). As a result, the protocol introduces no persistent overhead to the conversation history while preserving state-aware guidance at each iteration. Mode-guided behavior is implemented as a soft constraint enforced entirely through prompting, rather than through hard-coded control logic. The ephemeral injection reinforces expected behavioral patterns without explicitly prohibiting actions. The `NotifyUser` transition with `blocked = true` implements a cooperative yield mechanism, analogous to cooperative multitasking, in which the agent voluntarily returns control to the user. Finally, the agent state (∑) is serialized to disk after each transition, enabling robust recovery across process interruptions or restarts.

#### Context Compression

As the history *H* grows, total token count may approach the model’s context window limit. Thus, we use context compression to replace old messages with a structured LLM-generated summary, implementing a non-destructive sliding window over *H*.

#### Trigger Condition

After each assistant response, a token-counting pass populates metadata (*T*_used_, *T*_max_) on the last assistant message. Compression is triggered when the usage ratio exceeds a threshold:

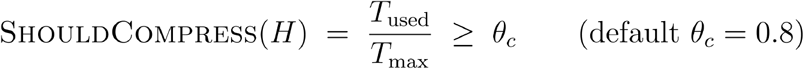

subject to a backoff rule: after a failed attempt (e.g. a failed context summarization trial), at least *n*_retry_ new messages (default 10) must arrive before retrying.

#### Compression Range

Let *j* be the index of the last compression-role message in *H* (or − 1 if none), and *p* the number of recent messages to preserve (default 5). The compression range is:

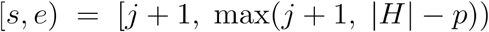

Messages *H* [*s*:*e*] are compressed; *H* [*e*:] remain untouched.

#### Message Formatting

The message formatting pipeline before agent summarization includes four steps: bracket format, per-value argument truncation, tool output truncation and file extraction. Each step reduces **token cost, noise**, and **failure risk**, while preserving **semantic structure**. Before LLM summarization, messages in the compression range are linearized into a bracket-format text representation: (1) **Bracket format**: Each message is rendered as a tagged block: [USER], [ASSISTANT], [TOOL_CALL: name] id=*id*, [TOOL_RESULT] id=*id*, or [PREVIOUS_SUMMARY] (for prior checkpoints). Similarly, the rest three steps are described in the following:

- **Per-value argument truncation**: Tool call arguments are parsed as JSON; each value *v* exceeding *ℓ*_*a*_ characters is truncated to *v*[:*ℓ*_*a*_], preserving short parameters (e.g. *file paths*) intact.
- **Tool output truncation**: Two strategies, selected by a *use_smart_truncate* flag: *Simple mode*: Truncate the string representation to *ℓ*_*o*_ characters. *Smart mode*: Operates on the raw structured result (before stringification). For JSON dicts: (1) filter base64 content, (2) if within *ℓ*_*o*_, return compact JSON, (3) otherwise, apply field-level recursive truncation: allocate budget *ℓ*_*o*_ / |fields| per field, recurse up to depth 2, preserving JSON types. Oversized content is saved to a file; a preview with the file path is returned.
- **File extraction**: During formatting, file paths are extracted from tool arguments: view-type tools (*read_file, grep, glob*, …) populate ℱ_*v*_, edit-type tools (*write_file, update_file*, …) populate ℱ_*e*_.

#### LLM Summarization

The formatted text, together with ℱ_*v*_ and ℱ_*e*_, is passed to a dedicated compression agent *LLMC* (which may use a different model than the main agent to improve the summarization quality). The agent produces a structured summary following a template:

| Section | Content |
| --- | --- |
| OBJECTIVE | User's main goal and sub-goals |
| WORKDONE | Completed items organized by logical category |
| KEYCONTEXT | Decisions, constraints, user preferences |
| FILEINTERACTION | Per-file description of what was viewed/learned and edited/changed |
| NEXTSTEPS | Pending actions and open questions |

#### Checkpoint Insertion and Failure Recovery

Before inserting checkpoints, we will first validate the success of the compression: Let *T*_orig_ be the estimated token count of *H*[*s*:*e*] and *T*_sum_ be the token count of the summary. The compression is accepted only if *T*_sum_ < *T*_orig_ (or if forced); otherwise it is rejected as FailedInflated.

Next, we will ensure non-destructive insertion. That is, on success, a checkpoint message *m*_*c*_ is inserted at position :

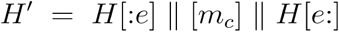

Original messages are not deleted; *m*_*c*_ carries role compression and content wrapped as {{*CHECKPOINT*_*k*_ }}, where is the compression index.

When preparing messages for the LLM, *get_messages(for_llm=True)* finds the last compression message *m*_*c*_, discards everything before it, and converts *m*_*c*_’s role from compression to user.

The LLM thus sees: 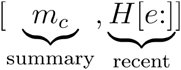 . Therefore it creates a sliding window memory model in which history grows internally while LLM always sees bounded context.

##### Failure Recovery

We summarize the failure-handling logic for checkpoint compression in PantheonOS in the following table.

| Status | Behavior |
| --- | --- |
| COMPRESSED | Success; reset failure counter. |
| FAILEDINFLATED | Summary ≥ original; increment failure counter, retry after $n_{\text{retry}}$ messages. |
| FAILEDERROR | LLM call failed; same backoff as above. |
| SKIPPED | Chunk too small ($< p$ messages); no action. |

Where *p* messages is the minimum number of new messages that must accumulate before the system even tries to compress them.

#### Properties

PantheonOS’ context compression has the following properties: (1) **Non-destructive**: Original messages remain in *H* for audit, cost tracking, and the Agentic Context Engineering (see below) learning pipeline, which reads the full uncompressed history; (2) **Incremental**: Each compression only covers messages since the previous checkpoint, avoiding re-summarization of already-compressed content; (3) **Bounded context**: After compression, the LLM’s effective context is |*m*_*c*_|+ *p* messages, independent of total conversation length; (4) **Dual usage**: The same message formatting and smart truncation utilities are shared between real-time compression and ACE trajectory compression.

#### Agentic Context Engineering (ACE)

A key design goal of PantheonOS is that agents should continuously improve from their own experience: mistakes made in one session should not recur in the next, and successful strategies should be retained and reused. To this end, we adopt the Agentic Context Engineering (ACE) framework^69^ as a cross-session learning layer. While the basic loop handles a single turn and MTP manages intra-session workflow, ACE operates between sessions, that is, extracting reusable knowledge from past trajectories and injecting it into future contexts via a *generate–reflect–curate* pipeline. **The unit of learned knowledge is called a *play* (a concrete, reusable insight extracted from execution experience), and the persistent store is the playbook**.

#### Play Representation

##### Play

A record = (id, *ξ, γ, h, d, n, ω, τ, δ, η*) : id, unique identifier (e.g. *str-00042*); *ξ*, section ∈{ *user_rules, strategies, patterns,workflows, guidlines,mistakes*}; *γ*, content (strategy text, code snippet, or guide); 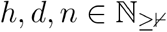, helpful / harmful / neutral tag counters; *ω*, agent scope (*global* or a specific agent name); *τ* ∈ {`user,system`,⊥}, provenance; *δ*, optional short description (≤ 20 words); *η* ∈ {`active,invalid`}, validity.

##### Net Score & Dominant Tag

*Score* (*σ*) = *h* − *d*. Plays are sorted by descending score.

The dominant tag: *DomTag* (*σ*) = *harmful* if ; *d* > *h* ⋀ *d* > *n*; *helpful* if *h* > *d* ⋀ *h* > *n*; if ; *n* > 0 and neither above; ⊥ otherwise.

##### Atomicity

Each learning carries an atomicity score *a* ∈ [0,1] measuring single-responsibility:

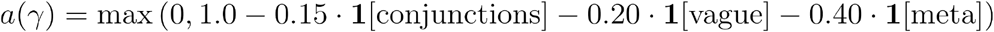

Play type: *atomic* if *a* ≥ 0.85, *systematic* otherwise. An atomic play corresponds to one clear lesson while systematic play is used for higher-level workflow.

##### Playbook

defined as the structured long-term memory that stores everything the system has learned across runs. It is indexed and organized so agents can retrieve and apply them later. A play can be represented as a tuple ℬ = (*S, π, next*) in which play store *S* : 𝒫 → *σ* (the store maps unique IDs to learned knowledge), section index *π* : 𝒮 → 𝒫 (ℐ) (from the store get all the sections where a particular play applies), monotonic ID counter *next*.

#### Trajectory Compression

Before ACE analyzes past experience, the raw history ℳ^*^ is compressed: *Compress* : ℳ^*^ × ℕ^2^ → (*Text* × 𝒫 (ℐ)) produces 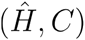, compressed trajectory text and cited play IDs (𝒩 ^2^). The steps for the compression include: (1) truncate tool arguments > *ℓ*_*a*_ chars, (2) truncate tool outputs > *ℓ*_*o*_ chars (both preserving JSON structure), (3) extract play ID references into, (4) linearize into text.

#### The Reflector

The reflector is an LLM agent analyzing trajectories to produce structured diagnostic output. We start with explaining the **reflector output** and **diagnostic priority** while with the following **Algorithm 3**, we define the workflow for the reflector.

##### Reflector Output

ℛ = (analysis, 𝒢, ℒ,*c*) : narrative analysis; play tags 𝒢 ={(*id,g,r*)} with *g* ∈ {*helpful,harmful,neutral*}; extracted learnings ℒ = {(*ξ,γ,a,e*)} with section, content, atomicity, evidence; confidence *c* ∈ [0,1].

##### Diagnostic Priority

The Reflector applies the first matching condition:

| $P$ | Condition | Action |
| --- | --- | --- |
| 0 | USERPREFERENCE | User said “remember”/“always”/“prefer”. Extract as **user_rules**. |
| 1 | SUCCESSCASE | Solved correctly. Extract strategies; tag **helpful**. |
| 2 | STRATEGYMISAPPLICATION | Right strategy, wrong execution. Tag **neutral**. |
| 3 | WRONGSTRATEGY | Wrong strategy. Tag **harmful**. |
| 4 | MISSINGSTRATEGY | No strategy existed. Mark for ADD. |

In the following algorithm, the Reflector takes a past execution trajectory, comprises it, contextualizes it with the current playbook, and asks an LLM to analyze what happened and extract candidate learnings.

###### Algorithm 3

Reflector Analysis

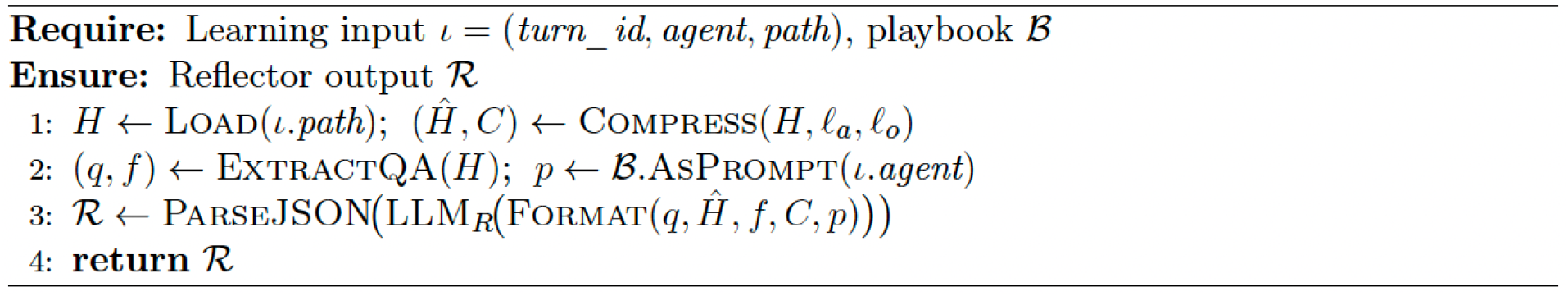

Where *q,f* corresponds to the original user question / task or the final outcome, respectively. In addition, ℬ.AsPrompt(ı.agent) describes the playbook will be converted into prompt form and filtered for the specific agent from a learning record *l*.

#### The Curator

The curator is a separate LLM agent translating reflector output into playbook operations, responsible for changing memory.

The curator can perform four primitive operations: *Add*(*ξ, γ,a, δ, ω*), insert new play from section ([\episilon?]), content (*γ*), atomicity/confidence (*a*), metadata (*σ*), and scope (*ϕ*); *Update* (*id, γ*^′^), replace content of an existing play;

*Tag* (*id*,g), increment feedback counter *g* ∈ {*h,d,n*} ; *Remove* (*id*), soft delete a play by making it invalid (set *η* = *invalid*). User-defined plays (*τ* = *user*) are protected: only Tag is permitted.

To define which action to take first when multiple conditions apply, we provide the following priority table with decision logic:

| $P$ | Condition | Action |
| --- | --- | --- |
| 1 | CRITICALERROR | ADD corrective strategy ($a > 0.85$); TAG harmful. |
| 2 | MISSINGCAPABILITY | ADD atomic play ($a > 0.70$). |
| 3 | STRATEGYREFINEMENT | UPDATE with better content. |
| 4 | CONTRADICTIONRESOLUTION | REMOVE/UPDATE conflicting entries. |
| 5 | SUCCESSREINFORCEMENT | TAG as helpful. |

This ensures safety fixes override optimization, new capabilities precede refinement, reinforcement never masks errors. To ensure presentation of memory bloating, quality control and boundary memory, we will implement **Pre-Add Deduplication, Atomicity Gate** and **Eviction** mechanisms.

##### Pre-Add Deduplication

Before any Add operation, the curator must find the most similar existing play and test semantic equivalence: if *DedupCheck*(*γ*_new_, 𝔅) = *Update*(*id*^*^, *γ′*) if ∃σ^*^ : *Same Meaning(γ*_new_, σ^*^ ) ; otherwise . *Add* (…)

##### Atomicity Gate

Add is accepted if: (a) type is systematic; or (b) type is atomic and *α ≥ α* _min_ = 0.85. Hard rejection below *α*_reject_ = 0.40. Therefore, only **sharp, reusable knowledge** enters memory.

##### Eviction

When section *ξ* reaches capacity 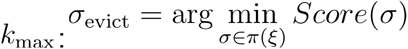, subject to σ_evict_.*d* > σ_evict_.*h* or a play can be evicted only if it has more negative than positive evidence. If no candidate qualifies, the Add is rejected. This ensures memory stays unbounded and high quality plays are preserved.

#### Learning Pipeline

In the following algorithm, the Curator is a policy-constrained LLM agent that safely translates reflective analysis into minimal, high-confidence, non-duplicative updates to the playbook while preserving user intent and bounded memory growth.

##### Algorithm 4

ACE Learning Pipeline

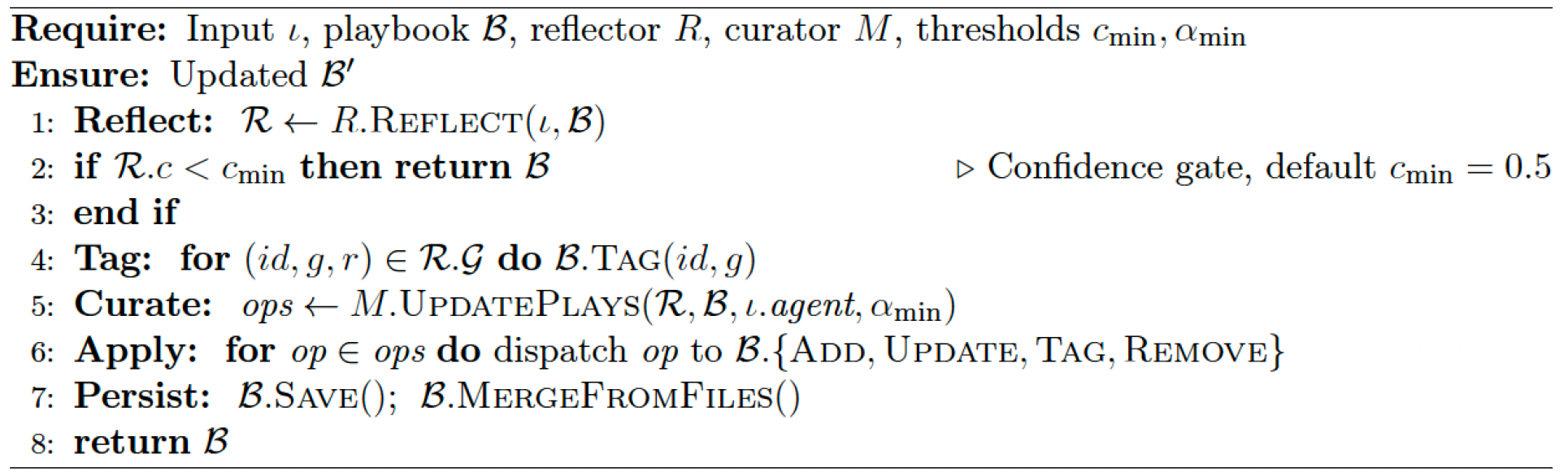

#### Playbook Injection

Two complementary channels, static or dynamic injection, inject plays back into agent contexts.

##### Static Injection

At agent creation, it injects selected plays once, by directly appending to system prompt:

*S′ = s∥ Format Static({σ∣ Is Static (σ) ∧ In Scope (σ, a)})* where

*Is Static(σ) = (σ*.*ξ = user _rules) ∨ (σ. ξ = strategies ∧ σ. τ ∈ {user, ⊥})* and

*In Scope(σ,a) = (σ. ω = global) ∨ (σ. ω = a)*. Persists for the agent’s lifetime.

##### Dynamic Injection

At each user message: (1) semantic search over *𝔅*.*S* with user query top-candidates; (2) filter out *IsStatic* plays; (3) wrap in <EPHEMERAL_PLAYS> tags. Like MTP ephemeral messages, visible for one turn only.

The static and dynamic injection ensures (1) **Disjointness**: *Static Plays ⋂ Dynamic Plays* = ∅, no play is presented through both channels; (2) **User preference protection**: Plays with *τ = user* are always statically injected and cannot be modified/removed by the pipeline (only tagged). They also have the following properties: (1) **Async non-blocking**: The pipeline runs in a background task; the updated playbook becomes available for the next interaction; (2) **Incremental deltas**: Each operation affects one play record, avoiding context collapse from monolithic rewriting; (3) **Dual-LLM separation**: Reflector (“what happened?”) and Curator (“how to change the playbook?”) are distinct agents, preventing conflation of analysis with curation; (4) **Dual gating**: Confidence gate (*c < c*_min_) discards unreliable reflections; atomicity gate (*α < α*_min_) rejects vague learnings; (5) **Grow-and-refine**: Growth via Add; refinement via Update/Remove/eviction; deduplication converts would-be duplicates into updates. Thus, PantheonOS injects learned plays into agent contexts via a dual-channel mechanism: persistent static rules embedded at agent creation and ephemeral dynamic plays retrieved per query, ensuring relevance, stability, and user preference protection.

##### To sum up, in PantheonOS, the reflector answers “What happened?", while the curator answers “how should the playbook change?”

The implementation follows the generate–reflect–curate paradigm of ACE69 with three refinements: (1) the curator is a separate LLM agent rather than non-LLM merge logic; (2) pre-add deduplication replaces post-hoc embedding-based pruning; (3) injection is bifurcated into static (user rules) and dynamic (semantic search) channels.

#### Gene Panel Design Workflow

PantheonOS implements a built-in workflow to efficiently design gene panels for targeted spatial transcriptomics technologies, such as MERFISH, STARMap and others with skill based workflow mechanisms. Specifically, the goal of this workflow is to build a gene panel whose expression maximizes representability of the cell states while at the same time meet criteria sought by a user query, including biological context constraints, e.g., requirement of the inclusion of a specific category of genes, such as cancer signaling pathway and others. In more detail, the gene panel design workflow includes: (1) **Single cell data acquisition**. In this step, PantheonOS automatically retrieves the relevant data from cellxgene database based on user queries, and then have the Agent assess the data to determine whether to perform preprocessing. Once the data is downloaded, Pantheon will automatically define the train, test, and validation dataset, based on the size of the dataset. (2) **Construction of a gene panel balancing cell-type discrimination and biological relevance**.

The gene panel design involves several critical steps, namely (1) Pareto optimal gene panel seed selection, (2) Seed selection enhanced with reinforcement learning, (3) biological prior lookup and context matching, (4) consensus filing, (5) benchmarking, where the first two involves two different strategies for the gene panel seed selection. To ensure the panel contains sufficient information to differentiate cell types in the biological system of interest, we need to build a seed panel that includes sufficient genes capable of distinguishing cell types while keeping the panel size as small as possible. To obtain such a gene panel seed, first, we will run multiple different panel design methods, leveraging established algorithms such as: HVGs (Highly Variable Genes), DEGs (Differentially Expressed Genes), RandomForest, Spapros^18^, ScGeneFit^19^ on the training dataset. Two alternative strategies are implemented to define the seed gene panel: (1) **Pareto optimal Gene Panel**: the Agent will study the stability of the Adjusted Rand Index or ARI between leiden clustering results and ground truth under different panels size across all methods, then it will select the resultant subpanel from the method that showcased overall best ARI and stability combination to form the seed. (2) **RL enhancement**: While the default Pareto optimal gene panel design approach is often effective, this alternative may offer better performance although generally with a much higher computational footprint. When deciding the seed gene panel, if the user specified to use the RL enhancement option, PantheonOS will launch the RL enhancement option. With RL enhancement, PantheonOS will gather the subpanels by finding an optimal subset of genes coming from the union set of the subpanels from the individual methods. (3) **Biological prior lookup and context matching**: Once the seed is determined, PantheonOS will lookup for genes not already in the seed but reported by literature, website and database like Genecards, Uniprot, Oncogene, related to the user query. This will ensure the system to include genes on top of the seed panel that will match biological prior constraints imposed by the user to complete the gene panel. (4) **Consensus filling**: If after seed determination and inclusion of genes based on biological priors, there is still some room in the gene panel, PantheonOS will perform consensus analysis to fill the gene list. Specifically, after running the predefined gene-selection methods, each method produces a ranked list of genes according to its own selection criterion. PantheonOS then aggregates these rankings, followed by applying a rank normalization across methods, operating on relative rank positions rather than raw scores. For instance, if a gene is ranked 3rd by one method and 10th by another, its normalized consensus rank can be computed by averaging these ranks. The final gene panel is constructed iteratively by following the resulting consensus ranking and adding the highest-priority genes that are not yet included, until the user-specified panel size is reached. By doing so, we can include additional genes to fill the gap based on the consensus rank across all methods. In the end, each accepted gene is assigned to a biological category, contextually justified with respect to the target objective and supported by appropriate references. (5) **Benchmarking**: Once PantheonOS has established the final gene panel, it benchmarks the gene panel on validation batches (Default 5 batches not seen during training). It computes the default adjusted rand index or ARI, normalized mutual information or NMI, silhouette index or SI, or Procrustes similarity or PS for the final gene set from PantheonOS (Final) and panels from different algorithms. To allow flexibility, Pantheon also allows computing other metrics that can be specified by the user input. In the end, PantheonOS will return a nicely formatted pdf report summarizing the full workflow and all the results in figures, tables. Thus, PantheonOS provides a general framework for gene panel design for targeted spatial transcriptomics that effectively reconciles maximal cell type information preservation and explicit biological knowledge constraints from the user queries.

### RL algorithm for seed panel generation

The objective of the reinforcement-learning (RL) module is to construct a targeted-size gene panel for spatial transcriptomics that maximally preserves cell-type separability, while allowing flexible combinations of genes beyond those produced by individual classical selection methods. The approach builds upon the formalism introduced by ^70^. This RL-based refinement is an enhancement to the deterministic seed-panel construction and will be activated when explicitly requested by the user.

Overall, gene panel design is formulated as a sequential decision-making problem and solved using an actor–critic reinforcement learning framework. The agent iteratively refines a gene panel by proposing add/remove actions, guided by a reward that balances clustering accuracy and panel compactness. A permutation-invariant encoding enables fixed-dimensional state representations, while knowledge injection from existing methods initializes learning near biologically meaningful configurations. The framework supports both decentralized multi-agent and centralized GPU-optimized implementations. Specifically, gene panel selection is to maximize biological separability while keeping the panel size close to a target *K*_*target*_ . It is framed as a sequential decision-making problem to iteratively refine the panel. The search space is restricted to a candidate set, 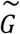 formed by the union of genes selected by classical methods (HVG, DEGs, scGeneFit, Spapros, Random Forest) to ensure computational tractability. Two equivalent RL implementations are supported: (1) **CPU mode (Multi-Agent RL):** Each candidate gene is treated as an agent that votes for its inclusion/exclusion, following a decentralized multi-agent Markov Decision Process (MDP). (2) **GPU mode (Centralized RL):** A single centralized agent directly learns an optimal gene-combination policy, optimized for GPU-friendly batched operations. For both implementation, it involves **state representation** and **actions and panel transitions**, which we will describe in details next:

#### State Representation

The current gene panel (*Gt*) is encoded into a fixed-dimensional, permutation-invariant representation. This representation aggregates global panel statistics (e.g., mean, variance, sparsity, normalized size of the expression matrix), pooled over all genes in the current gene panel, and is mapped into a latent state *z*_*t*_ for the policy and value networks.

#### Actions and Panel Transitions

The agent proposes modifications (adding or removing genes) to the current panel. Panel transitions are constrained to remain close to the requested size, preventing uncontrolled growth or collapse of the gene set. To enable panel transition, the RL module follows an **actor–critic architecture**: (1) The **Critic network** evaluates the fitness of a panel that is quantified by a function that we call value Function *V*ψ(*z*_*t*_) which reflects how good a panel is; (2) The **Actor network** proposes panel modifications, improving the policy via gradient ascent guided by the Critic’s estimates. Next, we design the **reward function** to balance biological alignment and panel compactness, defined as *r* (*G*_*t*_) = *α ARI* (*G*_*t*_) + (1 − α)size_term (*G*_*t*_), where **ARI (Adjusted Rand Index)** measures the agreement between Leiden clustering using the panel and ground-truth cell-type labels while the **Size Term** is a smooth penalty that stabilizes the panel size towards the desired size.

To train the RL framework we use the following training **pipeline** that involves **Expert-informed initialisation, exploration** and **updates**. (1) **Expert-informed initialisation**: Panels obtained from established gene selection methods (HVG, DE, scGeneFit, SpaPROS) are used to initialize the replay buffer, providing the agent with biologically grounded starting configurations.. (2) **Exploration**: epsilon-greedy exploration is used, and the exploration rate is gradually decreased to favor exploitation as training progresses. That is to say at every step we will choose with a probability (1-epsilon) to use the action decided by the actor, otherwise we choose a random action. Therefore we ensure that in the beginning we explore many states while the actor is training, and once it gets sufficiently trained we can use its actions. (3) **Updates**: Actor and Critic networks are updated via temporal-difference learning and policy-gradient objectives. (Below is an overview of the loss optimisation of actor and critic networks. The critic network minimization is conducted via the following loss function

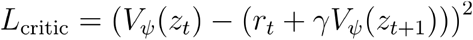

Where *V ψ* (*z*_*t*_): Critic neural network that evaluates the value function at state *z*_*t*_,*r*_*t*_ : reward at the same state. Similarly, the actor minimization is conducted via the following loss function

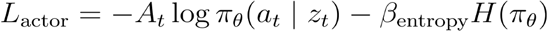

Where π_*θ*_ : Actor network that chooses the action, *a*_*t*_: Action taken at state t, *H*(*p*) = − (*p* log + (1 − *p*) log (l − p) is just an entropy term over the distribution of selected genes to encourage exploration. Furthermore, The advantage function is computed as follows (and typically represents how “advantageous” it is to go from gene panel *G*_*t*_ to gene panel *G*_*t*+1_.

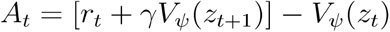

## Pantheon-Evolve framework: Agentic Code Evolution

### Core Algorithm

Pantheon-Evolve introduces a method for evolutionary optimization based on the genetic algorithm^71^ guided by agents. Unlike prior approaches such as AlphaEvolve^24^ or FunSearch^72^, which typically use LLMs as stateless, single-pass mutation operators (*f*: *C* → *C*′), this framework implements a two-phase mutation architecture. This design separates reasoning from implementation: an analyzer agent first performs diagnostic analysis, using auxiliary tools and a Python interpreter for experimentation, before a mutator agent executes targeted code transformations.This separation enables the system to conduct a more autonomous and strategic exploration of the algorithmic design space ℂ .

This workflow is formally described in Algorithm 5, which uses an iterative loop to refine the program population.

### System Architecture

The system is managed by a controller that runs an asynchronous evolution loop across *N* parallel workers (`num_workers`). The core data structure is a program database (*D*) that implements the Multi-dimensional Archive of Phenotypic Elites (MAP-Elites) algorithm^27^. This archive maintains a population *P* distributed across *K* distinct “islands” (ℐ = {*I*_1_, …, *I*_*K*_ }), simultaneously optimizing for solution quality (fitness *J*) and behavioral diversity (feature coverage ϕ). Each agent within the system (analyzer, mutator, feedback) is initialized with configurable LLM backends (`analyzer_model, mutator_model`), subject to specific temperature parameters (*T*) and token constraints.

### Evolutionary Algorithm (MAP-Elites with Island Topology)

The genotype is represented as a codebase snapshot *C*, which captures the state of a multi-file codebase. Evolution occurs via discrete search-and-replace block mutations. The phenotype is mapped via a feature extraction function ϕ: ℂ → ℤ^*d*^ to a discrete feature grid ℳ (default dimensions: complexity × speed). Each cell in the grid retains only the highest-performing program:

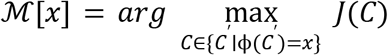

To prevent premature convergence, the system uses an island-based architecture with periodic migration (default: ring topology, each island only exchanges individuals with its immediate neighbors) occurring every *T*_*mig*_ generations. Parent selection follows a mixed strategy distribution defined in `Select` (`strategy=“Mixed”`) :

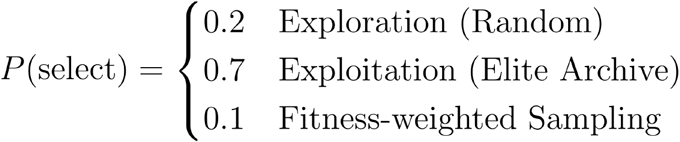

### Agent-Based Code Evolution

The core evolutionary step is defined by a three-phase loop:

1. **Phase 1 - Analyzer Agent (***A*_*gent*_): Performs strategic analysis using a reasoning-plus-action approach. It receives the parent code *C*_*parent*_, the optimization objective *0*, evolution history *H*_*history*_ (ancestor chain and sibling attempts) and several inspiration programs *P*_*insipiration*_ randomly sampled from MAP-Elites on the island they belong to.
  - **Adaptive Exploration Strategy:** The analyzer toggles between exploration (algorithm-level changes) and exploitation (code-level optimization) based on a time-dependent probability *P*(*g*) that corresponds to the exploration at generation *g*:

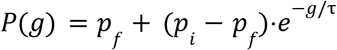

Where *g* is the current generation, *p*_*i*_ = 0. 9 is the initial probability, *p*_*f*_ = 0. 1 is the final probability, and τ ≈ *G*/2. 3 ensures decay within the generation budget *G* .
  - **Output:** A structured optimization plan (π) detailing the identified problem, proposed solution, and specific location for modification.
2. **Phase 2 - Mutator Agent (**ℳ_*ut*_ ): Executes precise code changes based on the plan π, without redundant context analysis.

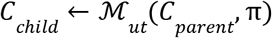
3. **Phase 3 - Summarizer Agent:** Extracts a structured mutation summary from the change diff, updating the evolution history *H*_*history*_ for future iterations.

### Hybrid Evaluation System

The system employs a hybrid fitness function *J*(*C*) that combines quantitative metrics from the sandbox execution (*S*_*func*_) with qualitative feedback from an LLM code reviewer (*S*_*llm*_):

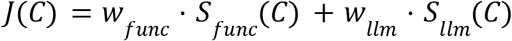

Default weights are set to *w*_*func*_ = 0. 8 and *w*_*llm*_ = 0. 2. Runtime errors and debugging output (*E*_*err*_) are captured and fed back into the analyzer in subsequent iterations for self-correction.

### Diversity and Quality Maintenance

The population is managed with a configurable limit (default: 500 programs), maintaining an elite archive representing the top 25% of solutions. Diversity is enforced through the MAP-Elites grid mechanism, where programs compete only within their specific feature bin. To accommodate shifting phenotypic distributions, the system uses adaptive feature ranges, dynamically adjusting grid boundaries with padding (default: 10%) as evolution progresses.

### Application and Usage

Pantheon-Evolve is a general purpose evolutionary framework for many complex scientific and engineering problems with a quantifiable fitness function. In this work, we use it for evolving the state-of-the-art single cell batch correction algorithm, Harmony and our RL algorithm for gene panel design. It is worth noting that even if we cannot define a performance-based fitness function, we can also use the computational speed reduction and accuracy preservation as the fitness function to improve the computational algorithm improvement.

#### Algorithms 5

The Pantheon-Evolve Framework

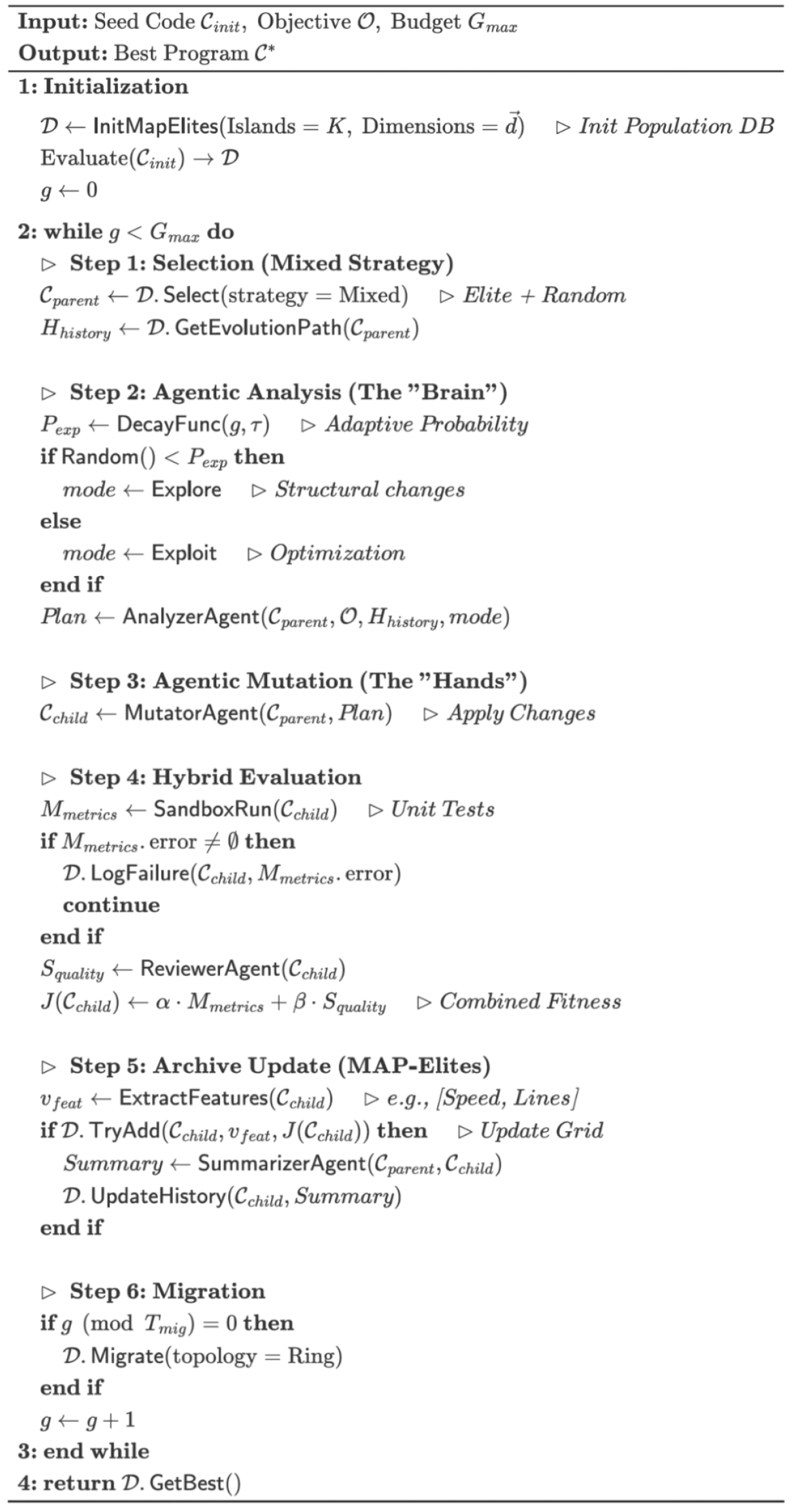

### Pantheon-Evolve Integration and Visualization

The PantheonOS agent ecosystem seamlessly integrates Pantheon-Evolve with other agents and comprehensive reporters for evolution analysis and debugging. PantheonOS provides a series of designs to allow other agents to seamlessly interact with Pantheon-Evolve:

(1) Evolve Agent in Omics-Expert-Team interfaces with Pantheon-Evolve module for algorithm optimization requests; (2) EvolutionToolSet exposes *evolve_code*/*evolve_codebase* tools to agents, with session management and progress tracking via evolution_id; (3) Evolved algorithms stored in code libraries managed by Coding Agent, enabling reuse across analysis workflows; (4) Evolution state saved as JSON with checkpoint support for resumability. In addition, we develop the Evolution Reporter for comprehensive visualization and analyses of the evolution: (1) Interactive HTML report generation via EvolutionVisualizer class with D3.js-based evolution tree showing parent-child mutation relationships; (2) Score history line plot tracking fitness progression across iterations with best-score envelope; (3) MAP-Elites heatmap displaying quality-diversity grid coverage across feature dimensions; (4) Diff viewer for examining code changes at each mutation step with syntax highlighting; (5) Program detail panels showing metrics, LLM feedback, and evaluation artifacts. Thus, the integration layer enables Pantheon agents to leverage evolutionary optimization as a tool, while the reporter system provides interpretable insights into the evolution process, supporting both automated workflows and human-in-the-loop analysis of algorithm improvements.

## Evolving state-of-the-art batch correction algorithms with Pantheon-Evolve

Justification why we choose batch correction algorithm (can move to the main text):

1. As the single cell genomics continue to democratize in the research community, increasingly more labs will generate scRNA-seq data that inevitably lead to profile the same system with different technologies and protocols, resulting in significant batch effect
2. Even within the same lab, batch effects may occur because different people or machines generate the data.
3. The emergency of large-scale atlasing initiatives such as human cell atlas also make the batch correction as one of the first prominent tasks for analyzing such data.
4. The metrics for the performance of batch correction are well established and there are a large-range algorithms for us to explore the capability of Pantheon-evolve

To demonstrate the evolutionary capability of PantheonOS, we apply Pantheon-Evolve to evolve state-of-the-art batch correction algorithms for single-cell RNA-seq data with domain-specific configurations, including Harmony for single-file evolution and Scanorama or BBKNN for multi-file codebase evolution.

The universal Pantheon-Evolve framework has several shared configurations for **feature dimensions, evaluation metrics, function weight** and **checkpoint interval**, reserved for all evolutionary applications. When applying to batch correction algorithms, we tailored our configurations for the specific needs. Specifically, we define *batch mixing score*, biological conservation score, *speed score* as the feature dimensions for batch corrections. The Evaluation Metrics, implemented through a script (*metrics*.*py)*, include all the following scores across all batch correction evolution cases:

1. **Batch Mixing Score**: We compute the global batch proportions and compare them to each cell’s local neighborhood. The global proportion of batch *b* (where *n* is the total number of cells and *b*_*i*_ is the batch label of cell *i*) is:

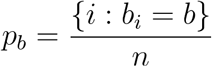

For each cell *i*, we find its k-nearest neighbors (*k* is the neighborhood size) and compute the local proportion of batch *b* in that neighborhood:

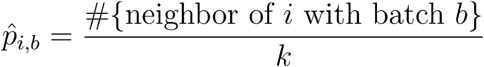

Using the difference between local and global proportions across all *B* batches (*B* is the number of batches), the per-cell mixing score is *s*_*i*_ defined as:

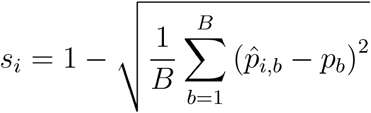

Finally, the batch mixing score is the average of *s*_*i*_ over all cells:

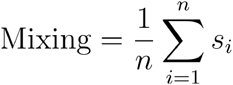
2. **Biological Conservation Score**: Silhouette score computed on the batch-corrected embedding using true cell type labels. For each sample *i*, we compute the average distance to all other points in its own cluster *C* :

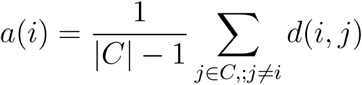

And the minimum average distance to points in any other cluster:

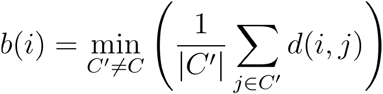

The silhouette value is then:

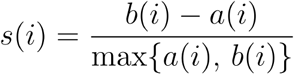

The overall silhouette score is the mean of *s* (*i*) across all samples:

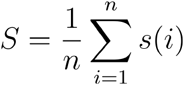

It measures how well cells of the same type cluster together after correction.
3. **Speed Score**: We define the speed score as an inverse function of the runtime. Let *t*_exec_ denote the execution time in seconds. To keep the score bounded in (0,1] and smoothly decreasing with longer runtimes, we use a reciprocal form:

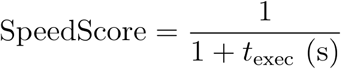

This gives a score close to 1 when *t*_exec_ is very small, and it decays toward 0 as *t*_exec_ grows. The + 1 term prevents division by zero and ensures the score is always well-defined. Score approaches 1.0 for near-instant execution and decreases asymptotically toward 0 for longer runtimes.

Furthermore, for each specific algorithm to be evolved, we further tailor evolution parameters based on its complexity (single vs multi-file), computational requirements (timeout), and optimization priorities (fitness weights reflecting algorithm-specific trade-offs). In addition, in all evolution examples, the LLM we use is GPT-5.2. We list the detailed configurations for all evolution parameters in the following:

### Harmony

#### Objective prompt

~~~
***Optimize the Harmony algorithm implementation for:
1. **Integration Quality** (40% weight): Improve batch mixing while preserving biological structure.
 - The algorithm should effectively remove batch effects
 - Biological clusters should remain distinct after correction
2. **Performance** (20% weight): Reduce execution time.
 - Optimize hot loops and matrix operations
 - Consider vectorization opportunities
 - Avoid redundant computations
3. **Convergence** (10% weight): Improve convergence behavior.
 - Reduce number of iterations needed
 - Ensure stable convergence
4. **Biological Conservation** (30% weight): Preserve biological variance.
 - Don’t over-correct and remove biological signal
 - Maintain cluster separation
Constraints:
- Keep the public API (run_harmony function signature)
- Maintain numerical stability
- Don’t remove essential functionality***
~~~

#### Other configurations

- Target File: harmony.py
- Evolution Config:
  - num_workers: 8, num_islands: 3
  - Num_inspirations (Number of inspiration programs in the analyzer agent context): 2, num_top_programs: 3
  - evaluation_timeout: 120s, analyzer_timeout: 120s
  - early_stop_generations: 200
- Fitness Weights:
  - Integration Quality: 40% (batch mixing)
  - Biological Conservation: 30% (preserve variance)
  - Performance: 20% (execution time)
  - Convergence: 10% (iteration count)

### Scanorama

#### Objective prompt

~~~
***Optimize the Scanorama batch correction algorithm for:
1. **Integration Quality** (45% weight): Improve batch mixing while preserving biological structure.
 - The algorithm uses mutual nearest neighbors (MNN) to find correspondences
 - The ‘assemble()’ function applies nonlinear corrections using RBF kernels
 - The ‘transform()’ function computes bias vectors for correction
2. **Biological Conservation** (45% weight): Preserve biological variance.
 - Don’t over-correct and remove biological signal
 - Maintain cell type separation after correction
3. **Performance** (10% weight): Reduce execution time.
 - The ‘nn_approx()’ function uses Annoy for approximate nearest neighbors
 - The ‘batch_bias()’ function can be a bottleneck for large datasets
 - Consider vectorization opportunities in ‘mnn()’ and ‘transform()’
## File Structure (3 files):
### scanorama/scanorama.py (Main Algorithm)
Key functions to consider optimizing:
- ‘integrate()’ / ‘correct()’: Main entry points
- ‘assemble()’: Core panorama assembly (orchestrates alignment and correction)
- ‘find_alignments()’: Finds matching cells between datasets
- ‘transform()’: Computes nonlinear bias vectors using RBF kernel
- ‘mnn()’: Mutual nearest neighbor detection
- ‘nn_approx()’: Approximate nearest neighbor search
- ‘batch_bias()’: Computes smoothed bias vectors (potential bottleneck)
### scanorama/utils.py (Utilities)
- ‘reduce_dimensionality()’: PCA-based dimensionality reduction
- ‘handle_zeros_in_scale()’: Numerical stability helper
### scanorama/ init .py (Package Init)
- Just exports from scanorama.py
## Key Parameters (in scanorama.py):
- ALPHA (0.10): Alignment score minimum cutoff
- KNN (20): Number of nearest neighbors
- SIGMA (15): RBF kernel smoothing parameter
- DIMRED (100): Dimensionality for integration
## Constraints:
- Keep the public API (integrate, correct function signatures)
- Maintain numerical stability (handle edge cases)
- Don’t break imports between files
- Keep the package structure intact
## Areas for Algorithm-Level Improvement:
- The RBF kernel in transform() could use different kernel functions
- The alignment scoring in find_alignments() could be improved
- The panorama merging strategy in assemble() could be optimized
- Consider adaptive sigma based on local density***
~~~

#### Other configurations

- Evolution Mode: Multi-file via CodebaseSnapshot.from_directory()
- Target Files: scanorama/ (3 files)
  - scanorama.py: Main algorithm (integrate, assemble, transform, mnn)
  - utils.py: Dimensionality reduction helpers
  - init .py: Package exports
- Evolution Config:
  - num_workers: 8, num_islands: 2
  - num_inspirations: 1, num_top_programs: 2
  - evaluation_timeout: 180s, max_code_length: 80000
  - early_stop_generations: 100
- Fitness Weights:
  - Integration Quality: 45% (MNN-based correction)
  - Biological Conservation: 45% (cell type separation)
  - Performance: 10% (RBF kernel optimization)
- Key Parameters: ALPHA=0.10, KNN=20, SIGMA=15, DIMRED=100

### BBKNN

#### Objective prompt

~~~
***Optimize the BBKNN (Batch Balanced KNN) algorithm for:
1. **Integration Quality** (45% weight): Improve batch mixing while preserving biological structure.
- BBKNN identifies each cell’s top neighbors in each batch separately
- The ‘get_graph()’ function constructs the batch-balanced KNN graph
- The ‘compute_connectivities_umap()’ function computes the fuzzy simplicial set
2. **Biological Conservation** (45% weight): Preserve biological variance.
- Don’t over-correct and remove biological signal
- Maintain cell type separation in the resulting graph
- The ‘trimming()’ function affects connectivity strength
3. **Performance** (10% weight): Reduce execution time.
- The ‘create_tree()’ function builds KNN indices (uses cKDTree by default)
- The ‘query_tree()’ function performs neighbor queries
- Consider algorithmic improvements to ‘get_graph()’
## File Structure (2 files):
### bbknn/ init .py (High-level API)
- ‘bbknn()’: Main AnnData-based entry point (wraps matrix.bbknn)
- ‘ridge_regression()’: Optional preprocessing
- ‘extract_cell_connectivity()’: Helper for visualization
### bbknn/matrix.py (Core Algorithm)
Key functions to optimize:
- ‘bbknn()’: Scanpy-independent entry point (takes pca and batch_list)
- ‘get_graph()’: Constructs the batch-balanced KNN graph
- ‘create_tree()’: Creates KNN index for each batch
- ‘query_tree()’: Queries neighbors from each batch
- ‘compute_connectivities_umap()’: Computes fuzzy simplicial set
- ‘trimming()’: Trims connectivities to top values
## Key Parameters:
- neighbors_within_batch (3): Top neighbors per batch
- n_pcs (50): PCA dimensions to use
- trim: Connectivity trimming threshold
- metric: Distance metric (euclidean, manhattan, etc.)
- set_op_mix_ratio: UMAP fuzzy set mixing parameter
- local_connectivity: UMAP local connectivity parameter
## Constraints:
- Keep the public API (bbknn, matrix.bbknn function signatures)
- Maintain compatibility with sparse matrix outputs
- Don’t break imports between files
- Ensure numerical stability
## Areas for Algorithm-Level Improvement:
- The neighbor selection strategy in get_graph() could be improved
- The connectivity computation could use different kernels
- Adaptive neighbors_within_batch based on batch sizes
- Different distance metrics or weighted combinations
- Improved trimming strategies***
~~~

Other configurations:

- Evolution Mode: Multi-file via CodebaseSnapshot.from_directory()
- Target Files: bbknn/ (2 files)
  - init .py: High-level AnnData API
  - matrix.py: Core algorithm (get_graph, create_tree, query_tree, trimming)
- Evolution Config:
  - num_workers: 8, num_islands: 8
  - num_inspirations: 1, num_top_programs: 2
  - evaluation_timeout: 180s, max_code_length: 80000
  - early_stop_generations: 100
- Fitness Weights:
  - Integration Quality: 45% (batch-balanced KNN graph)
  - Biological Conservation: 45% (connectivity preservation)
  - Performance: 10% (tree query optimization)
- Key Parameters: neighbors_within_batch=3, n_pcs=50, metric=euclidean

## Evolving RL algorithm for gene panel selection with Pantheon-Evolve

We demonstrate Pantheon-Evolve can be used to evolve a RL-based gene panel selection algorithm for single-cell RNA-seq data, newly designed by us. This demonstration involves evolution of a complex reinforcement learning system with batched actor-critic networks, exploration strategies, and reward shaping. Furthermore, it includes two experimental configurations comparing different evolution priorities (exploration vs. optimization). In the following, we first itemize the share configurations for both evolutionary cases, followed by introducing further configuration details for each case.

### Shared Configuration

#### Feature Dimensions

- **size_score**: Normalized panel size (panel_size / K_max), distinguishes an extremely exhaustive (i.e. 1000 genes) panel from a just right panel (K_target, defaulting to 500) to a too sparse panel (i.e. 0 genes)
- **prior_overlap**: Fraction of selected genes that overlap with prior method results, measures exploitation vs. exploration
- **convergence_improvement**: Early improvement / overall improvement ratio, measures fast vs. slow-and-steady convergence
- **reward_spread**: Coefficient of variation or CV of reward history, measures exploratory vs. exploitative behavior. Define mean of the reward in the history trajectory is and variance is, the CV is calculated as 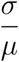

### Evaluation Metrics (evaluator.py, shared across all cases)

- **Adjusted Rand Index (ARI)**: Clustering agreement with ground truth cell types. Computed by: (1) principal component analysis or PCA on selected genes, (2) Leiden clustering in the PCA space, (3) ARI between clusters and true labels. The range is [0, 1], higher is better. Calculated using sklearn’s adjusted_rand_score function, called on the true cell type labels versus the clusters derived from the gene panel genes

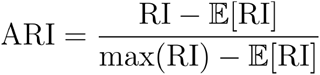

Where

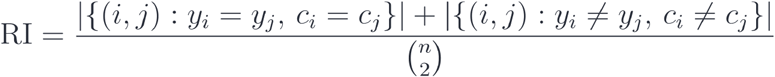

for true cell type labels Y and predicted clusters C.
- **Normalized Mutual Information (NMI)**: Information-theoretic clustering quality metric. Measures shared information between predicted clusters and true cell types. The range is [0, 1], higher is better. Calculated using sklearn’s normalized_mutual_info_score, called on the true cell type labels versus the clusters derived from the gene panel genes

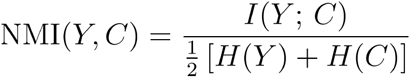

where H(*) is the Shannon entropy, Y are the true cell type labels, and C are the predicted cluster labels and I(*) is the mutual information between Y and C.
- **Separation Index (SI)**: Inter-cluster distance / intra-cluster distance ratio in PCA space. Measures how well-separated clusters are. Higher values indicate better cluster separation. Calculated across all of the predicted clusters:

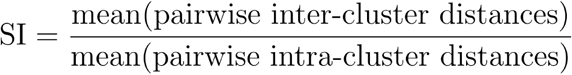
- **Training Speed**: Inverse time metric 1/(1 + training_time/60). Encourages faster convergence without over-prioritizing speed. Approaches 1.0 for fast training.

### Evolution Settings

- function_weight: 1.0, llm_weight: 0.0 (pure metric-based fitness)
- checkpoint_interval: 5 iterations
- early_stop_generations: 50 (original), 100 (explore primary)
- Epochs of training per evolution: 25

## Case 1: Original/Baseline Configuration (results_original_primary)

**Target File**: rl_gene_panel.py (single file evolution)

### Evolution Config

- num_workers: 1 (limited by GPU memory for RL training)
- num_islands: 2
- num_inspirations: 2
- num_top_programs: 3
- max_parallel_evaluations: 2 (limited by GPU memory)
- evaluation_timeout: 600s (10 minutes for RL training)
- analyzer_timeout: 180s
- early_stop_generations: 100

### Fitness Weights

- Final ARI: 35% (primary clustering quality)
- Final NMI: 35% (complementary clustering metric)
- Final SI: 20% (cluster separation quality)
- Training Speed: 10% (convergence efficiency

### Evolution Prompt

~~~
Optimize the RL-based gene panel selection algorithm for:
1. **Final Panel Quality** (90% weight): Maximize ARI, NMI, SI clustering
metrics.
 - ARI (Adjusted Rand Index) measures clustering agreement with ground truth
   - NMI (Normalized Mutual Information) provides additional clustering quality
signal
 - SI (Separation Index) measures inter-cluster vs intra-cluster distances
2. **Training Efficiency** (10% weight): Faster convergence to good solutions.
 - Reduce number of epochs needed to find good panels
 - Improve early convergence behavior
 - Better reward signal leads to faster learning
Evolution Targets (priority order):
1. **Exploration strategy (SmartCurationTrainer.explore)** - Primary target
 - Current: epsilon-greedy with Gaussian noise, top-K selection
  - Possible improvements: Boltzmann/temperature-based sampling, UCB-style
exploration,
elite gene preservation, adaptive noise schedules
2. **Optimization (SmartCurationTrainer.optimize)** - Secondary target
 - Current: single-step TD, fixed entropy coefficient
  - Possible improvements: GAE (Generalized Advantage Estimation), entropy
scheduling,
advantage normalization, PPO-style clipping
3. **Reward function (reward_panel)** - Tertiary target
 - Current: ‘reward = alpha * ari + (1 - alpha) * size_term’
  - Possible improvements: multi-metric rewards (include NMI/SI), progressive
penalties,
 non-linear ARI scaling, diversity bonuses for pathway coverage
Constraints:
- **Keep public API unchanged**: ‘train_gene_panel_selector’ function signature
must remain stable
- **Maintain numerical stability**: Clamp probabilities, handle edge cases
properly
- **Don’t remove essential functionality**: Keep core RL components working
- **Keep imports stable**: Don’t add new external dependencies
~~~

### Key Parameters

- K_target: 500 genes (target panel size, no penalty below this)
- K_max: 1000 genes (maximum allowed, zero reward above)
- alpha: 0.8 (weight for ARI vs size penalty in reward)
- beta: 1.5 (shape parameter for size penalty curve)
- N_explore: 8 (exploration steps per epoch)
- N_optimize: 5 (optimization steps per epoch)

## Case 2: Optimize-Primary Configuration (results_explore_primary)

**Target File**: rl_gene_panel.py (same file, different evolution priorities)

### Evolution Config

- num_workers: 1
- num_islands: 2
- num_inspirations: 2
- num_top_programs: 3
- max_parallel_evaluations: 2
- evaluation_timeout: 600s
- analyzer_timeout: 180s
- early_stop_generations: 50 (reduced from 100 for faster experimentation)

### Fitness Weights (Same as Case 1)

- Final ARI: 35%
- Final NMI: 35%
- Final SI: 20%
- Training Speed: 10%

### Evolution Prompt

~~~
Optimize the RL-based gene panel selection algorithm for:
1. **Final Panel Quality** (90% weight): Maximize ARI, NMI, SI clustering
metrics.
 - ARI (Adjusted Rand Index) measures clustering agreement with ground truth
  - NMI (Normalized Mutual Information) provides additional clustering quality
signal
 - SI (Separation Index) measures inter-cluster vs intra-cluster distances
2. **Training Efficiency** (10% weight): Faster convergence to good solutions.
 - Reduce number of epochs needed to find good panels
 - Improve early convergence behavior
 - Better reward signal leads to faster learning
Evolution Targets (priority order):
1. **Optimization (SmartCurationTrainer.optimize)** - Primary target
 - Current: single-step TD, fixed entropy coefficient
  - Possible improvements: GAE (Generalized Advantage Estimation), entropy
scheduling,
 advantage normalization, PPO-style clipping
2. **Exploration strategy (SmartCurationTrainer.explore)** - Secondary target
- Current: epsilon-greedy with Gaussian noise, top-K selection
 - Possible improvements: Boltzmann/temperature-based sampling, UCB-style
exploration,
 elite gene preservation, adaptive noise schedules
3. **Reward function (reward_panel)** - Tertiary target
 - Current: ‘reward = alpha * ari + (1 - alpha) * size_term’
  - Possible improvements: multi-metric rewards (include NMI/SI), progressive
penalties,
 non-linear ARI scaling, diversity bonuses for pathway coverage
Constraints:
- **Keep public API unchanged**: ‘train_gene_panel_selector’ function signature
must remain stable
- **Maintain numerical stability**: Clamp probabilities, handle edge cases
properly
- **Don’t remove essential functionality**: Keep core RL components working
- **Keep imports stable**: Don’t add new external dependencies
~~~

### Key Parameters (Same as Case 1)

- K_target: 500, K_max: 1000
- alpha: 0.8, beta: 1.5
- N_explore: 8, N_optimize: 5

## Single cell foundation model router

PantheonOS integrates 22 mainstream single-cell foundation models through an intelligent routing mechanism that automatically selects the optimal model based on data characteristics, task requirements, and hardware availability. In the following, we will explain the key building blocks of the router, including routing architecture, model registry, intelligent model selection, adaptive routing, and execution environment.

### Routing architecture

The routing architecture is general and flexible to allow the autonomic selection of optimal foundation models. Firstly, we design a unified interface that abstracts model-specific APIs through a common toolset with centralized model registry. Secondly, the router agent is implemented to allow it to accept analysis requests from users or other agents in the multi-agent team. Thirdly, isolated execution environments prevent dependency conflicts between heterogeneous model implementations. Lastly, we use standardized output format to ensure consistent downstream integration regardless of source model.

### Model Registry

The registry registers 22 single cell foundation models covering diverse model architectures, training data scales (33M to 126M cells), and specializations. These 22 models cover modeling of many different modalities, including RNA, ATAC, spatial transcriptomics, protein, and multi-omics. In addition, they support analyses of up to 7 organisms. Furthermore, they support a large range of downstream analyses, including producing latent dimension embeddings of single cells, batch integration, cell type annotation, perturbation prediction, and spatial aware analyses.

### Intelligent Model Selection

The model selection follows a transparent pipeline, including: (1) Data profiling: Automatic detection of species, modality, and gene identifier format from input data; (2) Hardware awareness: Considers GPU availability and memory constraints for model suitability; (3) Constraint filtering: Narrows down foundation model candidates based on task type, species compatibility, and resource requirements; (4) LLM-based ranking: Matches user intent and data characteristics against model-specific differentiators; (5) Interpretable output: Provides selection rationale and ranked alternatives for transparency.

### Adaptive Routing

We implement three complimentary strategies to enable adaptive routing:

(1) Priority-based selection: Unique capability requirements take precedence over general task matching; (2) Fallback hierarchy: Graceful degradation through ranked alternatives when primary selection is unavailable; (3) Automatic rerouting: Detects data-model incompatibility and switches to compatible alternatives

### Execution Environment

Firstly, we isolate the computational environment to ensure reproducibility across diverse model dependencies. Next, we implement a hardware-adaptive execution mechanism to route the computation to CPU or GPU based on availability and model requirements. Additionally, we leverage a unified output schema to standardize embeddings, predictions, and provenance metadata across all models. Lastly, we develop a general automatic preprocessing approach to handle gene ID conversion and normalization for model compatibility.

In conclusion, the routing mechanism abstracts the complexity of single cell foundation model selection, enabling users to leverage the growing ecosystem of foundation models through a unified interface with interpretable selection decisions.

## Supplementary methods

### Analysis details of the human fetal heart single-cell multiomics and 3D MERFISH+ data

To demonstrate Pantheon’s capability for integrative multi-modal spatial analysis, we interactively chat with the Omics-Expert-Team, leveraging built-in skills for optimal transport mapping with Moscot, 3D visualization with PyVista, and cell-cell communication analysis with Spateo to comprehensively jointly analyze a single cell multi-omics data of human fetal heart and 3D spatial profiling of an entire human fetal heart to unveil the spatially informed molecular mechanisms underlying congenital heart disease.

We firstly use PantheonOS to integrate single cell multi-omics data with 3D MERFISH+ data via Optimal Transport. PantheonOS calls built-in Single-Cell to Spatial Mapping skill to invoke Moscot’s unbalanced optimal transport algorithm to integrate single-cell multiomics data (95,684 cells, multi-timepoint) of human fetal heart samples with 3D MERFISH+ spatial data (3M cells, 238 genes) of human fetal heart from post-conception week 12. To address memory constraints, PantheonOS utilizes a batch mapping strategy (20,000-cell batches). After the integration, PantheonOS enables bidirectional transfer for cell type label prediction and gene expression imputation.

We next perform spatially aware disease gene expression analysis with PantheonOS. Congenital heart disease-associated genes (241 genes in total curated from CHDgene database^38,40^) aggregated into specific heart disease-associated expression scores per cell through agent-generated analysis code. With the computed expression scores, we can then use PantheonOS to visualize their spatial distribution in the 3D space leveraging built-in 3D Spatial Visualization skill using PyVista to render disease patterns with interactive rotation and GIF animation.

In addition, to understand the disease vulnerability difference between males and females for different heart diseases, we also performed Sex Difference Analysis. Firstly, PantheonOS stratifies the single-cell data by sex which are then mapped to the MERFISH+ 3D data with separate optimal transport mappings. Next, PantheonOS computes differential expression patterns by comparing sex-specific imputed values at each spatial location. Thirdly, spatial neighbor composition differences are quantified between sexes.

Next, to study the sex differences of different spatial neighborhood and communication, we firstly compute pairwise cell type proximity statistics for neighbor enrichment via agent-generated KNN-based code. Next we calculate the enrichment scores as normalized co-occurrence frequencies. PantheonOS then uses 3D analysis skills to run ligand-receptor interaction analysis via the Spateo package to identify significant communication pairs.

Lastly, PantheonOS is used to reveal the heart diseases-associated enhancers’ chromatin accessibility in 3D space. It firstly computes enhancer activation scores per cell per disease gene by aggregating chromatin accessibility reads across enhancer regions, using the scE2G^42^ links from the original paper to connect enhancers to their target genes. It then visualizes the disease-aggregated ATAC patterns in 3D space mapped from the single cells via the optimal transport established in the above.

Thus, the workflow demonstrates seamless integration of built-in skills (Moscot mapping, PyVista visualization, Spateo communication analysis) with agent-generated custom analysis code, orchestrated through natural language dialogue to enable comprehensive multi-modal spatial analysis that may shed light on the underlying molecular and cellular mechanism of congenital heart diseases without manual programming.

### OpenST experiment for early mouse embryo

Adult C57BL/6J mice (The Jackson Laboratory) were housed under standard conditions with free access to food and water. For timed mating, sexually mature females were paired with males of the same strain, and the presence of a vaginal plug was assessed the following morning. Plug-positive females were designated as pregnant, with the day of plug detection defined as 0.5 days post-coitum (E0.5).

Embryos at E6.0 were collected from plug-positive pregnant females. Following euthanasia, uterine horns were isolated and transferred to ice-cold DMEM supplemented with 10% FBS. Decidual tissues were opened under a stereomicroscope, and embryos were dissected from the decidua. Reichert’s membrane was mechanically removed using fine forceps and a tungsten needle. Embryo developmental stages were determined based on established morphological criteria as described previously^73^. All procedures related to animals were performed in accordance with the ethical guidelines of the University of Texas Southwestern Medical Center (UTSW). Animal protocols were reviewed and approved by the UTSW Institutional Animal Care and Use Committee (IACUC) before any experiments were performed (Protocols #2018-102430).

Six freshly isolated E6.0 embryos were maintained on ice whenever possible to minimize sample deterioration. To prevent osmotic damage caused by direct exposure to OCT compound (Tissue-Tek, Cat# 4583), embryos were equilibrated stepwise in sucrose/PBS solutions (10%, 20%, and 30%) on ice, with each incubation performed for 5 min. When embryos exhibited nonspecific adhesion to plastic surfaces, BSA/PBS (0.5 g/mL stock) was diluted 1:100 into the equilibration solution to reduce attachment.

After cryoprotection, a small drop of the equilibration solution containing the embryos was placed on the inner surface of a 35-mm plastic dish lid. For visualization, 0.4% trypan blue solution (Gibco, Cat# 15250061) was added at approximately one-third to one-half of the drop volume, avoiding vigorous mixing. Trypan blue staining enhanced specimen contrast and visibility, thereby facilitating reliable identification of embryos during subsequent cryosectioning. The embryo-containing drop was gently placed onto the OCT top surface in the embedding molds (Simport, Cat# M475-1). Excess equilibration solution surrounding the embryos was carefully removed using a fine glass capillary to minimize residual aqueous solution. Embryos were then guided from the OCT top surface toward the mold bottom using a tungsten needle. Positional adjustments were performed by generating gentle OCT flow rather than direct mechanical pressure. This guiding procedure promoted gradual replacement of the surrounding solution with OCT compound. Embryos were positioned centrally at the mold bottom whenever feasible. Consistent anterior–posterior orientation was not enforced due to the limited visibility of morphological landmarks within OCT compound. Molds were rapidly frozen in a liquid nitrogen–based freezing system and stored at −80 °C until further processing.

After embedding, the OCT block containing the tissue array was mounted onto a cryostat for sectioning. Tissue sections of 10 µm thickness were collected onto OpenST slides pre-patterned with spatially indexed DNA barcodes. The sections were then fixed in pre-cooled methanol (−20 °C) for 30 minutes and stained for histological imaging. Following tissue permeabilization, released mRNAs hybridized to the surface-immobilized barcodes and were reverse-transcribed in situ at 42 °C overnight to generate spatially barcoded cDNA. The tissue was subsequently removed, and the resulting cDNA was amplified by PCR and purified using 1× AMPure XP beads. The libraries were sequenced using a paired-end 150-bp (PE150) strategy. Sequencing reads were then mapped back to their spatial coordinates based on the known barcode layout with PantheonOS’s OpenST data upstream processing skill, enabling reconstruction of gene expression patterns across the tissue section.

### Data analysis details for OpenST data

OpenST data per slide

| slide_id | n_cells | n_genes |
| --- | --- | --- |
| A9_L1 | 7867 | 29950 |
| B8_L3 | 7838 | 30204 |
| D8_L1 | 13182 | 30844 |
| D9_L3 | 6789 | 28495 |
| E4_L2 | 3623 | 23988 |
| E7_L2 | 1589 | 21412 |

#### 3D Spatial Transcriptomics Reconstruction of Mouse Embryo E6.0

To construct the three-dimensional (3D) spatial transcriptome of mouse embryos at embryonic day 6.0 (E6.0), we developed a multi-step computational pipeline that integrates OpenST sequencing data with corresponding hematoxylin and eosin (H&E)-stained histological images. The pipeline consists of four major stages: (1) fiducial marker calibration in OpenST data, (2) alignment of OpenST data to H&E images via fiducial markers, (3) serial section registration of H&E images into a unified 3D coordinate system, and (4) gene expression-guided refinement using Spateo. All spatial transformations were composed and propagated to map OpenST bead coordinates into a common 3D reference frame.

#### Fiducial Marker Detection and Calibration in OpenST Data

The OpenST capture areas contain two alternating types of fiducial marker patterns, each consisting of a 2×4 grid of circular markers with two concentric ring sizes. To precisely determine the fiducial marker positions, we first aggregated bead-level spatial coordinates and total UMI counts into 2D binned images (bin size = 20 units) for each puck column corresponding to each fiducial marker type. The binned images were then binarized after Gaussian smoothing (kernel size 3×3, σ = 2) and intensity thresholding. Circular fiducial markers were detected using the Hough Circle Transform (OpenCV ‘HoughCircles’, dp = 1.2, minDist = 400, param2 = 10, radius range 54–63 pixels). Detected circles were filtered by their x-coordinates to retain only markers at the expected grid positions. To improve robustness, we detected circles across all four OpenST sections (A9_L1, B8_L3, D8_L1, D9_L3), and averaged their positions. These calibrated fiducial marker templates were used for subsequent alignment to H&E images.

#### Alignment of OpenST Data to H&E-Stained Images

For each tissue section, the OpenST spatial transcriptomics data was aligned to the corresponding H&E-stained image using fiducial markers as landmarks. We first generated a binned count image (bin size = 50 units) from the bead spatial coordinates and total UMI counts. The pre-calibrated fiducial marker templates were overlaid on the binned image according to each puck’s tile type and spatial offset. In parallel, fiducial markers in the H&E images were detected using the Hough Circle Transform with parameters adapted for the image resolution. Corresponding fiducial marker pairs between the OpenST binned image and the H&E image were manually identified. Using these paired landmarks, we estimated a similarity transformation via a Procrustes analysis with least-squares optimization. The similarity transformation *T*_bin → HE_ maps coordinates from the binned OpenST space to the H&E image space. The transformation was validated by computing residuals between transformed source points and target points, and by visual inspection of the warped overlay.

### Serial Section Registration of H&E Images

To reconstruct a 3D volume, adjacent H&E-stained sections (slices 6 through 22, representing 17 serial sections) were registered pairwise. Prior to matching, embryo tissue regions in each H&E image were segmented using the Segment Anything Model (SAM)^74^ to generate binary masks, and background regions were removed by setting non-tissue pixels to black.

For each pair of adjacent sections, we employed RoMa (Robust Dense Feature Matching)^75^, a state-of-the-art image matching model pre-trained on natural images, to establish dense correspondences. RoMa was configured with a coarse resolution of 560×560 and an upsampling resolution of 864×1152. The model outputs a dense warp field and pixel-wise certainty scores. To suppress spurious matches in background regions, the certainty map was multiplied by a spatial mask derived from the tissue segmentation labels of both images. From the masked certainty-weighted warp field, 3,000 keypoint correspondences were sampled. A rigid transformation was then estimated from the sampled correspondences using RANSAC^76^ (10,000 iterations, inlier threshold = 5.0 pixels). For two section pairs (slices 14→15 and 16→17) where direct adjacent matching failed due to large structural differences or tissue loss, we performed skip-one matching (i.e., slices 14→16 and 16→18).

The pairwise rigid transformations were organized into a graph, where each node represents a section and each edge stores the estimated 3×3 homogeneous transformation matrix. Both forward and inverse (via matrix inversion) transformations were stored. Starting from a reference section (slice 6), global transformations *T*_*i* → ref_ for all sections were computed via breadth-first search (BFS) traversal, composing pairwise transforms along the shortest path:

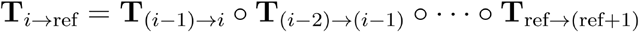

All H&E images were then warped into the reference coordinate system using their respective global transforms.

### Composition of Spatial Transformations and 3D Coordinate Assignment

For each OpenST section, the final transformation from raw bead coordinates to the global H&E reference frame was composed as:

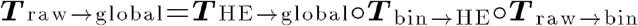

where *T*_raw → bin_ accounts for the coordinate shift and scaling between raw bead coordinates and the binned image space, *T*_bin → HE_ is the fiducial-based similarity transform, and *T*_HE → global_ is the global rigid transform from serial section registration. Each bead’s 2D coordinates were transformed to the global reference frame, producing the aligned coordinates. A uniform scaling factor was applied to convert pixel coordinates to physical units. The z-coordinate for each section was assigned based on the section index.

### Spatial Binning and Embryo Segmentation

To facilitate downstream analysis, bead-level data was aggregated into spatial bins (bin size = 200 units). For each bin, gene expression counts were summed and spatial coordinates were averaged. The six individual embryos present in the tissue sections were manually segmented in 3D.

### Gene Expression-based Alignment Refinement with Spateo

To further refine the 3D reconstruction, we applied Spateo’s alignment method (spateo.align.morpho_align) to one representative embryo (Embryo 3). This method leverages gene expression similarity, in addition to spatial proximity, to optimize pairwise section alignment. For each pair of adjacent sections, Spateo performed alignment using the following parameters: maximum 200 iterations, partial robust level of 10, batch size of 2,000, PCA embeddings as the expression representation, and cosine dissimilarity as the distance metric. These pairwise rigid corrections were sequentially composed to propagate alignments across all sections relative to the first section:

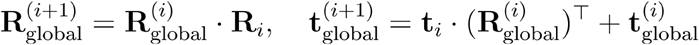

The refined 2D coordinates were combined with the original z-coordinates to produce the final 3D spatial transcriptome.

### Spatial Domain and Signaling Gradient Analysis by PantheonOS

To achieve a unified spatial transcriptomic landscape of the E6.0 mouse embryo, we integrated six independent biological replicates using an agent-optimized Harmony workflow within the PantheonOS. Raw count matrices from openST sections were first preprocessed using omicverse (v.1.7.9) ^33^. PantheonOS directly performed normalized, log-transformed and Highly variable genes (HVGs) using the `omicverse.pp.preprocess` function. To mitigate batch effects while preserving biological developmental gradients, PantheonOS utilized Pantheon-Evolve Harmony, which incorporates a log-domain Sinkhorn transport plan to optimize cell-type alignment across replicates. The integrated latent space was then used for UMAP visualization and downstream clustering.

To resolve cell-type compositions within the OpenST capture bins at bin200 resolution, PantheonOS performed spatial deconvolution using Tangram^35^ (v.1.0.3), orchestrated by the Pantheon-Agent. PantheonOS utilized the TOME^34^ (Trajectories Of Mouse Embryos) atlas as the single-cell RNA-seq reference. The reference data were filtered to include only genes shared with the spatial dataset. The `omicverse.space.Deconvolution` function was executed with the “cell” mode to learn the mapping matrix. This process projected single-cell transcriptomic profiles onto the spatial coordinates, allowing for the imputation of cell-type proportions and non-measured gene expression. The fidelity of the mapping was validated by calculating the correlation between observed and imputed expression of canonical markers (*Pou5f1, S100a6*).

To define the ExE–Epiblast signaling boundary, an automated spatial analysis pipeline generated and performed by PantheonOS. A binary spatial mask was first generated by thresholding the deconvolved *Cer1* intensity. The boundary was computationally derived using a contour detection algorithm on the *Cer1* spatial domain. For each spatial bin, we calculated the signed distance to the nearest point on the *Cer1* boundary, where negative values represent the proximal (ExE-facing) side and positive values represent the distal (Epiblast-facing) side. To analyze the paracrine inhibitory effect, we performed a distance-dependent regression of *Nodal* mRNA expression relative to the *Cer1* boundary. The geometric axis N was calculated as the local normal to the ExE-Epiblast boundary curve. For each cell, a signed distance was assigned based on its projection onto N, effectively mapping the 3D embryonic structure onto a 1D polar-like coordinate to resolve the Cer1-Nodal inhibitory interface.Statistical significance between the identified spatial domains was assessed using a two-sided Mann-Whitney U test.

### Spatial Signal Smoothing via Weighted Kernel Density Estimation

To resolve continuous spatial patterns of gene expression within the Pantheon-OmicVerse Skill, spatial smoothing was performed by computing weighted Gaussian kernel density estimates (KDE) on 2D embeddings to transform sparse, stochastic signals into spatially-correlated probability density fields. For each target feature, a validity mask excluded non-finite coordinates, and a minimum expression threshold was applied to filter out background noise, utilizing only transcriptionally active cells as “training points” for the KDE model. These raw expression values were min-max scaled to a [0, 1] range to ensure weight comparability across features before a Gaussian kernel was fitted to the coordinates of the training cells using the scaled expression as statistical weights. The kernel bandwidth was controlled by a scaling factor to balance local detail with global smoothness, and the resulting density function was evaluated across the coordinates of all cells in the dataset to map discrete expression counts into a continuous distribution, facilitating the robust identification of signaling interfaces like the ExE-Epiblast boundary.

## Supplementary Figures

### High Resolution Figures in PNG format can be found at Figures

**Figure S1.**
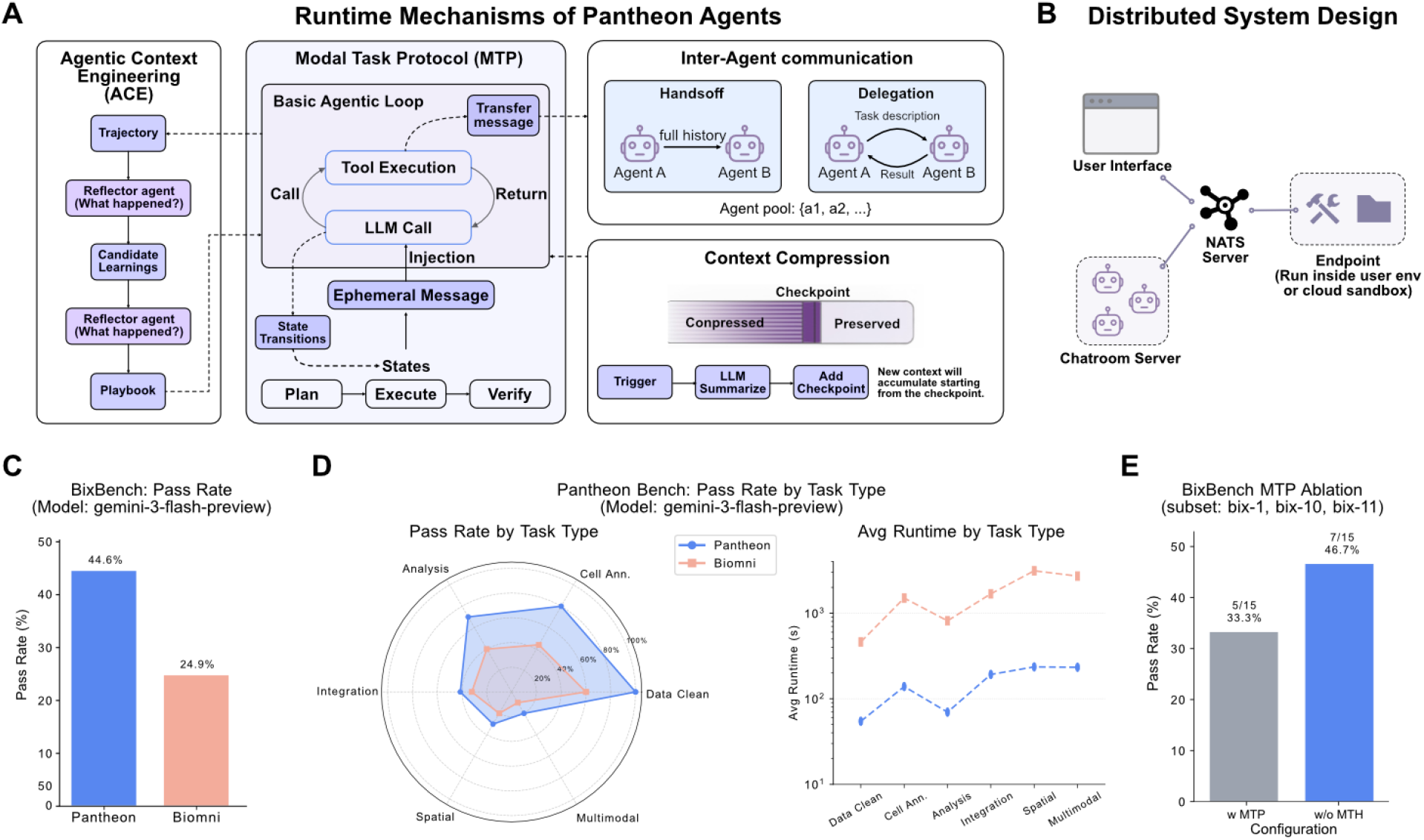
Runtime mechanisms and benchmarks for pantheon agents. **(A)** The runtime mechanism of an Agent consists of five parts: Basic Agentic Loop, Modal Task Protocol (MTP), Inter-Agent Communication, Context Compression, and Agentic Context Engineering (ACE69). Except for the Basic Agentic Loop, the other parts are optional integrations. **(B)** PantheonOS’s distributed system design connects the UI, Chatroom server, and Endpoint through a NATS server. The Agents and tools can run separately and interconnect over a network. This mechanism provides the PantheonOS architecture with significant flexibility and allows users to complete analyses without uploading data to the platform, thereby ensuring data privacy. **(C)** Comparison of results between Pantheon and Biomni6 on BixBench78, using the gemini-3-flash-preview model for testing. **(D)** Pantheon-Bench (a benchmark containing various types of single-cell data analysis tasks) compares Pantheon and Biomni, with all tests uniformly conducted using the gemini-3-flash-preview model. The left panel shows the pass rate for different types of tasks, while the right panel displays a running time comparison. **(E)** MTP ablation experiments demonstrate the effectiveness of the MTP mechanism in enhancing the agent’s analytical capabilities.

**Figure S2.**
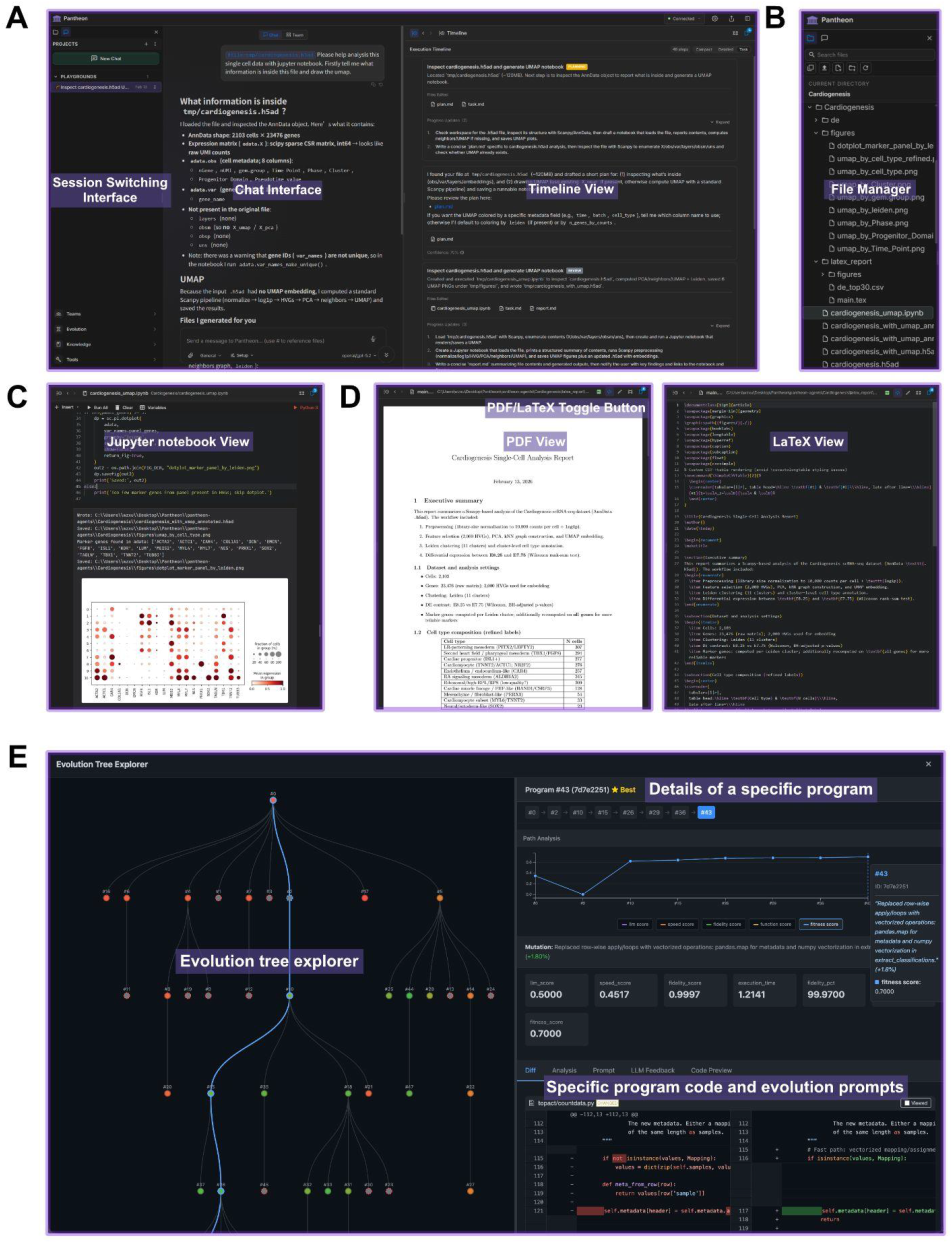
Pantheon web-based graphical user interface (GUI) for interactive genomics analysis and evolution visualization. **(A)** Overview of the main GUI layout. **Left**: session switching panel for managing multiple analysis sessions. **Center**: chat interface enabling natural language interaction with the agent teams such as the Omics-Expert-Team, showing an example single-cell data analysis conversation with agent-generated code, results, and visualizations. **Right**: execution timeline view displaying the chronological progression of agent activities, task dispatching, and intermediate outputs for real-time workflow monitoring. **(B)** Integrated file manager, accessible by switching from the left sidebar of the main interface shown in panel **A**, providing hierarchical navigation of analysis outputs, including figures, intermediate data files, and generated reports organized by analysis session. **(C)** Embedded Jupyter notebook view, accessible by switching from the right panel of the main interface (panel **A**), enabling interactive inspection, modification and execution of agent-generated analysis code and real-time visualization of results such as heatmaps and scatter plots. **(D)** Automated report generation module, also accessible from the right panel of the main interface (A), with a PDF/LaTeX toggle button. **Left**: rendered PDF view of a publication-ready analysis report including executive summary, methods, and tabular results. **Right**: corresponding LaTeX source view for direct editing and customization before export. **(E)** Pantheon-Evolve visualization dashboard. **Left**: interactive evolution tree explorer displaying parent–child mutation relationships across iterations, where node color indicates fitness level. **Top right**: detailed view of a selected program, including path analysis tracing its evolutionary lineage, mutation description, and quantitative evaluation metrics. **Right bottom**: side-by-side code diff viewer showing the specific code modifications and evolution prompts applied at each mutation step, enabling transparent inspection of agent-guided algorithmic changes.

**Figure S3.**
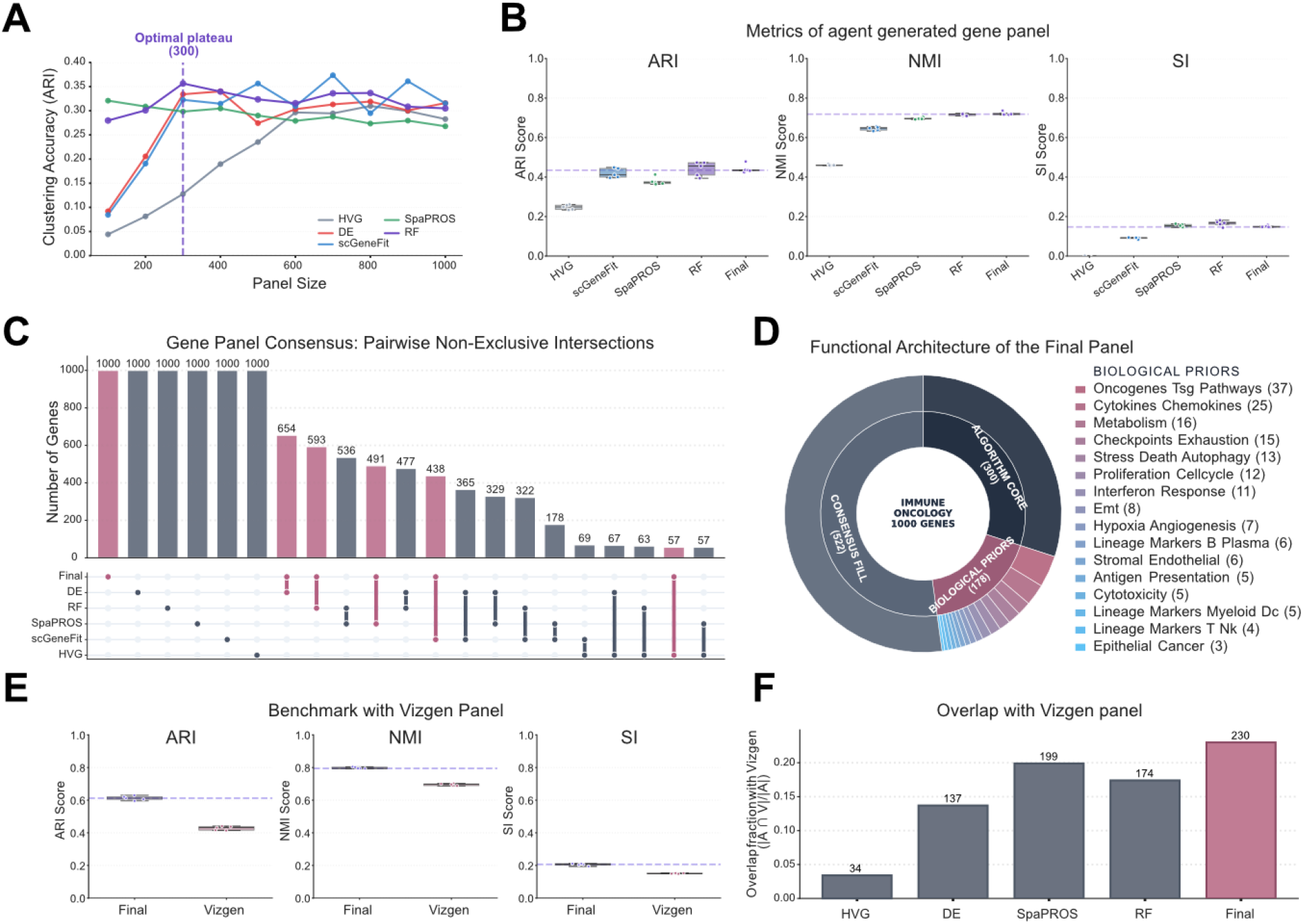
Fully automatic 1000-gene gene panel design for immuno-oncology applications. **(A)** The accuracy curves plotted against the number of genes for various methods are shown. The agent selects a Pareto optimum based on these curves to determine the size of the seed panel, which is 300 in this case. Pareto optimum means a solution that can not be further improved without letting other objectives get worse, in this case without increasing the panel size the ARI score cannot be further improved. **(B)** Performance metrics (ARI, Normalized Mutual Information or NMI, and Silhouette Index or SI) of the agent-generated final gene panel (final) compared to traditional methods (HVG, scGeneFit19, SpaPROS18, RF). **(C)** Overlap between the final gene panel and panels derived from alternative selection methods (DE, RF, SpaPROS18, scGeneFit19, HVG), shown as a UpSet79 plot where pairwise non-exclusive intersections with bar heights indicate the number of shared genes. **(D)** Functional architecture of the final 1,000-gene immune-oncology panel, showing the relative contributions of the algorithm core (300 genes), consensus fill (522 genes), and biological priors (178 genes); the biological priors segment is further subdivided into pathway and cell-state categories (legend). **(E)** Benchmark of the final gene panel against the Vizgen panel (expert-curated, commercial), comparing clustering performance across replicates with the original cell type annotations using ARI, NMI, and silhouette index (SI) scores (boxplots show score distributions). **(F)** Overlap of gene panels with the Vizgen panel (expert-curated, commercial), reporting the fraction (and number) of shared genes for HVG, DE, SpaPROS, RF, and the final panel.

**Figure S4.**
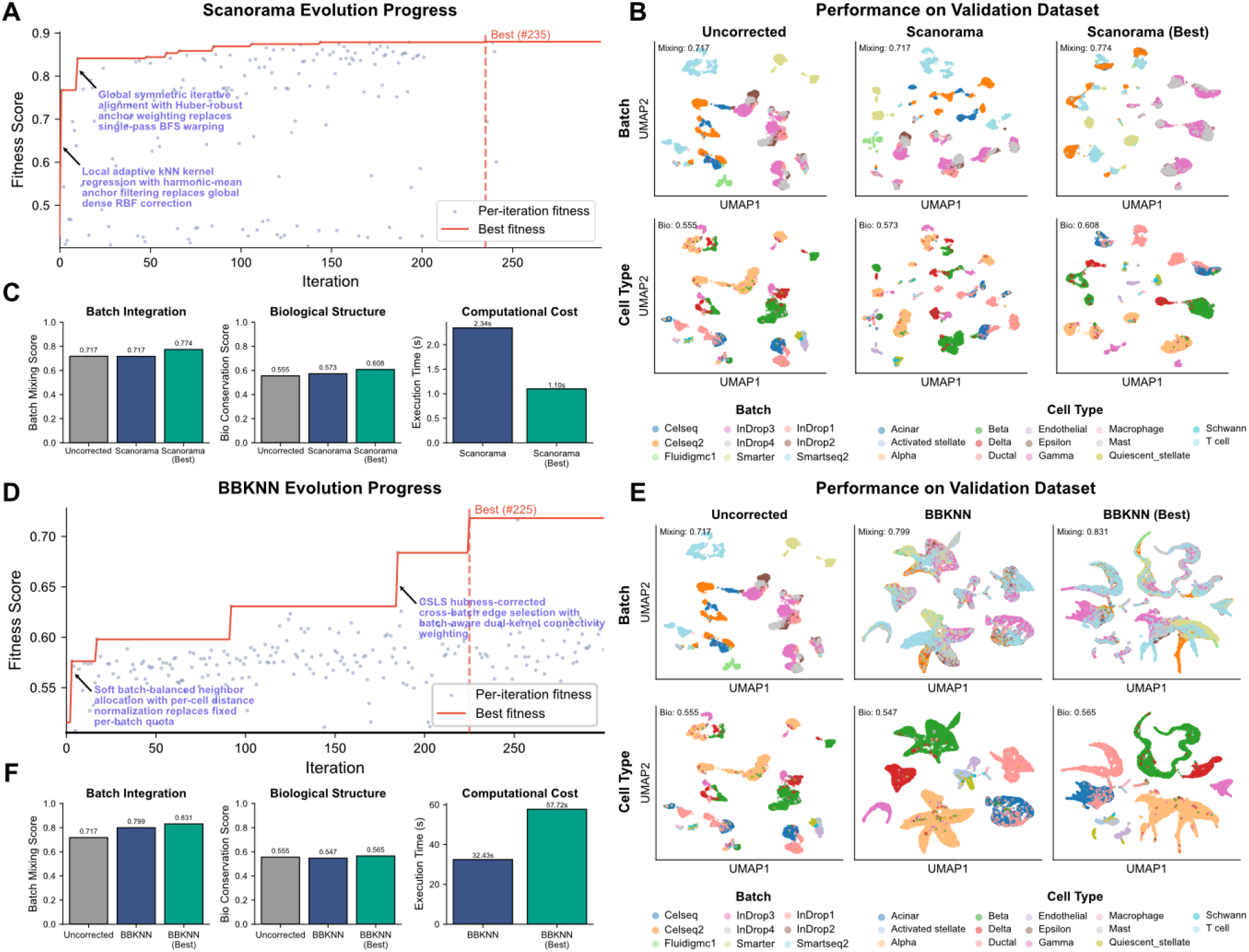
Evolutionary performance of Pantheon-Evolve on Scanorama8 and bbknn9. **(A)** Evolution trajectory of the Scanorama algorithm in 300 iterations. Grey dots represent per-iteration fitness scores; the red line traces the best fitness achieved. Key algorithmic innovations discovered during evolution are annotated, including replacement of global dense RBF correction with local adaptive kNN kernel regression and harmonic-mean anchor filtering (early stage), and replacement of single-pass BFS warping with global symmetric iterative alignment and Huber-robust anchor weighting (later stage). The best-performing program (#235) is indicated by the dashed line. **(B)** Performance on the validation dataset (9 batches, 14 cell types) comparing uncorrected data, original Scanorama, and the best evolved variant (Scanorama Best, #235). Top row: UMAP embeddings colored by batch identity with batch mixing scores. Bottom row: UMAP embeddings colored by cell type with biological conservation scores. The evolved variant achieves improved batch mixing while better preserving biological structure. **(C)** Quantitative comparison of batch integration score, biological conservation score, and computational cost between uncorrected, original Scanorama, and the evolved variant. The evolved Scanorama improves both integration quality and biological conservation while also reducing execution time (1.10s vs. 2.34s). **(D)** Evolution trajectory of the BBKNN algorithm over ∼275 iterations. Notable mutations include the introduction of soft batch-balanced neighbor allocation with per-cell distance normalization (early stage) and online sequential least squares or OSLS hubness-corrected cross-batch edge selection with batch-aware dual-kernel connectivity weighting (later stage), with the best program (#225) marked by the dashed line. **(E)** Performance on the same validation dataset comparing uncorrected data, original BBKNN, and the best evolved variant (BBKNN Best, #225). The evolved variant achieves substantially improved batch mixing while maintaining comparable biological conservation. **(F)** Quantitative comparison for BBKNN evolution. The evolved variant achieves improved batch integration and biological conservation scores at the cost of increased computational time.

**Figure S5.**
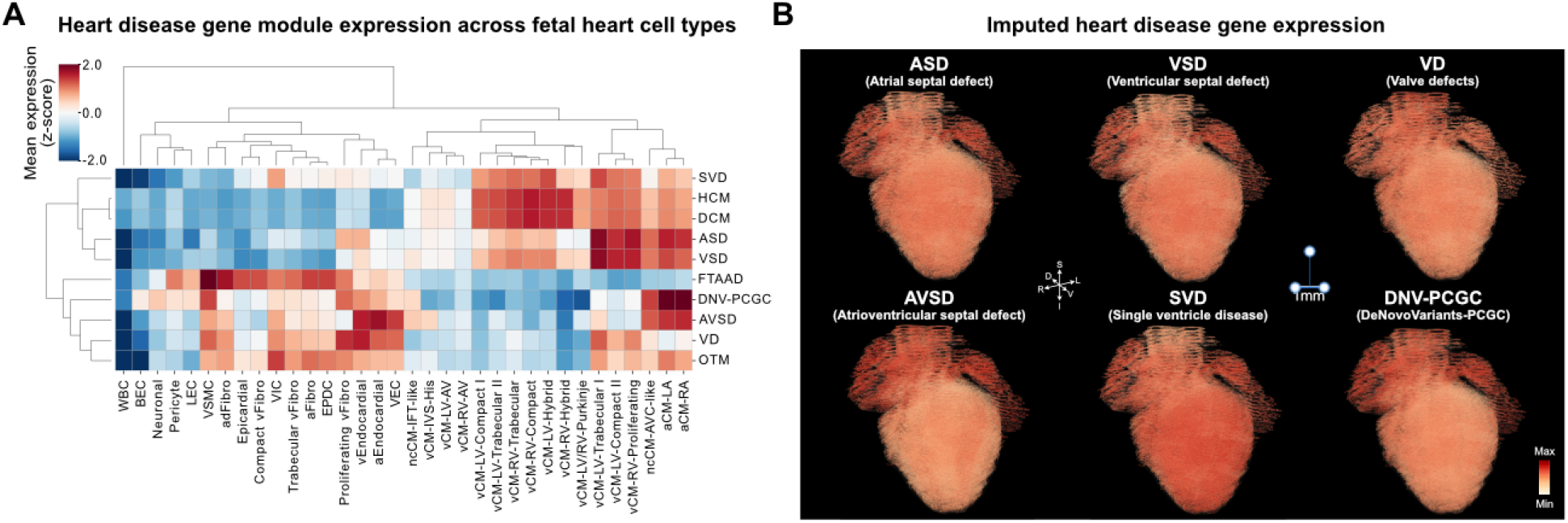
Cell type-resolved heart disease gene module expression in the 3D human fetal heart. **(A)** Hierarchically clustered heatmap showing mean expression (column-wise z-score) of 10 heart diseases gene modules across 34 MERFISH-defined cell types. Gene module scores were computed as the mean imputed expression of all genes associated with each disease phenotype. Red indicates higher relative expression; blue indicates lower. **(B)** 3D spatial visualization of imputed RNA expression for six heart diseases gene modules not shown in the main figure: atrial septal defect (ASD), ventricular septal defect (VSD), valve defects (VD), atrioventricular septal defect (AVSD), single ventricle disease (SVD), and PCGC de novo variants (DNV-PCGC). Color intensity (red) represents mean expression level across genes in each module. Scale bar, 1 mm.

**Figure S6.**
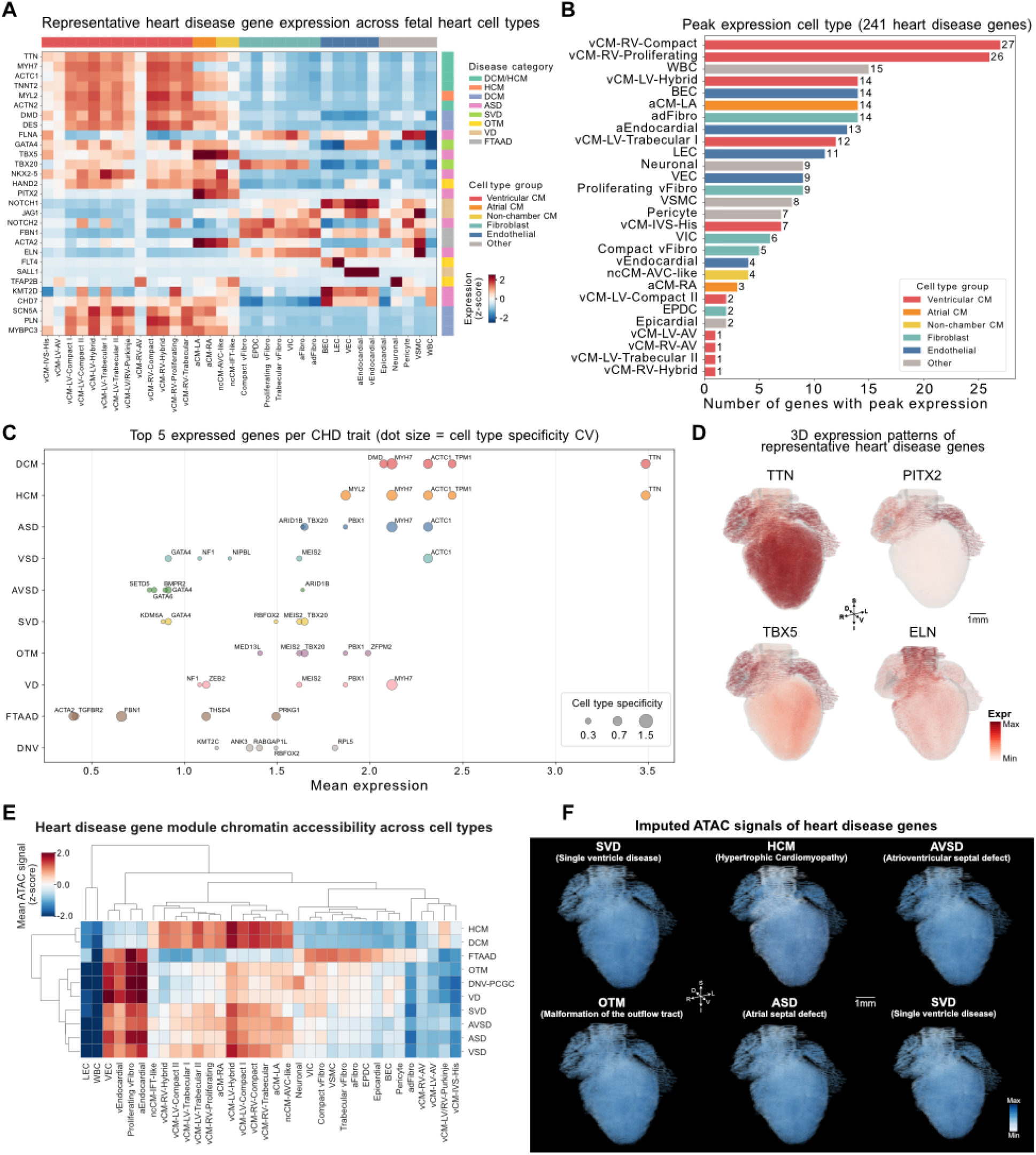
Cell type-resolved expression and chromatin accessibility landscape of heart disease genes in the 3D human fetal heart. **(A)** Heatmap of 29 representative heart diseases gene expression levels across 34 MERFISH-defined cell types. Values are row-wise z-scores of mean imputed expression. Genes (rows) are annotated by their primary associated disease category (left color bar); cell types (columns) are grouped by lineage (top color bar): ventricular cardiomyocytes (Ventricular CM), atrial cardiomyocytes (Atrial CM), non-chamber cardiomyocytes (Non-chamber CM), fibroblasts, endothelial cells, and others. Sarcomeric genes (TTN, MYH7, ACTC1, TNNT2, MYL2, ACTN2) show strong enrichment in ventricular cardiomyocytes, while transcription factors (TBX5, PITX2) are restricted to atrial cardiomyocytes, and vascular/ECM genes (ELN, FBN1) are enriched in stromal populations. **(B)** Number of heart disease genes (out of 241 total) with peak expression in each cell type. Right ventricular compact cardiomyocytes (vCM-RV-Compact, 27 genes) and proliferating cardiomyocytes (vCM-RV-Proliferating, 26 genes) harbor the most peak-expressed heart disease genes, followed by white blood cells (WBC, 15 genes). Bars are colored by cell type lineage group. **(C)** Top 5 most highly expressed genes per heart disease trait. Dot position along the x-axis represents mean expression across all cells; dot size encodes cell type specificity (coefficient of variation across cell types). Larger dots indicate genes with more cell type-restricted expression. Sarcomeric genes (TTN, ACTC1, MYH7, TPM1) dominate DCM and HCM with high expression and moderate-to-high cell type specificity, while developmental transcription factors and signaling molecules characterize structural defects (ASD, VSD, AVSD, SVD, OTM, VD). **(D)** 3D spatial expression patterns of four representative heart disease genes illustrating distinct spatial domains: TTN (diffuse ventricular expression), PITX2 (left atrial-restricted), TBX5 (atrial-enriched), and ELN (vascular/stromal-restricted). Color intensity represents imputed expression level. Scale bar, 1 mm. **(E)** Hierarchically clustered heatmap of heart disease gene module chromatin accessibility (ATAC signal, column-wise z-score) across cell types, analogous to (A). ATAC signals were imputed onto the 3D MERFISH atlas via MOSCOT mapping of single-cell multiome ATAC data. **(F)** 3D spatial visualization of imputed ATAC signals for six heart diseases gene modules: single ventricle disease (SVD), hypertrophic cardiomyopathy (HCM), atrioventricular septal defect (AVSD), malformation of the outflow tract (OTM), atrial septal defect (ASD), and single ventricle disease (SVD). Color intensity (blue) represents mean chromatin accessibility. Scale bar, 1 mm.

**Figure S7.**
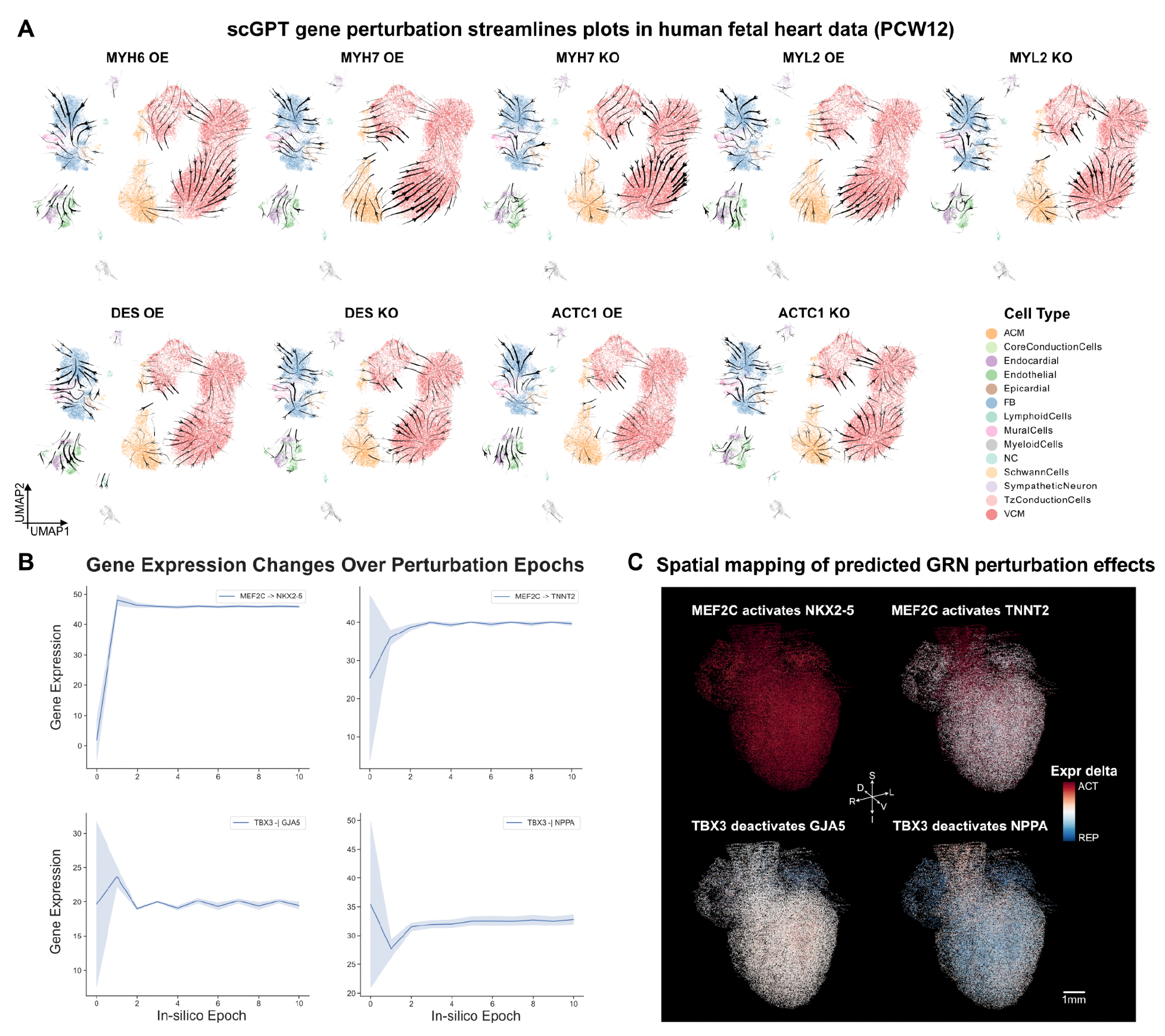
In-silico gene perturbation and gene regulatory network inference in PCW12 human fetal heart. **(A)** scGPT-based in-silico gene perturbation streamline plots on UMAP embeddings of PCW12 fetal heart scRNA-seq data. Overexpression (OE) and knockout (KO) perturbations are shown for cardiac structural genes MYH6, MYH7, MYL2, DES, and ACTC1. Streamlines indicate predicted cell state transition trajectories upon perturbation. Cells are colored by annotated cell type (14 types). **(B)** Tabula-predicted target gene expression changes over in-silico perturbation epochs for four literature-validated regulatory pairs. Top: MEF2C activates NKX2-5 and TNNT2, showing progressive upregulation of target expression. Bottom: TBX3 represses GJA5 and NPPA, showing downregulation of target expression. Lines and shaded areas represent mean ± s.d. across perturbed cells. **(C)** Spatial mapping of predicted GRN perturbation effects onto 3D MERFISH fetal heart coordinates via unbalanced optimal transport (moscot). Expression changes (epoch 9 minus epoch 0) from panel B are imputed onto ∼100k spatially resolved cells. Red indicates activation (increased expression) and blue indicates repression (decreased expression). Anatomical axes are indicated: D, dorsal; V, ventral; R, right; L, left; S, superior. Scale bar, 1 mm.

## Supplementary Tables

**Table S1.**
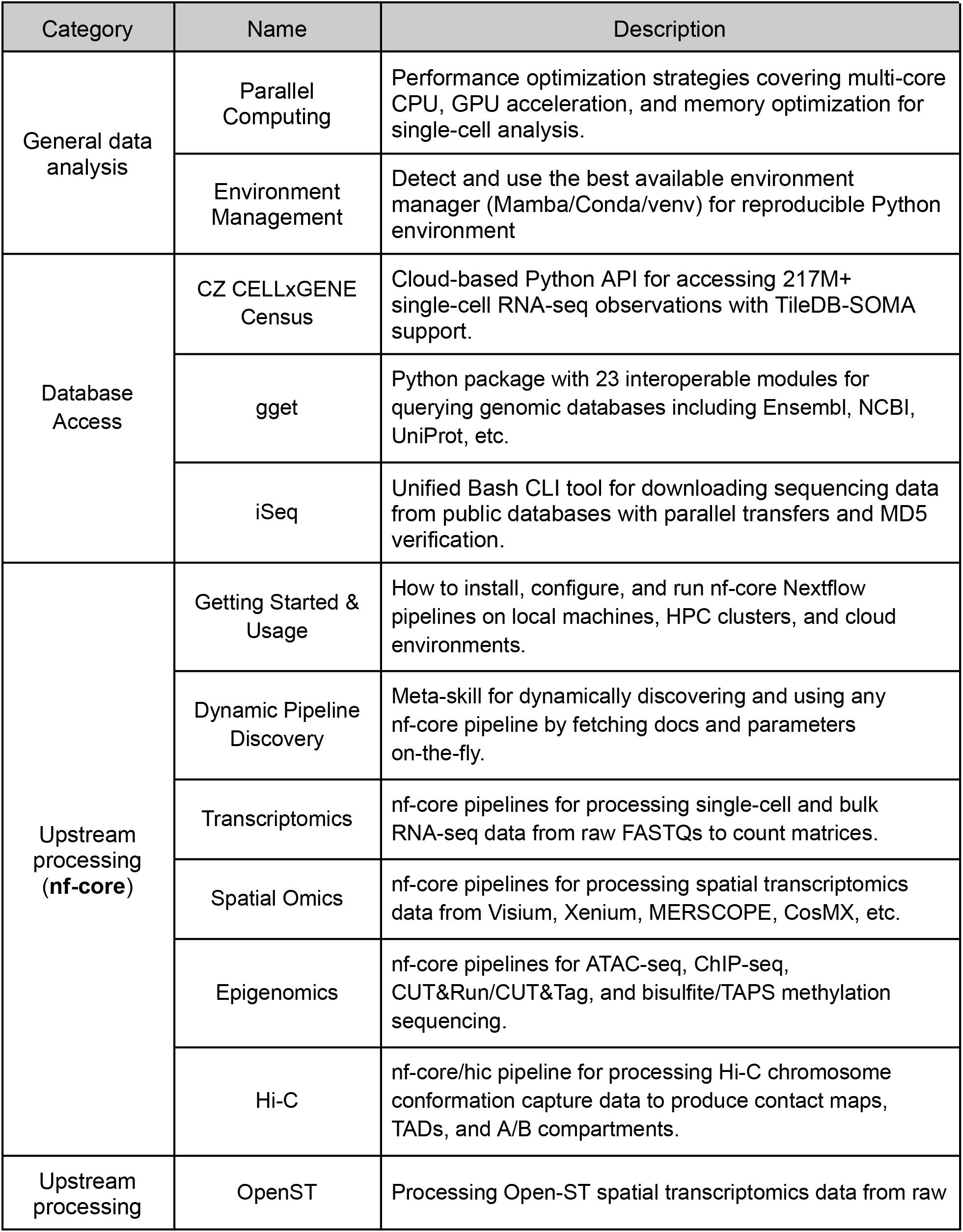
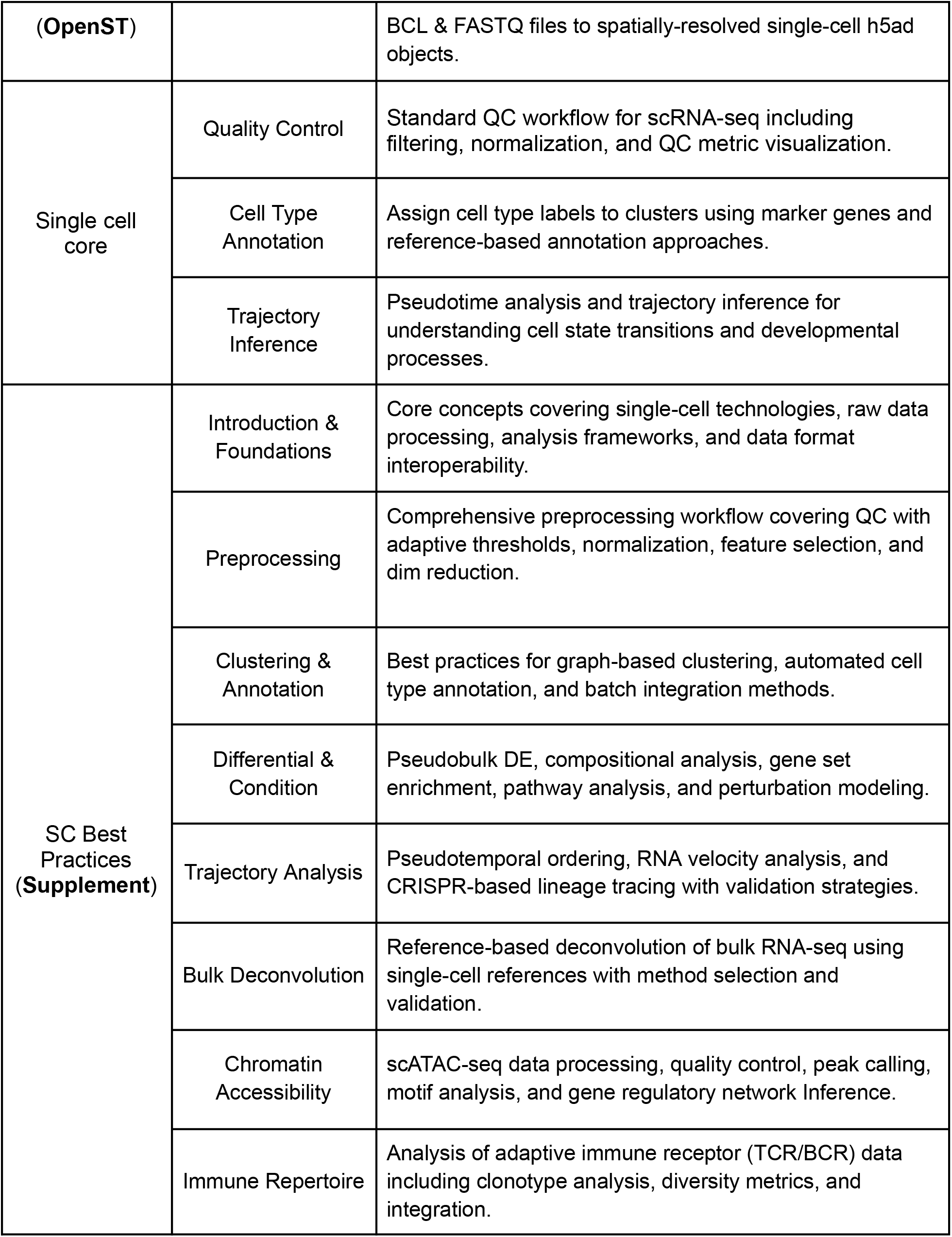
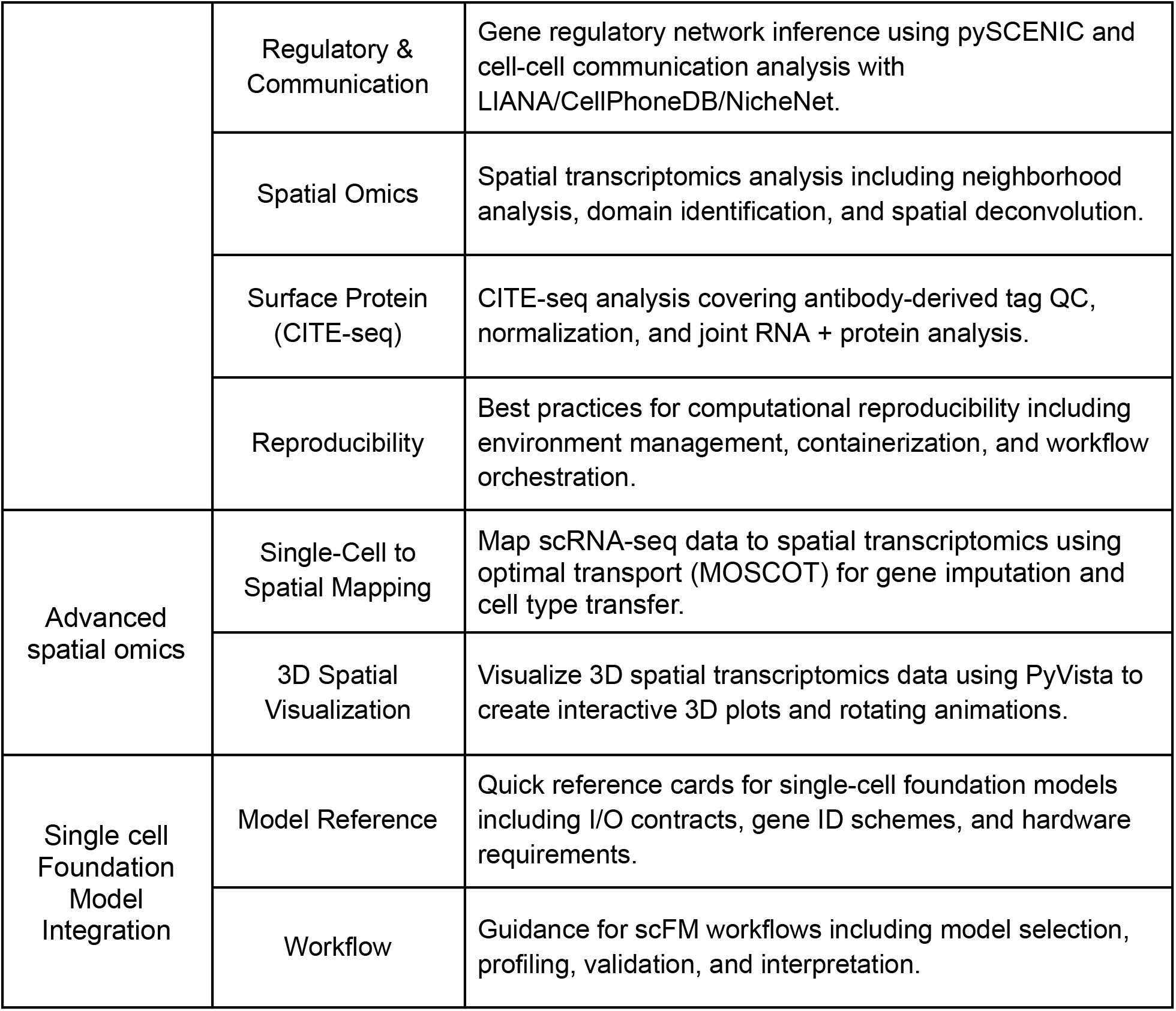
Builtin skills for data analyzer agent.

**Table S2.**
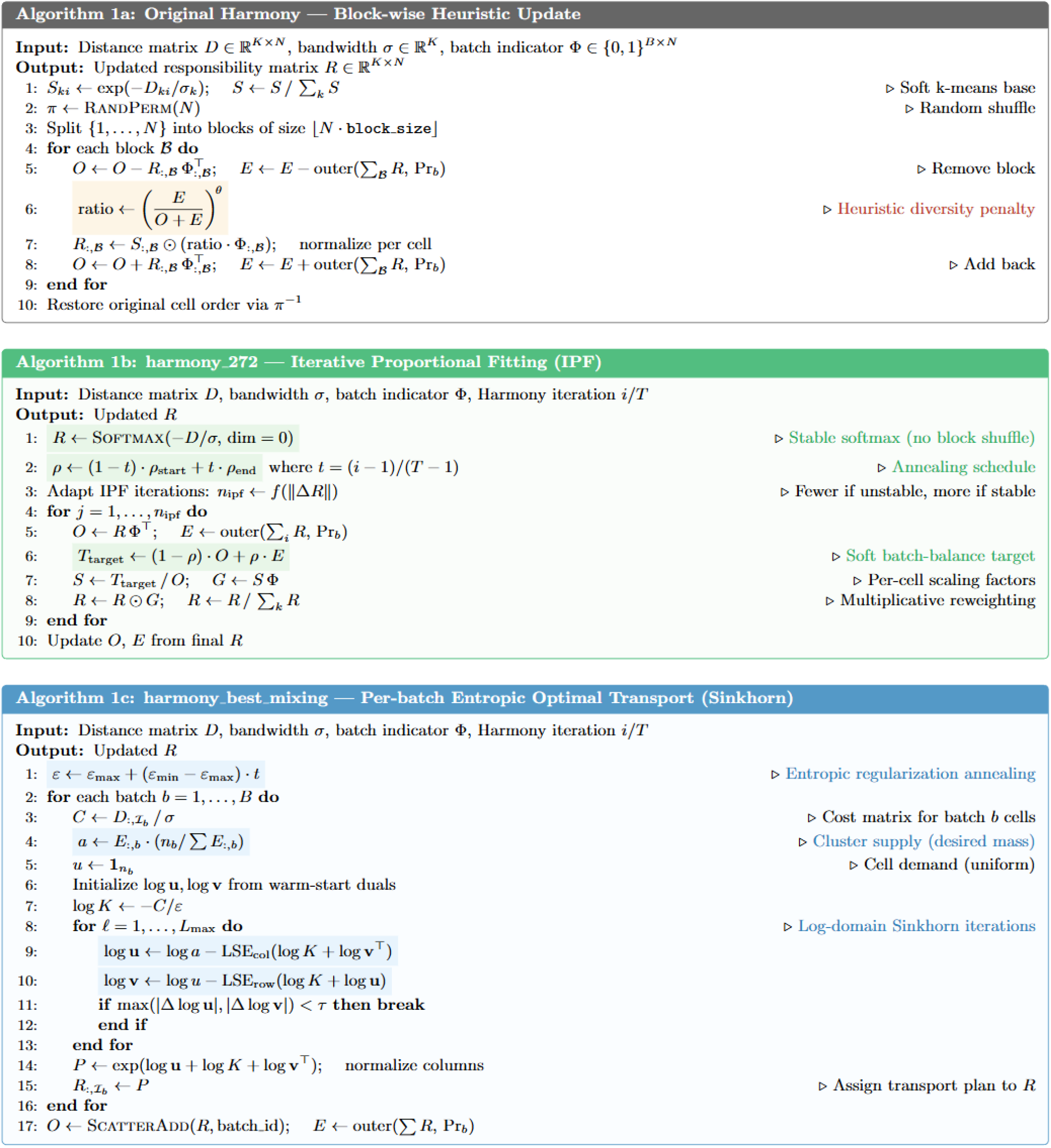
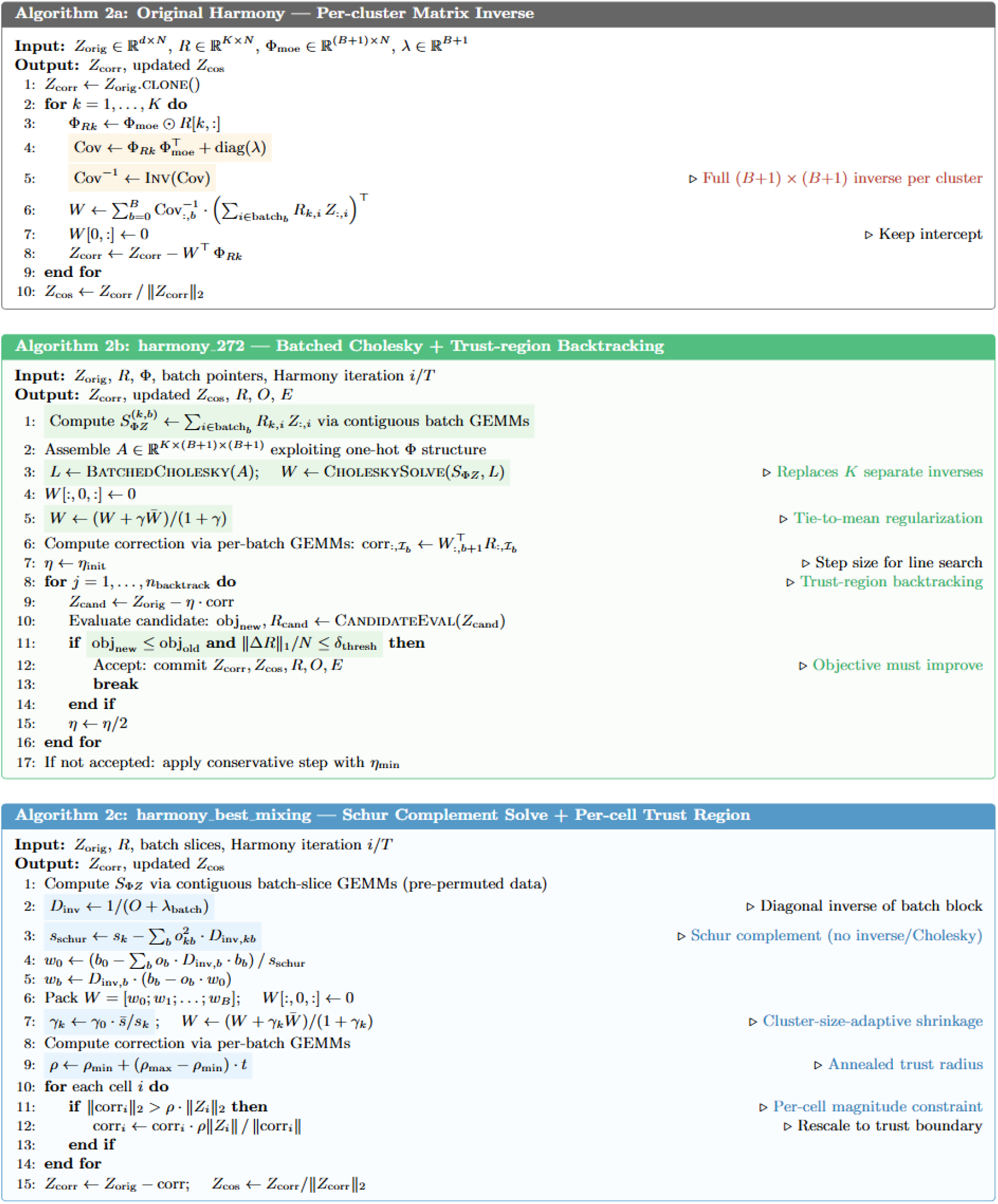
Pseudo-code comparison of two key parts of Harmony code evolution: soft cluster assignment (update_R) and batch effect correction (moe_correct_ridge). The original Harmony uses a block-wise heuristic ratio penalty for cluster assignment and per-cluster matrix inversion for ridge regression. Through automated code evolution, two variants independently emerged: harmony_272 replaces the block-wise heuristic with global Iterative Proportional Fitting (IPF) and introduces batched Cholesky solving with trust-region backtracking; harmony_best_mixing adopts per-batch entropic optimal transport (Sinkhorn) for hard batch-balance constraints and exploits the Schur complement structure for closed-form ridge solving with per-cell correction control. Key algorithmic changes are highlighted in color.

**Table S3.**
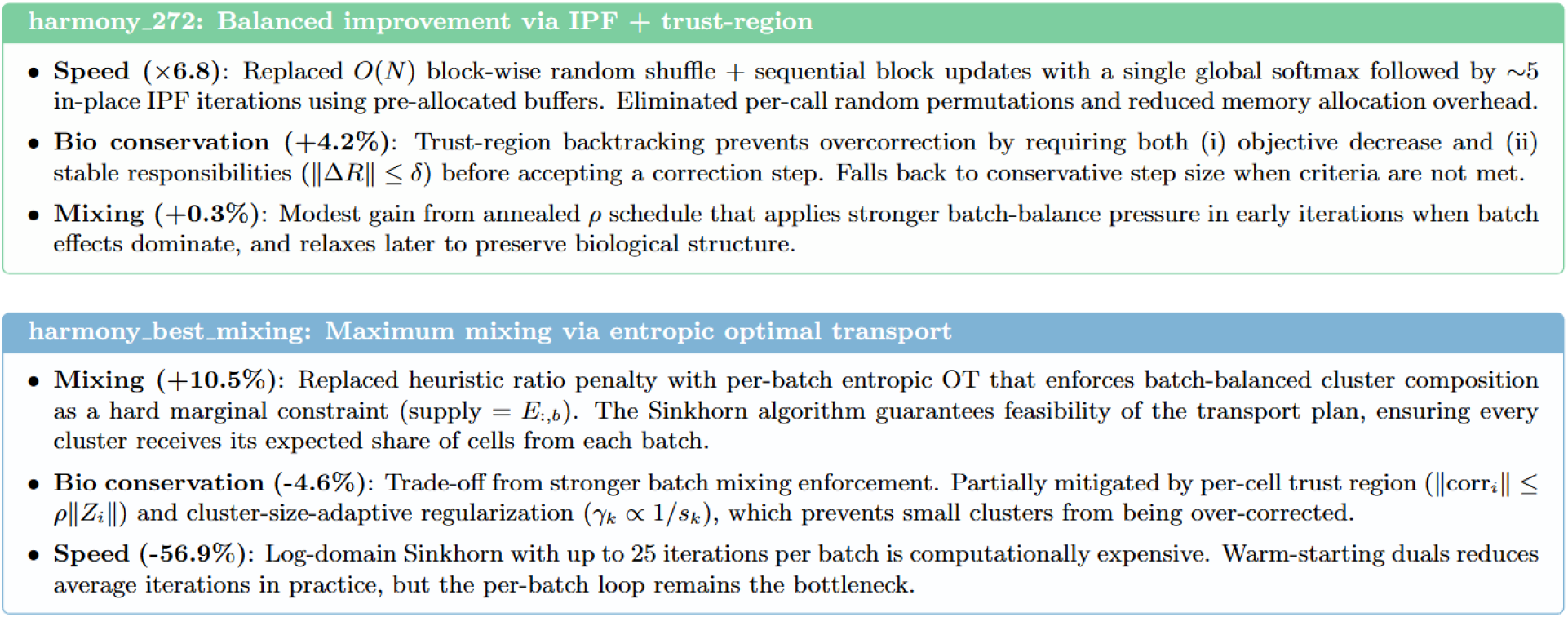
Analysis of the reasons for changes in algorithm performance in harmony evolution results.

**Table S4.**
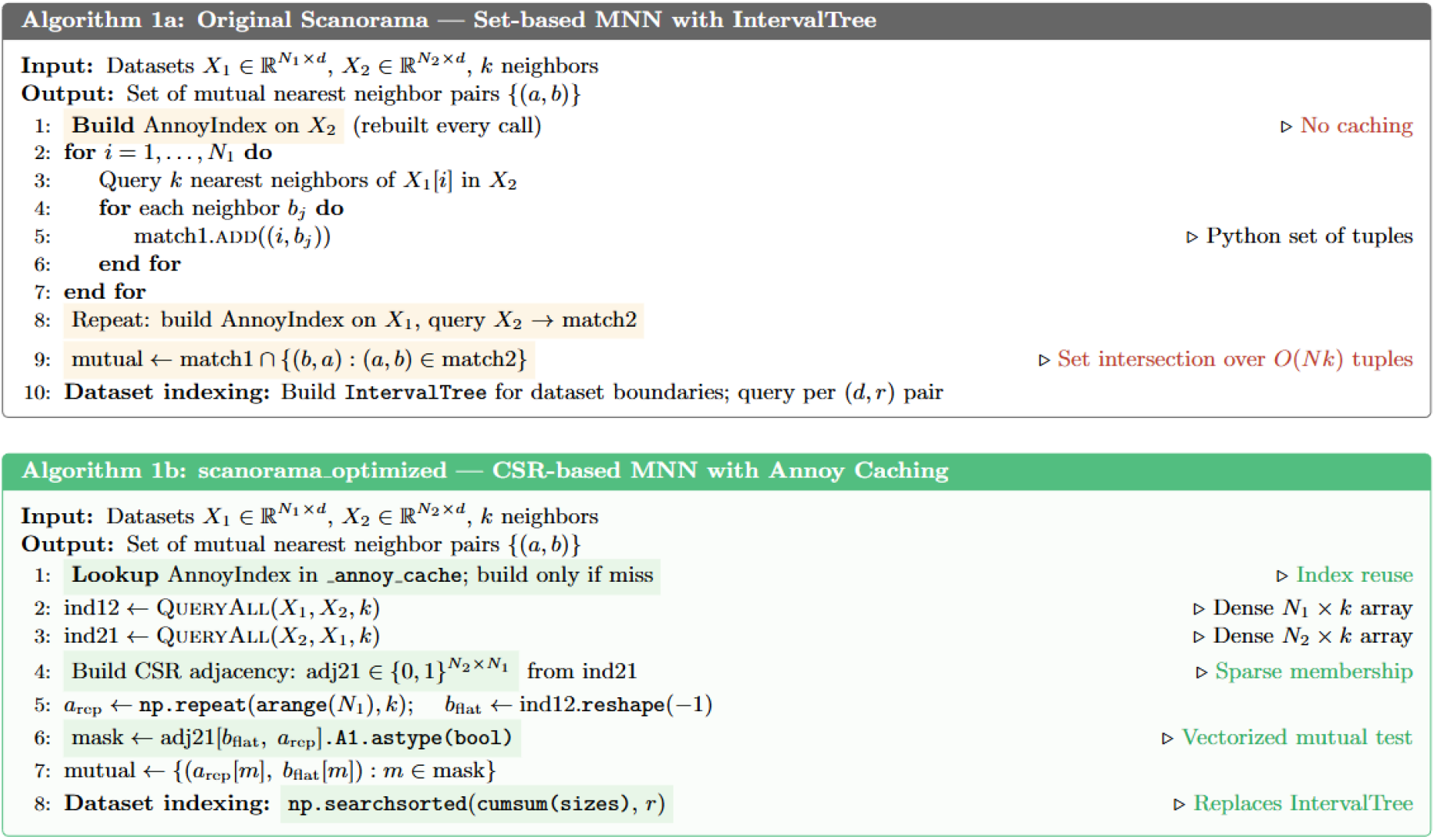
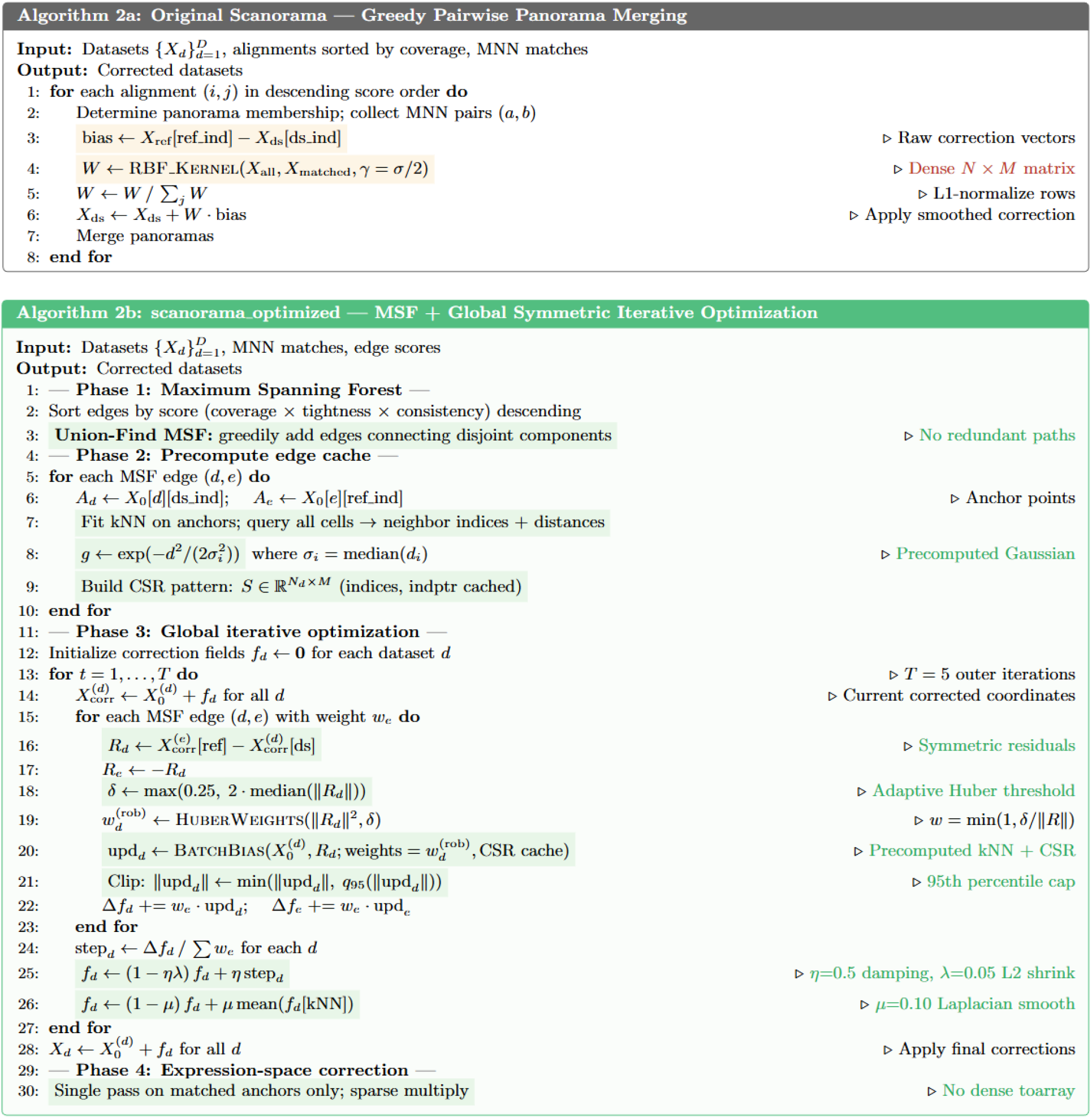
Pseudo-code comparison of two key parts of Scanorama code evolution: mutual nearest neighbor detection (mnn) and batch correction assembly (assemble). The original algorithm uses set-based MNN matching with greedy pairwise panorama merging, while the evolved variant introduces CSR-based vectorized MNN detection with Annoy index caching, and replaces the greedy merge with a Maximum Spanning Forest followed by global symmetric iterative optimization incorporating Huber robust weighting, update clipping, and Laplacian smoothing on the correction field. Key changes are highlighted in color.

**Table S5.**
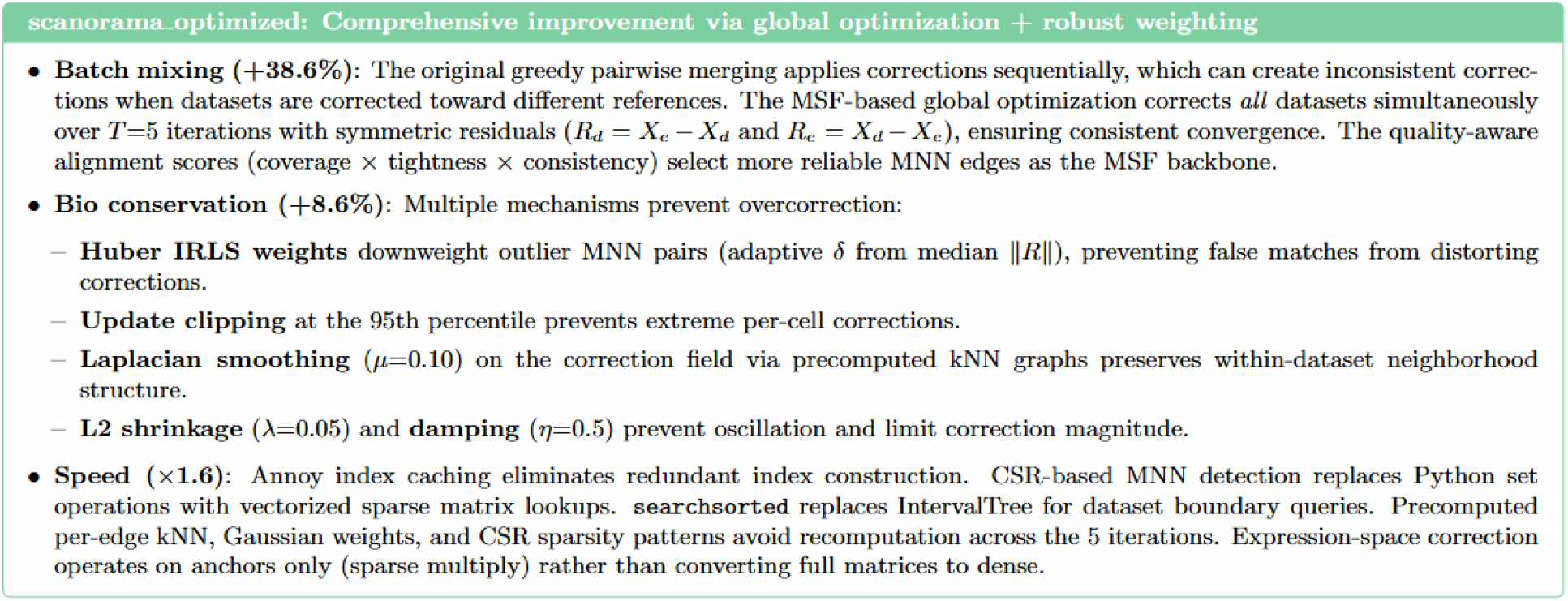
Analysis of the reasons for changes in algorithm performance in Scanorama evolution result.

**Table S6.**
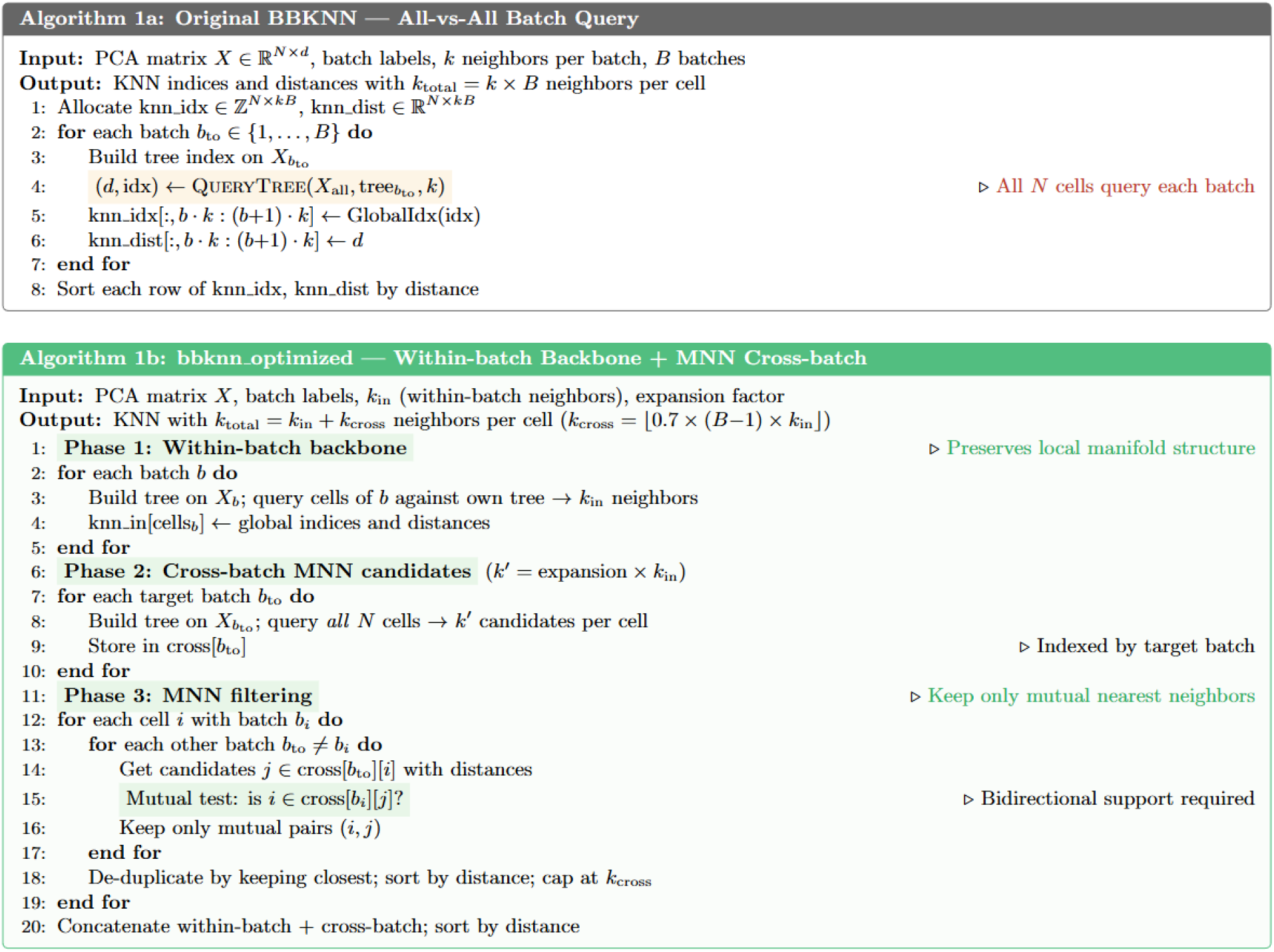
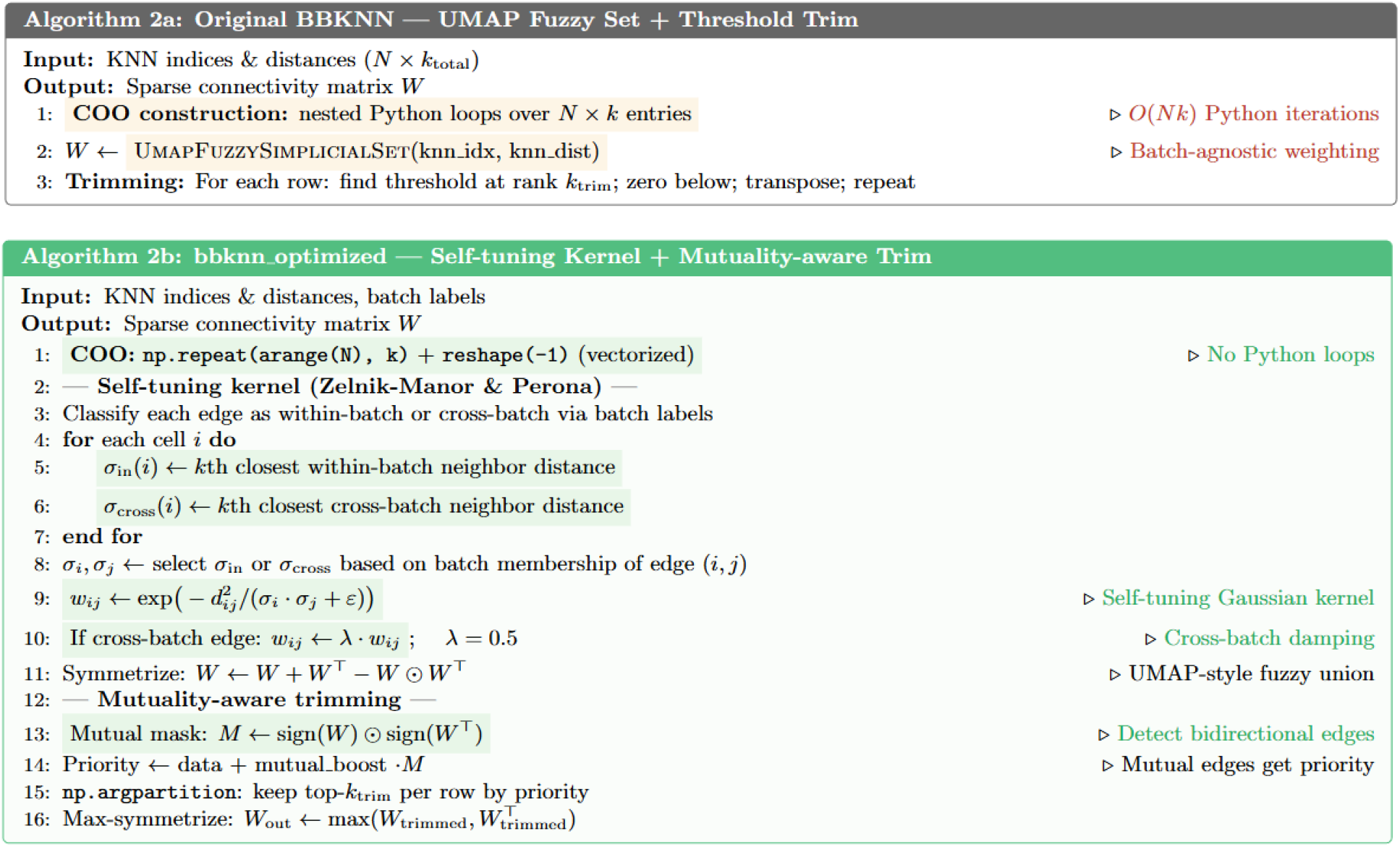
Pseudo-code comparison of two key parts of bbknn code evolution: KNN graph construction (get_graph) and connectivity weighting with graph trimming. The original algorithm queries all cells against every batch with batch-agnostic UMAP weighting, while the evolved variant introduces a within-batch backbone with MNN-filtered cross-batch edges, a self-tuning kernel with separate within/cross-batch bandwidth scales and cross-batch damping, and mutuality-aware trimming that preferentially preserves bidirectional edges. Key changes are highlighted in color.

**Table S7.**
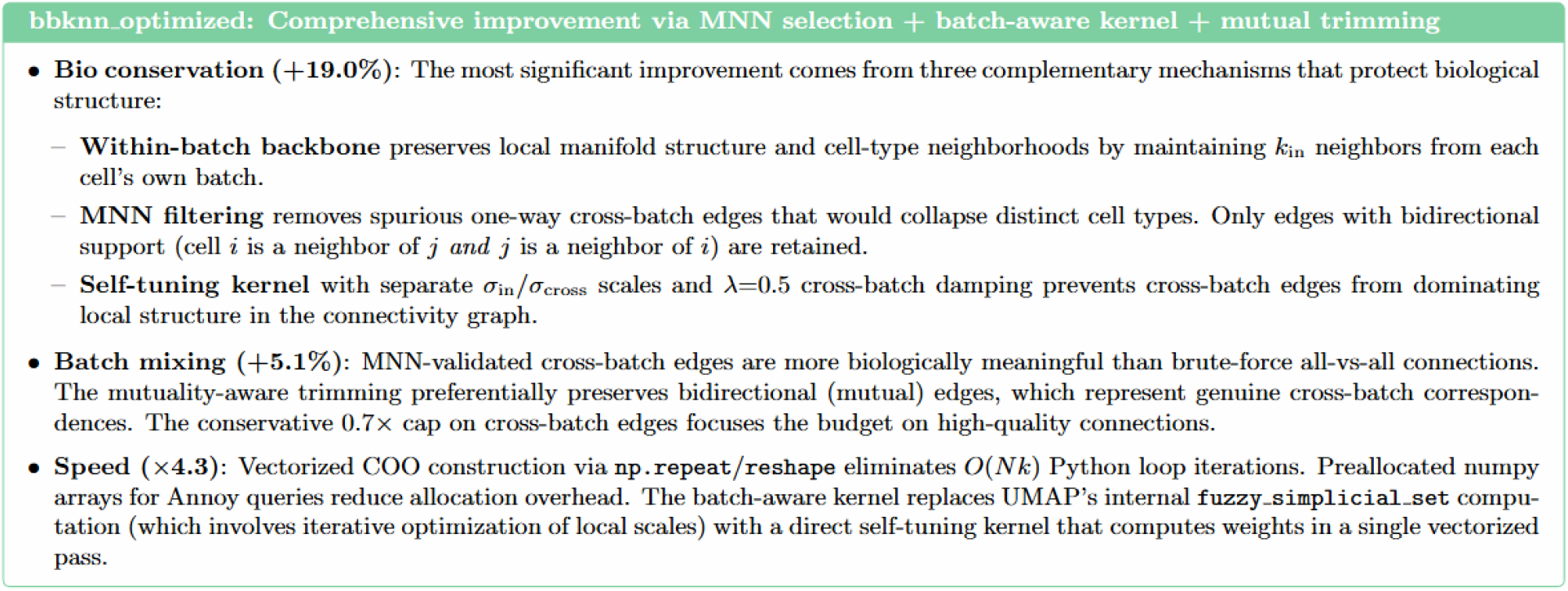
Analysis of the reasons for changes in algorithm performance in bbknn evolution result.

**Table S8.**
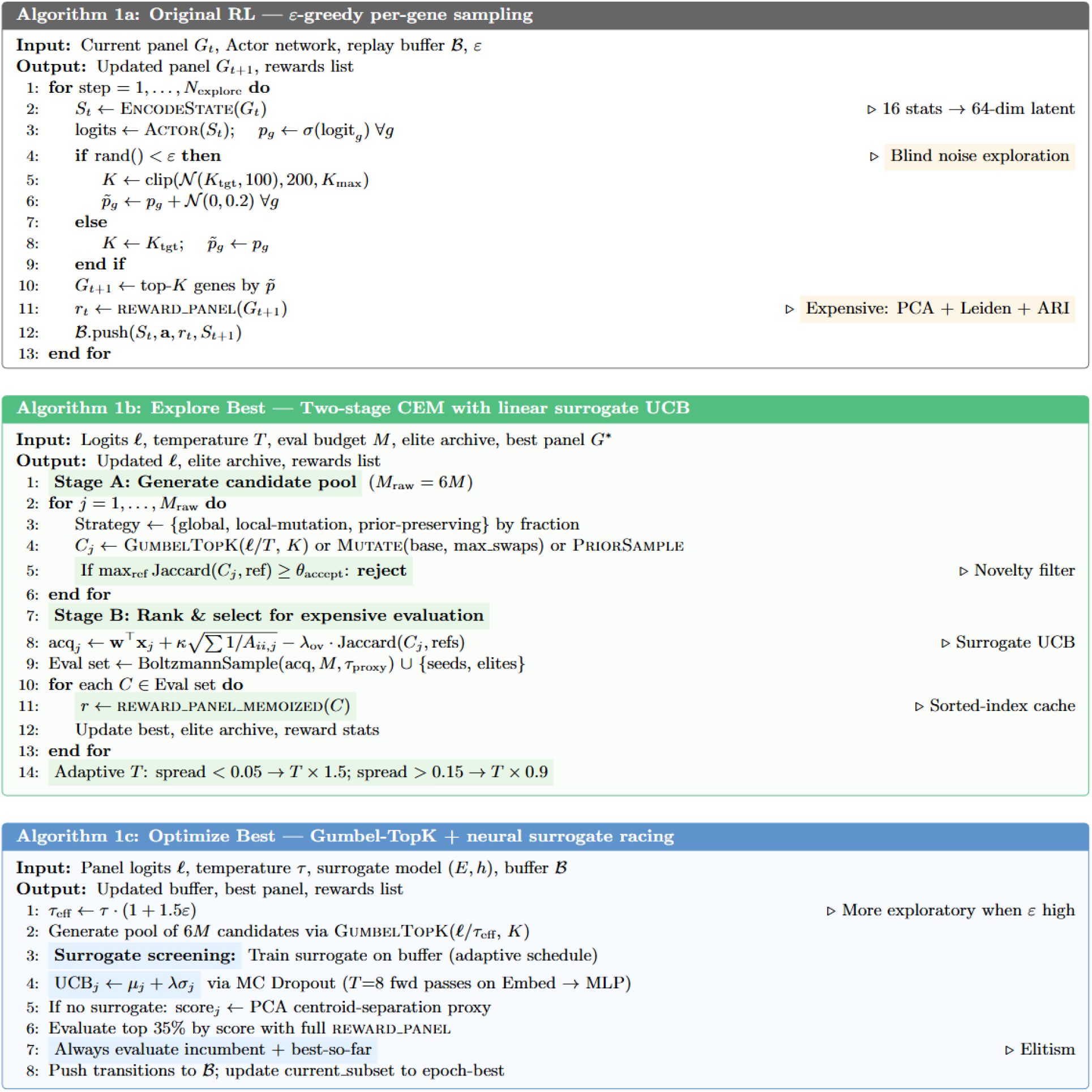
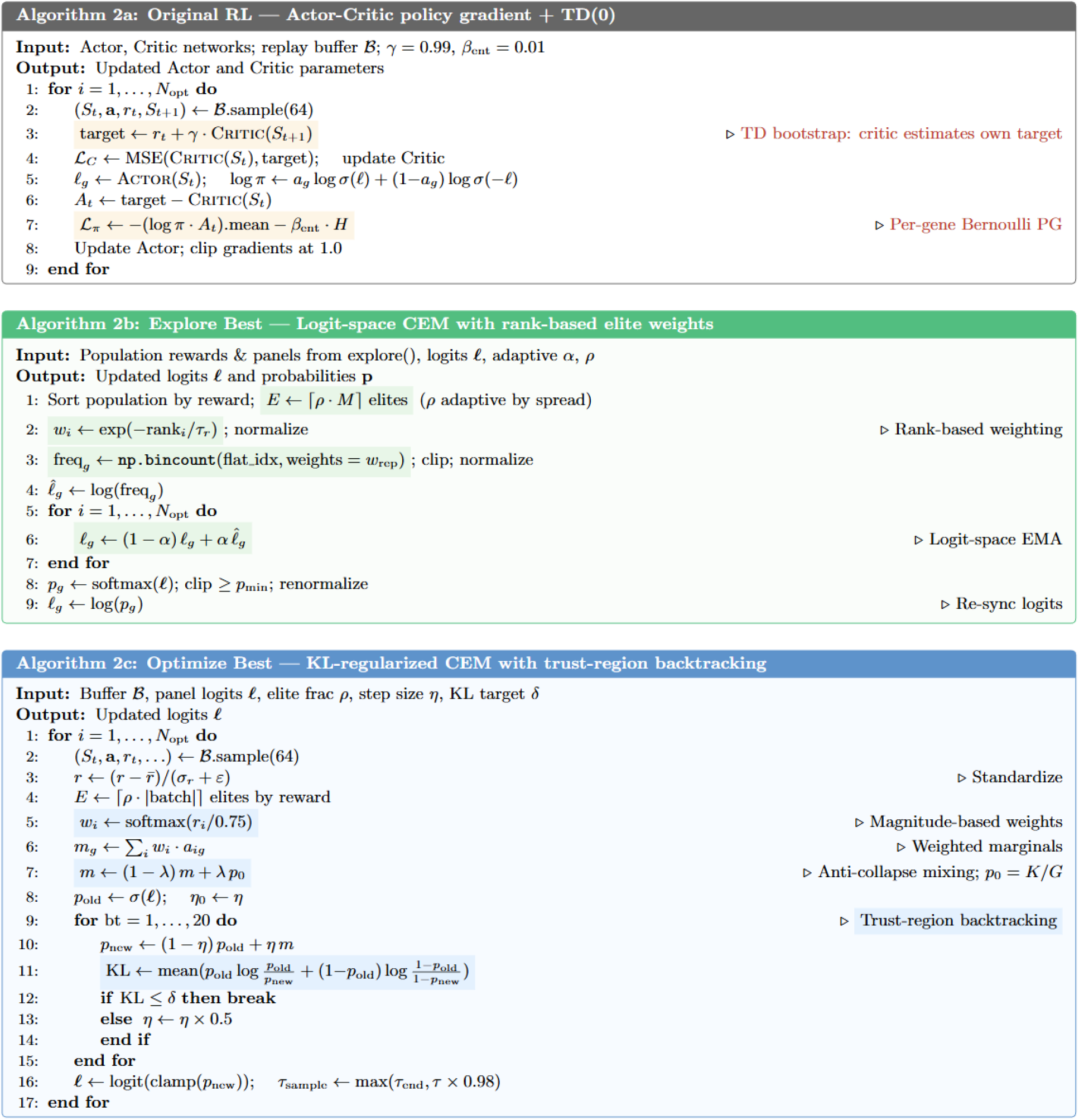
Pseudo-code comparison of two key parts of RL algorithm for panel design code evolution: exploration strategy (explore) and distribution update (optimize). The original RL actor-critic approach treats gene selection as sequential per-gene decisions with TD bootstrapping, while both evolved variants independently converge to CEM-based population search over complete panels, one using a linear surrogate with Jaccard novelty filtering and adaptive temperature kicks, the other using a neural surrogate with KL trust-region updates. Both also independently evolved multi-metric rewards and evaluation memoization. Key changes are highlighted in color.

**Table S9.**
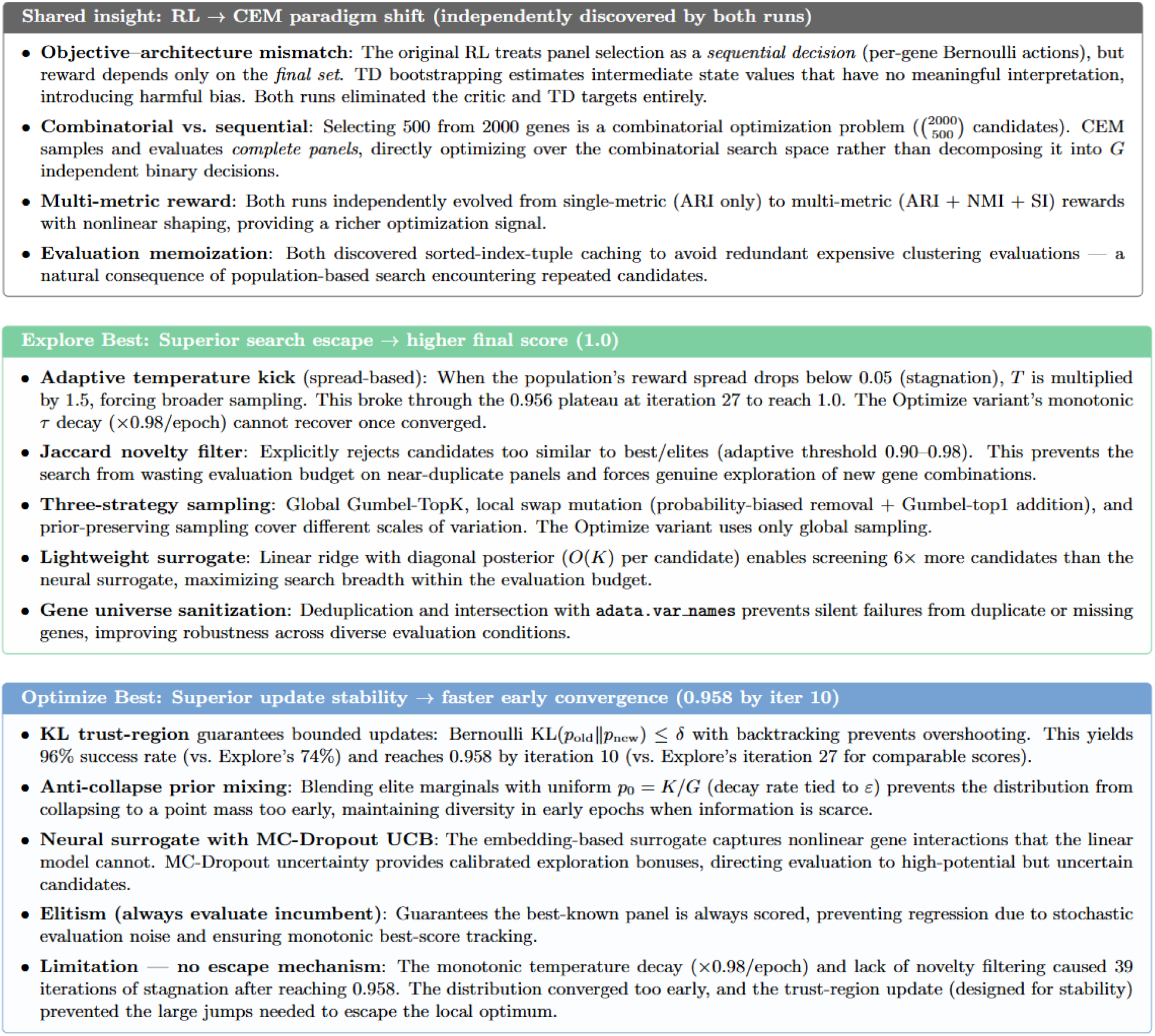
Analysis of the reasons for changes in algorithm performance in RL algorithm for panel design evolution result.

**Table S10.**
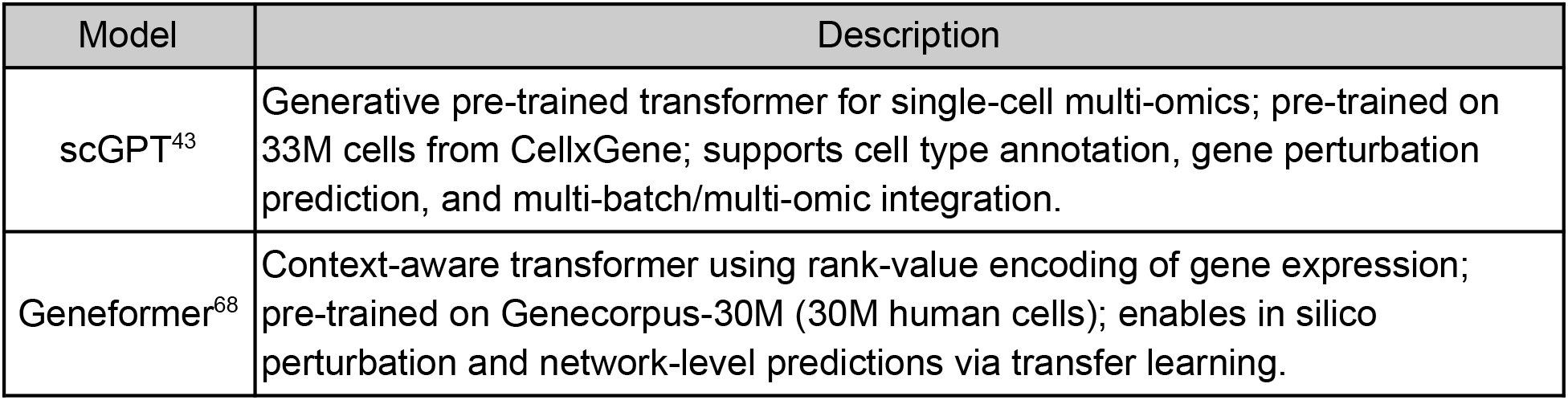
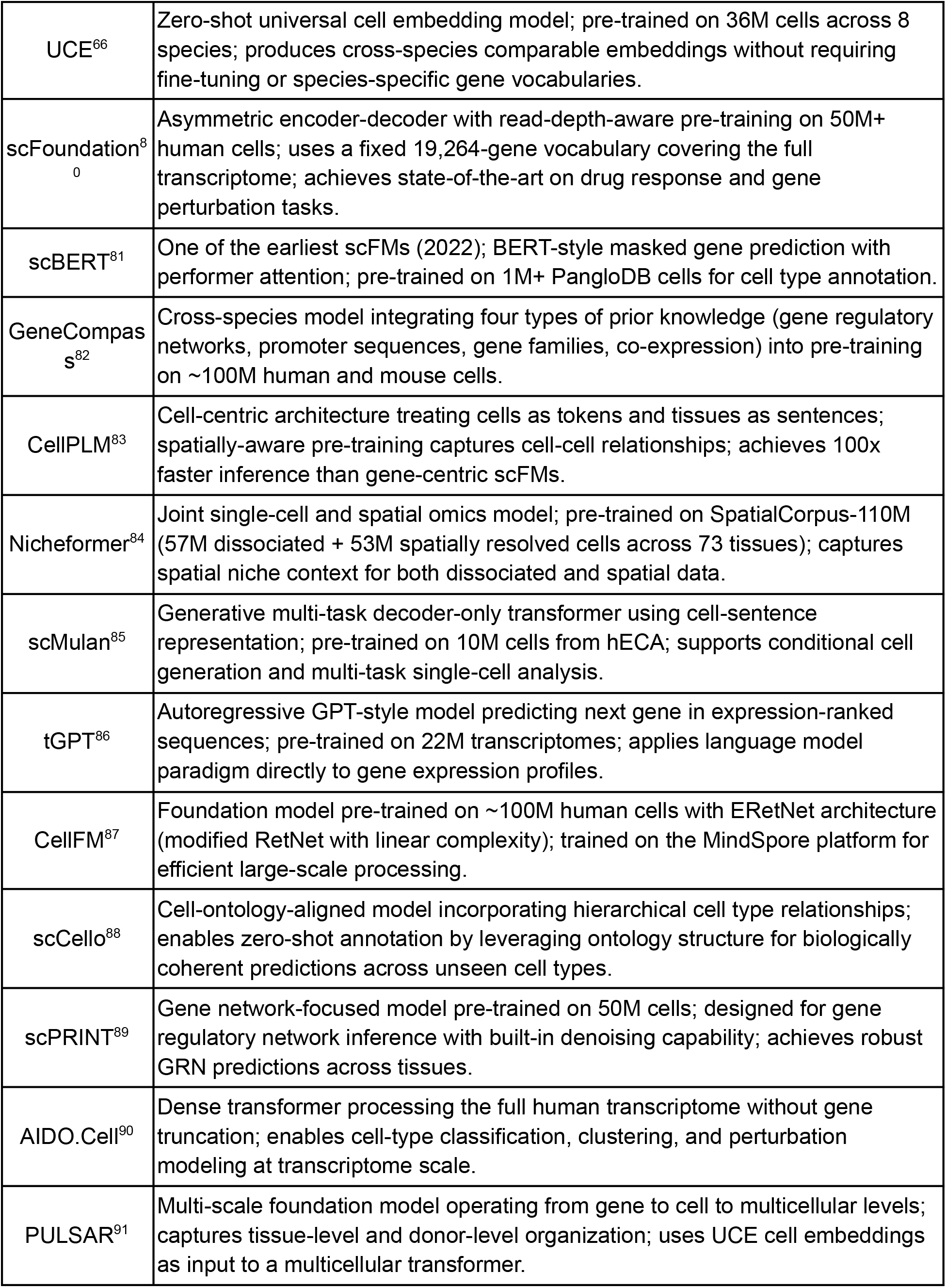
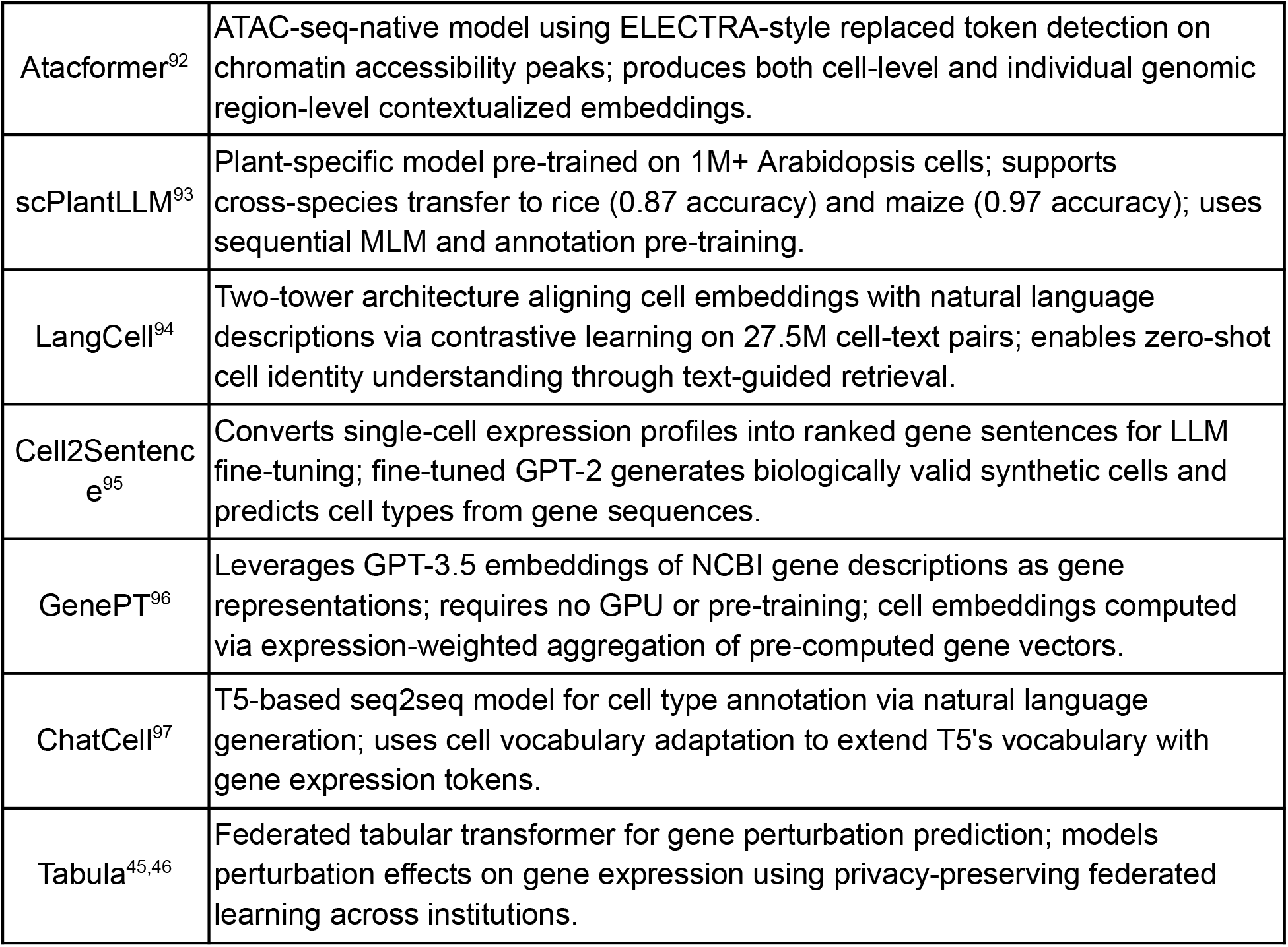
Single-cell foundation models integrated in the PantheonOS scFM module.

**Table S11.**
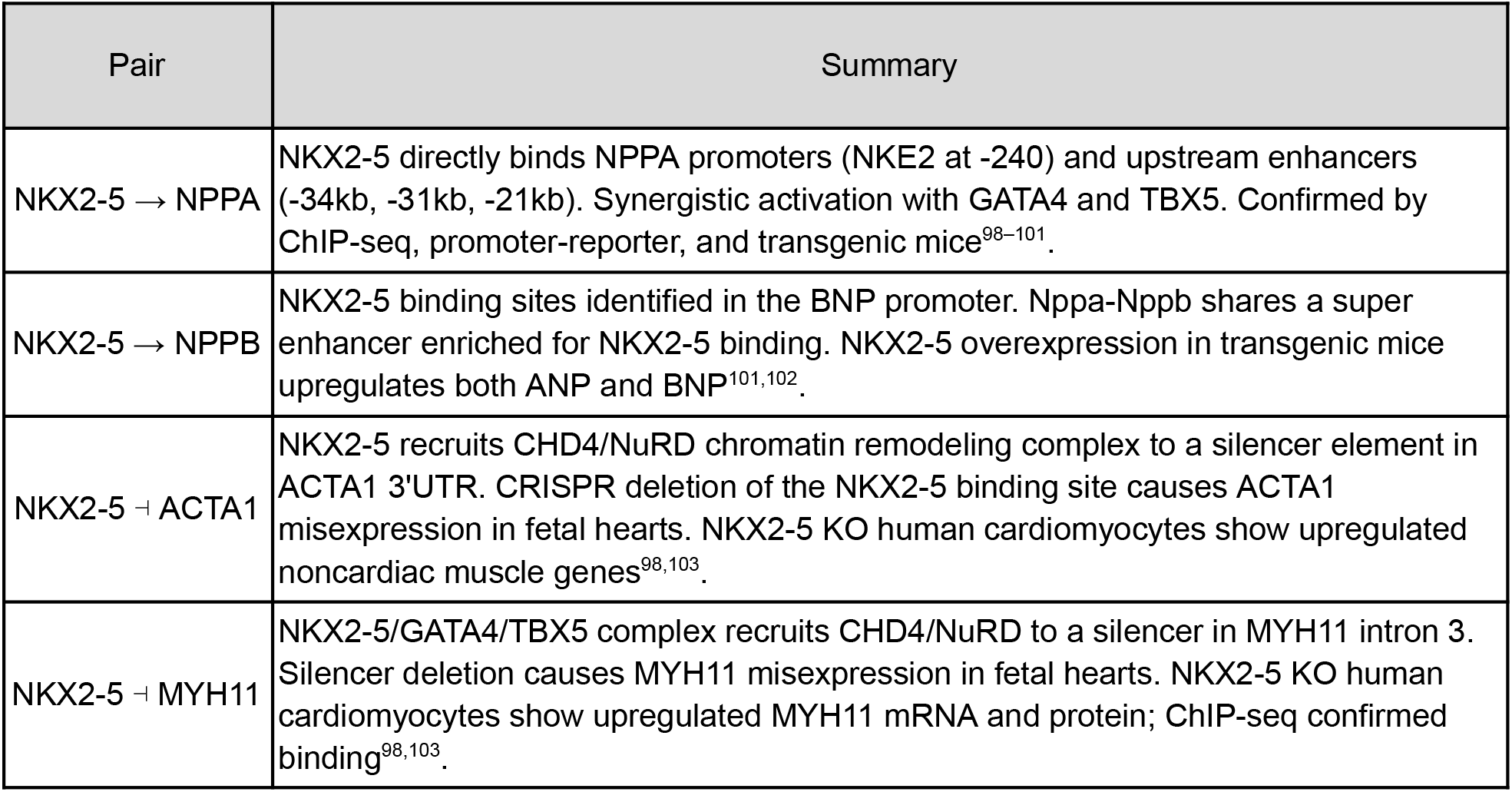
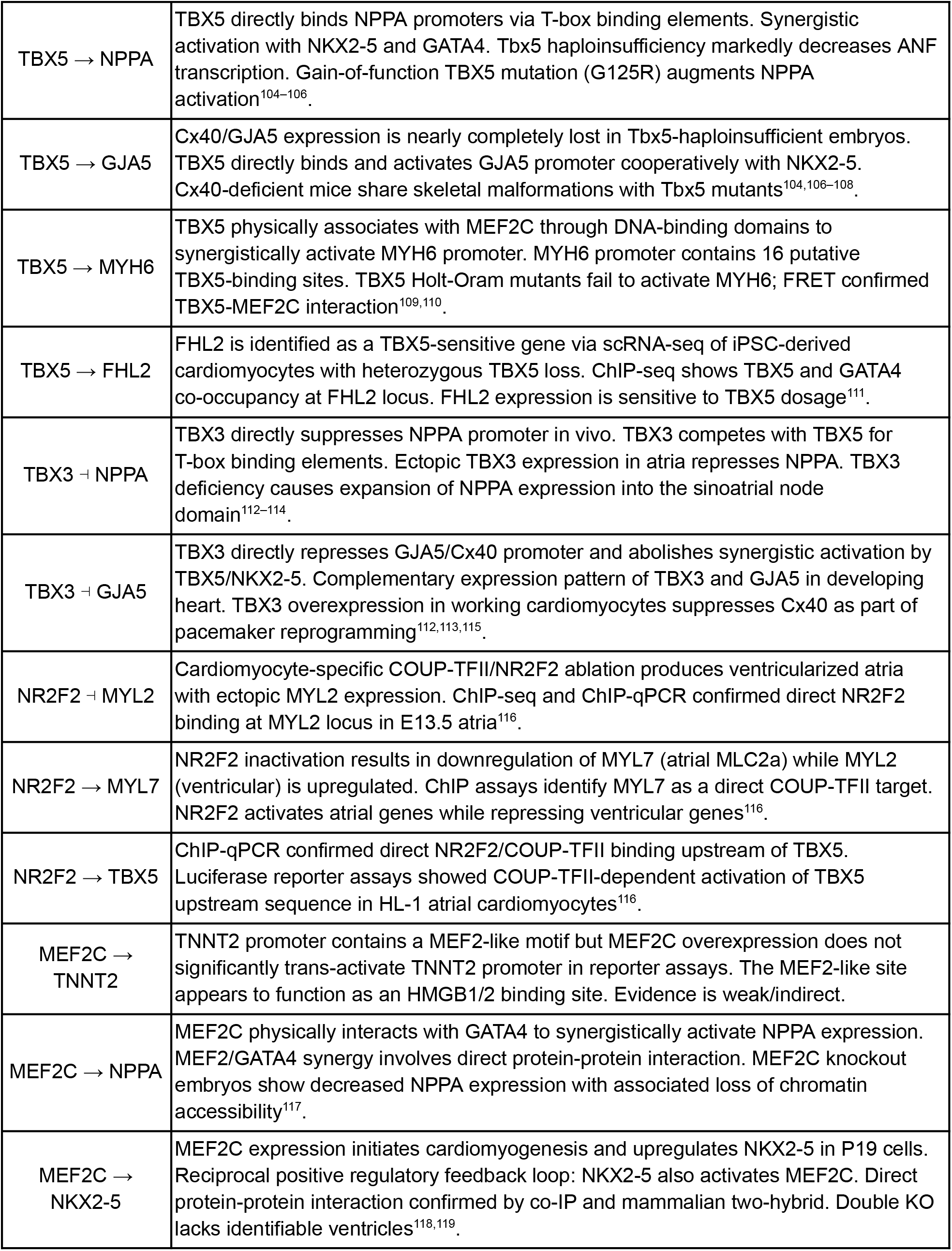
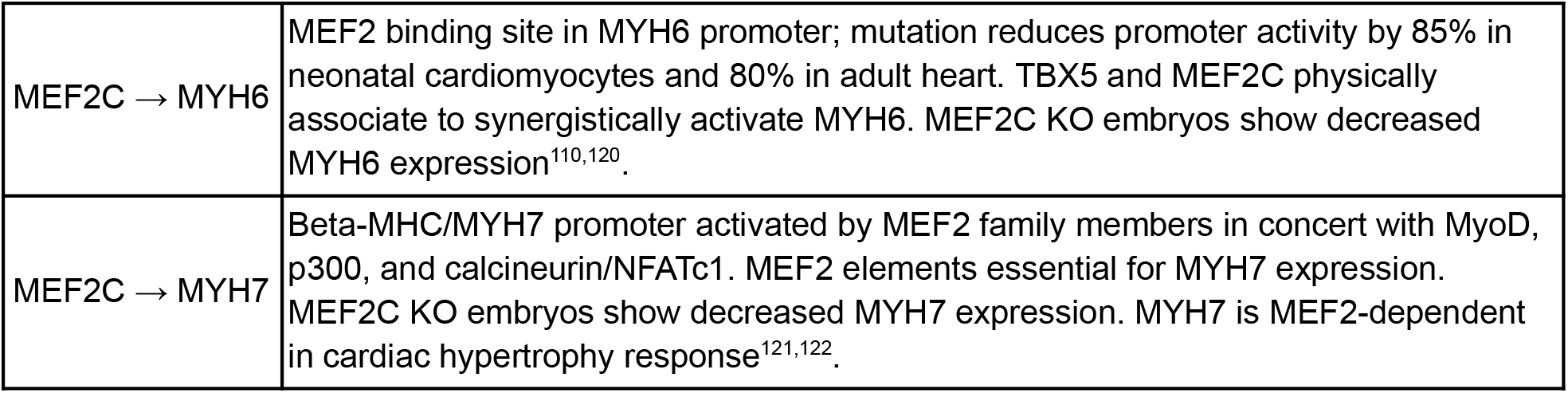
Literature-validated gene regulatory pairs used for in-silico GRN perturbation analysis.

## Reference

1. Yao, S. et al. ReAct: Synergizing Reasoning and Acting in Language Models. (2022).

2. Gao, S. et al. Empowering biomedical discovery with AI agents.,Cell 187, 6125–6151 (2024).

3. Lin, Z. et al. Spatial transcriptomics AI agent charts hPSC-pancreas maturation., bioRxiv (2025) doi:10.1101/2025.04.01.646731.

4. Wang, H. et al. SpatialAgent: An autonomous AI agent for spatial biology., bioRxiv 2025.04.03.646459 (2025) doi:10.1101/2025.04.03.646459.

5. Alber, S. et al. CellVoyager: AI CompBio Agent Generates New Insights by Autonomously Analyzing Biological Data., bioRxiv 2025.06.03.657517 (2025) doi:10.1101/2025.06.03.657517.

6. Huang, K. et al. Biomni: A General-Purpose Biomedical AI Agent., bioRxiv 2025.05.30.656746 (2025) doi:10.1101/2025.05.30.656746.

7. Korsunsky, I. et al. Fast, sensitive and accurate integration of single-cell data with Harmony., Nat Methods 16, 1289–1296 (2019).

8. Hie, B., Bryson, B. & Berger, B. Efficient integration of heterogeneous single-cell transcriptomes using Scanorama. Nature Biotechnology 37, 685–691 (2019).

9. Polański, K. et al. BBKNN: fast batch alignment of single cell transcriptomes., Bioinformatics 36, 964–965 (2020).

10. Bunne, C. et al. How to build the virtual cell with artificial intelligence: Priorities and opportunities., Cell 187, 7045–7063 (2024).

11. GitHub - langchain-ai/langchain: The platform for reliable agents. GitHub https://github.com/langchain-ai/langchain.

12. GitHub - microsoft/autogen: A programming framework for agentic AI. GitHub https://github.com/microsoft/autogen.

13. GitHub - crewAIInc/crewAI: Framework for orchestrating role-playing, autonomous AI agents. By fostering collaborative intelligence, CrewAI empowers agents to work together seamlessly, tackling complex tasks. GitHub https://github.com/crewAIInc/crewAI.

14. Website. https://openai.com/index/introducing-openai-frontier/.

15. Xiao, Y. et al. CellAgent: LLM-Driven Multi-Agent Framework for Natural Language-Based Single-Cell Analysis., bioRxiv 2024.05.13.593861 (2025) doi:10.1101/2024.05.13.593861.

16. Wang, J., Wang, J., Athiwaratkun, B., Zhang, C. & Zou, J. Mixture-of-Agents Enhances Large Language Model Capabilities. (2024).

17. Kuehl, M. et al. BioContextAI is a community hub for agentic biomedical systems., Nat Biotechnol 43, 1755–1757 (2025).

18. Kuemmerle, L. B. et al. Probe set selection for targeted spatial transcriptomics., Nat Methods 21, 2260–2270 (2024).

19. Dumitrascu, B., Villar, S., Mixon, D. G. & Engelhardt, B. E. Optimal marker gene selection for cell type discrimination in single cell analyses. Nat Commun 12, 1186 (2021).

20. Wolf, F. A., Angerer, P. & Theis, F. J. SCANPY: large-scale single-cell gene expression data analysis. Genome Biol 19, 15 (2018).

21. Kaelbling, L. P., Littman, M. L. & Moore, A. W. Reinforcement Learning: A Survey. jair 4, 237–285 (1996).

22. Stelzer, G. et al. The GeneCards suite: From gene data mining to disease genome sequence analyses., Curr. Protoc. Bioinformatics 54, 1.30.1–1.30.33 (2016).

23. Bairoch, A. et al. The Universal Protein Resource (UniProt)., Nucleic Acids Res. 33, D154–9 (2005).

24. Novikov, A. et al. AlphaEvolve: A coding agent for scientific and algorithmic discovery. (2025).

25. GitHub - algorithmicsuperintelligence/openevolve: Open-source implementation of AlphaEvolve. GitHub https://github.com/algorithmicsuperintelligence/openevolve.

26. Lange, R. T., Imajuku, Y. & Cetin, E. ShinkaEvolve: Towards Open-Ended And Sample-Efficient Program Evolution. (2025).

27. Mouret, J.-B. & Clune, J. Illuminating search spaces by mapping elites. (2015).

28. Büttner, M., Miao, Z., Wolf, F. A., Teichmann, S. A. & Theis, F. J. A test metric for assessing single-cell RNA-seq batch correction. Nat Methods 16, 43–49 (2019).

29. An AI system to help scientists write expert-level empirical software. https://arxiv.org/html/2509.06503v1.

30. Luecken, M. D. et al. Defining and benchmarking open problems in single-cell analysis., Nat Biotechnol 43, 1035–1040 (2025).

31. Schott, M. et al. Protocol for high-resolution 3D spatial transcriptomics using Open-ST., STAR Protoc 6, 103521 (2025).

32. Qiu, X. et al. Spatiotemporal modeling of molecular holograms., Cell 188, 1744 (2025).

33. Zeng, Z. et al. OmicVerse: a framework for bridging and deepening insights across bulk and single-cell sequencing., Nat Commun 15, 5983 (2024).

34. Qiu, C. et al. Systematic reconstruction of cellular trajectories across mouse embryogenesis., Nat Genet 54, 328–341 (2022).

35. Biancalani, T. et al. Deep learning and alignment of spatially resolved single-cell transcriptomes with Tangram., Nat Methods 18, 1352–1362 (2021).

36. Torres-Padilla, M.-E. et al. The anterior visceral endoderm of the mouse embryo is established from both preimplantation precursor cells and by de novo gene expression after implantation., Dev Biol 309, 97–112 (2007).

37. Katoh, M. & Katoh, M. CER1 is a common target of WNT and NODAL signaling pathways in human embryonic stem cells. Int. J. Mol. Med. 17, 795–799 (2006).

38. Rosa Ma, X. et al. Molecular convergence of risk variants for congenital heart defects leveraging a regulatory map of the human fetal heart., medRxiv 2024.11.20.24317557 (2024) doi:10.1101/2024.11.20.24317557.

39. Kern, C. et al. MERFISH+, a large-scale, multi-omics spatial technology resolves the molecular holograms of the 3D human developing heart., bioRxiv (2025) doi:10.1101/2025.11.02.686137.

40. Yang, A. et al. CHDgene: A Curated Database for Congenital Heart Disease Genes., Circ Genom Precis Med 15, e003539 (2022).

41. Klein, D. et al. Mapping cells through time and space with moscot., Nature 638, 1065–1075 (2025).

42. Sheth, M. U. et al. Mapping enhancer-gene regulatory interactions from single-cell data., bioRxiv 2024.11.23.624931 (2024) doi:10.1101/2024.11.23.624931.

43. Cui, H. et al. scGPT: toward building a foundation model for single-cell multi-omics using generative AI., Nat Methods 21, 1470–1480 (2024).

44. McCulley, D. J. & Black, B. L. Transcription factor pathways and congenital heart disease. Current topics in developmental biology 100, (2012).

45. Ding, J. et al. Toward a privacy-preserving predictive foundation model of single-cell transcriptomics with federated learning and tabular modeling., bioRxiv 2025.01.06.631427 (2025) doi:10.1101/2025.01.06.631427.

46. Ding, J. et al. Tabula: A Tabular Self-Supervised Foundation Model for Single-Cell Transcriptomics. in The Thirty-ninth Annual Conference on Neural Information Processing Systems (2025).

47. Munshi, N. V. Gene Regulatory Networks in Cardiac Conduction System Development. Circulation Research (2012) doi:10.1161/CIRCRESAHA.111.260026.

48. Li, W., Lin, Y., Xia, M. & Jin, C. Rethinking Mixture-of-Agents: Is Mixing Different Large Language Models Beneficial? (2025).

49. Luebbert, L. & Pachter, L. Efficient querying of genomic reference databases with gget. Bioinformatics 39, (2023).

50. Chao, H., Li, Z., Chen, D. & Chen, M. iSeq: an integrated tool to fetch public sequencing data. Bioinformatics 40, (2024).

51. CZI Cell Science Program et al. CZ CELLxGENE Discover: a single-cell data platform for scalable exploration, analysis and modeling of aggregated data., Nucleic Acids Res 53, D886–D900 (2025).

52. UniProt Consortium. UniProt: the Universal Protein Knowledgebase in 2023. Nucleic Acids Res 51, D523–D531 (2023).

53. Kuleshov, M. V. et al. Enrichr: a comprehensive gene set enrichment analysis web server 2016 update., Nucleic Acids Res 44, W90–7 (2016).

54. Ewels, P. A. et al. The nf-core framework for community-curated bioinformatics pipelines., Nat Biotechnol 38, 276–278 (2020).

55. Chen, K. H., Boettiger, A. N., Moffitt, J. R., Wang, S. & Zhuang, X. RNA imaging. Spatially resolved, highly multiplexed RNA profiling in single cells. Science 348, aaa6090 (2015).

56. Janesick, A. et al. High resolution mapping of the tumor microenvironment using integrated single-cell, spatial and in situ analysis., Nat Commun 14, 8353 (2023).

57. Young, M. D. & Behjati, S. SoupX removes ambient RNA contamination from droplet-based single-cell RNA sequencing data. Gigascience 9, (2020).

58. Yang, S. et al. Decontamination of ambient RNA in single-cell RNA-seq with DecontX., Genome Biol 21, 57 (2020).

59. Wolock, S. L., Lopez, R. & Klein, A. M. Scrublet: Computational Identification of Cell Doublets in Single-Cell Transcriptomic Data. Cell Syst 8, 281–291.e9 (2019).

60. McGinnis, C. S., Murrow, L. M. & Gartner, Z. J. DoubletFinder: Doublet Detection in Single-Cell RNA Sequencing Data Using Artificial Nearest Neighbors. Cell Syst 8, 329–337.e4 (2019).

61. Xu, C. et al. Automatic cell-type harmonization and integration across Human Cell Atlas datasets., Cell 186, 5876–5891.e20 (2023).

62. Xu, C. et al. Probabilistic harmonization and annotation of single-cell transcriptomics data with deep generative models., Mol Syst Biol 17, e9620 (2021).

63. Haghverdi, L., Büttner, M., Wolf, F. A., Buettner, F. & Theis, F. J. Diffusion pseudotime robustly reconstructs lineage branching. Nat Methods 13, 845–848 (2016).

64. Bergen, V., Lange, M., Peidli, S., Wolf, F. A. & Theis, F. J. Generalizing RNA velocity to transient cell states through dynamical modeling. Nat Biotechnol 38, 1408–1414 (2020).

65. Bane Sullivan, C. & Kaszynski, A. A. PyVista: 3D plotting and mesh analysis through a streamlined interface for the Visualization Toolkit (VTK). Journal of Open Source Software 4, 1450 (2019).

66. Rosen, Y. et al. Universal Cell Embeddings: A Foundation Model for Cell Biology., bioRxiv 2023.11.28.568918 (2024) doi:10.1101/2023.11.28.568918.

67. Magnusson, R. et al. The human metabolome and machine learning improves predictions of the post-mortem interval., Nat Commun 17, 1504 (2026).

68. Theodoris, C. V. et al. Transfer learning enables predictions in network biology., Nature 618, 616–624 (2023).

69. Zhang, Q. et al. Agentic Context Engineering: Evolving Contexts for Self-Improving Language Models. (2025).

70. Xiao, M. et al. Knowledge-guided gene panel selection for label-free single-cell RNA-seq data: A reinforcement learning perspective., IEEE Trans. Comput. Biol. Bioinform. 22, 3041–3054 (2025).

71. Lambora, A., Gupta, K. & Chopra, K. Genetic algorithm-A literature review. in 2019 International Conference on Machine Learning, Big Data, Cloud and Parallel Computing (COMITCon) (IEEE, 2019). doi:10.1109/comitcon.2019.8862255.

72. Romera-Paredes, B. et al. Mathematical discoveries from program search with large language models., Nature 625, 468–475 (2023).

73. Okamura, D., Hayashi, K. & Matsui, Y. Mouse epiblasts change responsiveness to BMP4 signal required for PGC formation through functions of extraembryonic ectoderm. Mol Reprod Dev 70, 20–29 (2005).

74. Kirillov, A. et al. Segment Anything. (2023).

75. Edstedt, J., Sun, Q., Bökman, G., Wadenbäck, M. & Felsberg, M. RoMa: Robust Dense Feature Matching. (2023).

76. Random sample consensus. Communications of the ACM (1981) doi:10.1145/358669.358692.

77. Srinivas, S. The anterior visceral endoderm-turning heads. Genesis 44, 565–572 (2006).

78. Mitchener, L. et al. BixBench: a Comprehensive Benchmark for LLM-based Agents in Computational Biology. (2025).

79. Lex, A., Gehlenborg, N., Strobelt, H., Vuillemot, R. & Pfister, H. UpSet: Visualization of Intersecting Sets. IEEE Trans Vis Comput Graph 20, 1983–1992 (2014).

80. Hao, M. et al. Large-scale foundation model on single-cell transcriptomics., Nat Methods 21, 1481–1491 (2024).

81. Yang, F. et al. scBERT as a large-scale pretrained deep language model for cell type annotation of single-cell RNA-seq data., Nat. Mach. Intell. 4, 852–866 (2022).

82. Yang, X. et al. GeneCompass: deciphering universal gene regulatory mechanisms with a knowledge-informed cross-species foundation model., Cell Res 34, 830–845 (2024).

83. Wen, H. et al. CellPLM: Pre-training of cell language model beyond single cells., bioRxiv (2023) doi:10.1101/2023.10.03.560734.

84. Tejada-Lapuerta, A. et al. Nicheformer: a foundation model for single-cell and spatial omics., Nat Methods 22, 2525–2538 (2025).

85. Chen, Y. et al. Toward mastering the cell language by learning to generate., bioRxiv (2024) doi:10.1101/2024.01.25.577152.

86. Shen, H. et al. Generative pretraining from large-scale transcriptomes for single-cell deciphering., iScience 26, 106536 (2023).

87. Zeng, Y. et al. CellFM: a large-scale foundation model pre-trained on transcriptomics of 100 million human cells., Nat Commun 16, 4679 (2025).

88. Yuan, X. et al. Cell-ontology guided transcriptome foundation model. (2024).

89. Kalfon, J., Samaran, J., Peyré, G. & Cantini, L. scPRINT: pre-training on 50 million cells allows robust gene network predictions. Nat Commun 16, 3607 (2025).

90. Ho, N. et al. Scaling dense representations for single cell with transcriptome-scale context., bioRxiv (2024) doi:10.1101/2024.11.28.625303.

91. Pang, K. et al. PULSAR: a Foundation Model for Multi-scale and Multi-cellular Biology., bioRxiv (2025) doi:10.1101/2025.11.24.685470.

92. LeRoy, N. J. et al. Atacformer: A transformer-based foundation model for analysis and interpretation of ATAC-seq data., bioRxiv (2025) doi:10.1101/2025.11.03.685753.

93. Cao, G., (曹广硕) et al. scPlantLLM: A Foundation Model for Exploring Single-cell Expression Atlases in Plants., Genomics Proteomics Bioinformatics 23, (2025).

94. Zhao, S., Zhang, J., Wu, Y., Luo, Y. & Nie, Z. LangCell: Language-Cell Pre-training for Cell Identity Understanding. (2024).

95. Levine, D. et al. Cell2Sentence: Teaching Large Language Models the Language of Biology., bioRxiv 2023.09.11.557287 (2024) doi:10.1101/2023.09.11.557287.

96. Chen, Y. & Zou, J. Simple and effective embedding model for single-cell biology built from ChatGPT. Nat Biomed Eng 9, 483–493 (2025).

97. Fang, Y. et al. ChatCell: Facilitating Single-Cell Analysis with Natural Language. (2024).

98. Anderson, D. J. et al. NKX2-5 regulates human cardiomyogenesis via a HEY2 dependent transcriptional network., Nature Communications 9, 1373 (2018).

99. Durocher, D., Chen, C. Y., Ardati, A., Schwartz, R. J. & Nemer, M. The atrial natriuretic factor promoter is a downstream target for Nkx-2.5 in the myocardium. Mol Cell Biol 16, 4648–4655 (1996).

100. Small, E. M. & Krieg, P. A. Transgenic analysis of the atrialnatriuretic factor (ANF) promoter: Nkx2-5 and GATA-4 binding sites are required for atrial specific expression of ANF. Dev Biol 261, 116–131 (2003).

101. Warren, S. A. et al. Differential role of Nkx2-5 in activation of the atrial natriuretic factor gene in the developing versus failing heart., Mol Cell Biol 31, 4633–4645 (2011).

102. Takimoto, E. et al. Up-regulation of natriuretic peptides in the ventricle of Csx/Nkx2-5 transgenic mice., Biochem Biophys Res Commun 270, 1074–1079 (2000).

103. Robbe, Z. L. et al. CHD4 is recruited by GATA4 and NKX2-5 to repress noncardiac gene programs in the developing heart., Genes Dev 36, 468–482 (2022).

104. Bruneau, B. G. et al. A murine model of Holt-Oram syndrome defines roles of the T-box transcription factor Tbx5 in cardiogenesis and disease., Cell 106, 709–721 (2001).

105. Hiroi, Y. et al. Tbx5 associates with Nkx2-5 and synergistically promotes cardiomyocyte differentiation., Nat Genet 28, 276–280 (2001).

106. Postma, A. V. et al. A gain-of-function TBX5 mutation is associated with atypical Holt-Oram syndrome and paroxysmal atrial fibrillation., Circ Res 102, 1433–1442 (2008).

107. Linhares, V. L. F. et al. Transcriptional regulation of the murine Connexin40 promoter by cardiac factors Nkx2-5, GATA4 and Tbx5., Cardiovasc Res 64, 402–411 (2004).

108. Pizard, A. et al. Connexin 40, a target of transcription factor Tbx5, patterns wrist, digits, and sternum., Mol. Cell. Biol. 25, 5073–5083 (2005).

109. Ching, Y.-H. et al. Mutation in myosin heavy chain 6 causes atrial septal defect., Nat Genet 37, 423–428 (2005).

110. Ghosh, T. K. et al. Physical interaction between TBX5 and MEF2C is required for early heart development., Mol Cell Biol 29, 2205–2218 (2009).

111. Kathiriya, I. S. et al. Modeling Human TBX5 Haploinsufficiency Predicts Regulatory Networks for Congenital Heart Disease., Dev Cell 56, 292–309.e9 (2021).

112. Hoogaars, W. M. H. et al. The transcriptional repressor Tbx3 delineates the developing central conduction system of the heart., Cardiovasc Res 62, 489–499 (2004).

113. Hoogaars, W. M. H. et al. Tbx3 controls the sinoatrial node gene program and imposes pacemaker function on the atria., Genes Dev 21, 1098–1112 (2007).

114. Hoogaars, W. M. H. et al. TBX3 and its splice variant TBX3 + exon 2a are functionally similar., Pigment Cell Melanoma Res 21, 379–387 (2008).

115. Bakker, M. L. et al. T-box transcription factor TBX3 reprogrammes mature cardiac myocytes into pacemaker-like cells., Cardiovasc Res 94, 439–449 (2012).

116. Wu, S.-P. et al. Atrial identity is determined by a COUP-TFII regulatory network., Dev Cell 25, 417–426 (2013).

117. Morin, S., Charron, F., Robitaille, L. & Nemer, M. GATA-dependent recruitment of MEF2 proteins to target promoters. EMBO J 19, 2046–2055 (2000).

118. Skerjanc, I. S., Petropoulos, H., Ridgeway, A. G. & Wilton, S. Myocyte enhancer factor 2C and Nkx2-5 up-regulate each other’s expression and initiate cardiomyogenesis in P19 cells. J Biol Chem 273, 34904–34910 (1998).

119. Vincentz, J. W., Barnes, R. M., Firulli, B. A., Conway, S. J. & Firulli, A. B. Cooperative interaction of Nkx2.5 and Mef2c transcription factors during heart development. Dev Dyn 237, 3809–3819 (2008).

120. Molkentin, J. D. & Markham, B. E. Myocyte-specific enhancer-binding factor (MEF-2) regulates alpha-cardiac myosin heavy chain gene expression in vitro and in vivo. J. Biol. Chem. 268, 19512–19520 (1993).

121. Meissner, J. D., Umeda, P. K., Chang, K.-C., Gros, G. & Scheibe, R. J. Activation of the beta myosin heavy chain promoter by MEF-2D, MyoD, p300, and the calcineurin/NFATc1 pathway. J Cell Physiol 211, 138–148 (2007).

122. Warkman, A. S. et al. Developmental expression and cardiac transcriptional regulation of Myh7b, a third myosin heavy chain in the vertebrate heart., Cytoskeleton (Hoboken) 69, 324–335 (2012).

